# Enabling Systemic Identification and Functionality Profiling for Cdc42 Homeostatic Modulators

**DOI:** 10.1101/2024.01.05.574351

**Authors:** Satyaveni Malasala, Fereshteh Azimian, Yan-Hua Chen, Jeffery L Twiss, Christi Boykin, Shayan Nik Akhtar, Qun Lu

## Abstract

Homeostatic modulation is pivotal in modern therapeutics. However, the discovery of bioactive materials to achieve this functionality is often random and unpredictive. Here, we enabled a systemic identification and functional classification of chemicals that elicit homeostatic modulation of signaling through Cdc42, a classical small GTPase of Ras superfamily. Rationally designed for high throughput screening, the capture of homeostatic modulators (HMs) along with molecular re-docking uncovered at least five functionally distinct classes of small molecules. This enabling led to partial agonists, hormetic agonists, *bona fide* activators and inhibitors, and ligand-enhanced agonists. Novel HMs exerted striking functionality in bradykinin-Cdc42 activation of actin remodelingand modified Alzheimer’s disease-like behavior in mouse model. This concurrent computer-aided and experimentally empowered HM profiling highlights a model path for predicting HM landscape.

**One Sentence Summary:** With concurrent experimental biochemical profiling and *in silico* computer-aided drug discovery (CADD) analysis, this study enabled a systemic identification and holistic classification of Cdc42 homeostatic modulators (HMs) and demonstrated the power of CADD to predict HM classes that can mimic the pharmacological functionality of interests.

**Introduction:** Maintainingbody homeostasisis the ultimate keyto health. Thereare rich resources of bioactive materials for this functionality from both natural and synthetic chemical repertories including partial agonists (PAs) and various allosteric modulators. These homeostatic modulators (HMs) play a unique role in modern therapeutics for human diseases such as mental disorders and drug addiction. Buspirone, for example, acts as a PA for serotonin 5-HT_1A_ receptor but is an antagonist of the dopamine D_2_ receptor. Such medical useto treat general anxietydisorders (GADs) has become one of the most-commonly prescribed medications. However, most HMs in current uses target membrane proteins and are often derived from random discoveries. HMs as therapeutics targeting cytoplasmic proteins are even more rare despite that they are in paramount needs (e. g. targeting Ras superfamily small GTPases).

**Rationale:** Cdc42, a classical member of small GTPases of Ras superfamily, regulates PI3K-AKT and Raf-MEK-ERK pathways and has been implicated in various neuropsychiatric and mental disorders as well as addictive diseases and cancer. We previously reported the high-throughput *in-silico* screening followed by biological characterization of novel small molecule modulators (SMMs) of Cdc42-intersectin (ITSN) protein-protein interactions (PPIs). Based on a serendipitously discovered SMM ZCL278 with PA profile as a model compound, we hypothesized that there are more varieties of such HMs of Cdc42 signaling, and the model HMs can be defined by their distinct Cdc42-ITSN binding mechanisms using computer-aided drug discovery (CADD) analysis. We further reasoned that molecular modeling coupled with experimental profiling can predict HM spectrum and thus open the door for the holistic identification and classification of multifunctional cytoplasmic target-dependent HMs as therapeutics.

**Results:** The originally discovered Cdc42 inhibitor ZCL278 displaying PA properties prompted the inquiry whether this finding represented a random encounter of PAs or whether biologically significant PAs can be widely present. The top ranked compounds were initially defined by structural fitness and binding scores to Cdc42. Because higher binding scores do not necessarily translate to higher functionality, we performed exhaustive experimentations with over 2,500 independent Cdc42-GEF (guanine nucleotide exchange factor) assays to profile the GTP loading activities on all 44 top ranked compounds derived from the SMM library. The N-MAR-GTP fluorophore-based Cdc42-GEF assay platform provided the first glimpse of the breadth of HMs. A spectrum of Cdc42 HMs was uncovered that can be categorized into five functionally distinct classes: Class I-partial competitive agonists, Class II-hormetic agonists, Class III-*bona fide* inhibitors (or inverse agonists), Class IV-*bona fide* activators or agonists, and Class V-ligand-enhanced agonists. Remarkably, model HMs such as ZCL278, ZCL279, and ZCL367 elicited striking biological functionality in bradykinin-Cdc42 activation of actin remodeling and modified Alzheimer’s disease (AD)-like behavior in mouse model.

Concurrently, we applied Schrödinger-enabled analyses to perform CADD predicted classification of Cdc42 HMs. We modified the classic molecular docking to instill a *preferential binding pocket order* (PBPO) of Cdc42-ITSN, which was based on the five binding pockets in interface of Cdc42-ITSN. We additionally applied a structure-based pharmacophore hypothesis generation for the model compounds. Then, using Schrödinger’s Phase Shape, 3D ligand alignments assigned HMs to Class I, II, III, IV, and V compounds. In this HM library compounds, PBPO, matching pharmacophoric featuring, and shape alignment, all put ZCL993 in Class II compound category, which was confirmed in the Cdc42-GEF assay.

**Conclusion:** HMs can target diseased cells or tissues while minimizing impacts on tissues that are unaffected. Using Cdc42 HM model compounds as a steppingstone, GTPase activation-based screening of SMM library uncovered five functionally distinct Cdc42 HM classes among which novel efficacies towards alleviating dysregulated AD-like features in mice were identified. Furthermore, molecular re-docking of HM model compounds led to the concept of PBPO. The CADD analysis with PBPO revealed similar profile in a color-coded spectrum to these five distinct classes of Cdc42 HMs identified by biochemical functionality-based screening. The current study enabled a systemic identification and holistic classification of Cdc42 HMs and demonstrated the power of CADD to predict an HM category that can mimic the pharmacological functionality of interests. With artificial intelligence/machine learning (AI/ML) on the horizon to mirror experimental pharmacological discovery like AlphaFold for protein structure prediction, our study highlights a model path to actively capture and profile HMs in potentially any PPI landscape.

**Graphic Abstract:** 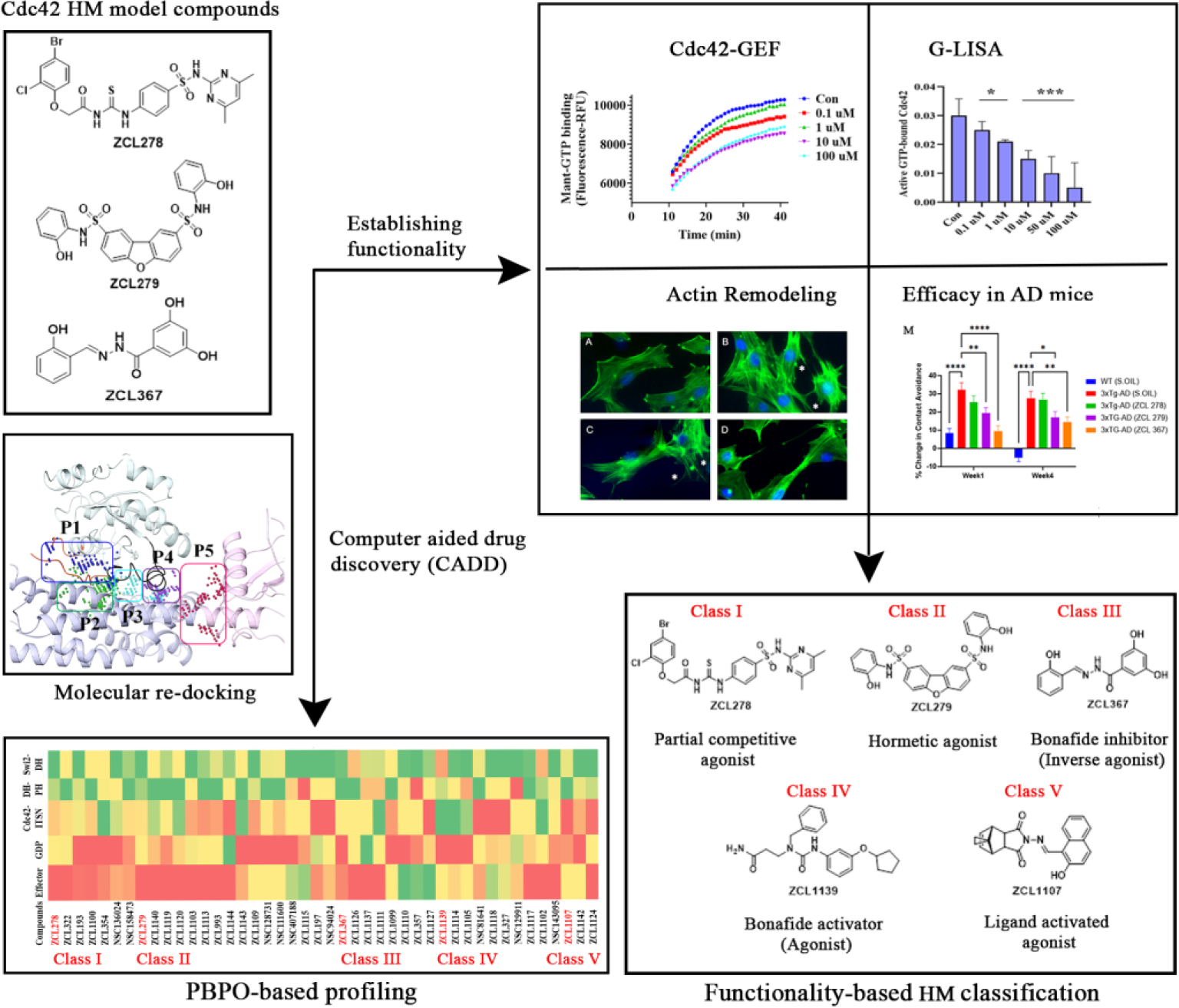

**Identification and functional classification of Cdc42 homeostatic modulators (HMs):** Using Cdc42 HM model compounds as reference, GTPase activation-based screening of compound libraries uncovered five functionally distinct Cdc42 HM classes. HMs showed novel efficacies towards alleviating dysregulated Alzheimer’s disease (AD)-like behavioral and molecular deficits. In parallel, molecular re-docking of HM model compounds established their preferential binding pocket orders (PBPO). PBPO-based profiling (Red reflects the most, whereas green reflects the least, preferable binding pocket) revealed trends of similar pattern to the five classes from the functionality-based classification.

---

Maintaining body homeostasis is the ultimate key to health. There are rich resources of bioactive materials from both natural and syntheticchemical repertoriesfor this functionality, such as partial agonists (PAs) and various allosteric modulators (*1*). The characteristic features of PAs are that they often can activate receptors to give a desired response when inadequate amounts of the endogenous ligand are present, or they can lessen the overstimulation of receptors when excess amounts of the endogenous ligands are produced in the body pathologically (*2*). PAs often induce less desensitization and can be powerful modulators to help restore body’s homeostasis (*3*).

PAs and their alike play a unique role in modern therapeutics for management of human diseases such as neuropsychiatric disorders and drug addiction. Buspirone, for example, acts as a PA for serotonin 5-HT_1A_ receptor with high affinity, but is an antagonist of dopamine D_2_ receptor (*4*). Such medical use to treat general anxiety disorders (GAD) has become one of the most-commonly prescribed medications (*5*).

However, most homeostatic modulators (HMs) such as PAs in current uses target membrane proteins suchas plasma andnuclear membrane receptors, transporters, and ion channels (*6*). PAs for targeting cytoplasmic proteins are rare and often unplanned discoveries. Nevertheless, therapeutics targeting cytosolic enzymes are in paramount needs, like the recent FDA approved Lumakras in treating K-Ras mutant linked lung cancer (*7*).

Ras superfamily proteins are small GTPases that act as molecular switches and regulate a myriad of essential cellular functions in proliferation, vesicular transport, cell motility, and others (*8, 9*). Cdc42, a classical member of the Rho GTPase subfamily, is an important regulator of many biological processes (*10–12*). Cdc42 function is dependent on its GDP and GTP bound state. The activity of Cdc42 is controlled by GTPase activating proteins (GAPs) that promote the hydrolysis of GTP, guanine nucleotide exchange factors (GEFs) that stimulate therelease of GDPand binding of GTP, and guanine nucleotide dissociation inhibitors (GDIs) that prevent the nucleotide exchange (*13, 14*). When Cdc42 bound to GTP, it interacts with over 20 downstream effectors involved in a variety of cellular signals, such as through PI3K-AKT and Raf-MEK-ERK pathways (*15, 16*). Cdc42 signaling has been implicated as an important target in various human diseases and is being actively pursued pharmacologically (*11, 17-24*).

We previously reported the discovery and characterization of novel small molecule modulators (SMMs) of Cdc42-intersectin (ITSN) interaction by using computer-assisted high throughput *in silico* screening (*25*). Based on a serendipitously discovered SMM ZCL278 with PA profile (*26*), we hypothesized that there may be more varieties of such HMs of Cdc42 signaling in the existing SMM libraries, and the model HMs such as ZCL278 can be defined by their distinct Cdc42-ITSN binding mechanisms using computer-aided drug discovery (CADD) analysis. We further reasoned that molecular modeling coupled with experimental profiling can predict HM spectrum and thus open the door for the holistic identification and classification of multifunctional cytoplasmic target-dependent HMs as therapeutics. We thus sought to enable a systemic capture of SMM repertoires with Cdc42 PA signaturesmodulating Cdc42-ITSNinteractionsand signaling. We investigated molecular modeling to structural-function prediction for profiling HMs that are previously hard to capture.

## Enabling systemic identification of HMs on Cdc42-ITSN signaling

Computer aided high throughput screening of chemical compound library has identified the first Cdc42 selective inhibitor ZCL278 that dislodges the interactions between Cdc42 and its GEF, ITSN (*25*). ZCL278 regulates Cdc42 mediated filopodia and actin cytoskeletal remodeling (*25*). As an important element in the epidermal growth factor (EGF) and Ras oncogene signaling, Cdc42 promotes cancer cell proliferation and migration, which can be suppressed by ZCL278 and another SMM ZCL367 targeting the same pharmacophore (*26*). However, in a serendipitous finding, we observed that ZCL278 can also activate Cdc42 under certain conditions and appeared to display PA features (*26*).

To determine whether this finding represented a random encounter of PA or whether biologically significant PAs can be widely captured, we rescreened the same chemical libraries for compounds that display the same or similar PA features employing an N-MAR-GTP fluorophore-based Cdc42-GEF assay platform. Supporting the previous studies, ZCL278 promoted GTP loading in the absence of GEF, but inhibited GTP loading in the presence of GEF (**Fig. 1A** and **B**). While ZCL367 inhibited Cdc42 GTP loading with or without the presence of an activating GEF (**Fig. 1E** and **F**), some compounds display features that go beyond the traditional categories of Cdc42 inhibitors or activators. ZCL279, for example, showed that whether in the absence or presence of GEF, it promoted GTP loading to Cdc42 at the concertation up to low micromoles (10 µM), but inhibited GTP loading at higher concentrations (**Fig. 1C** and **D**). This feature of ZCL279 is reminiscent of previously described hormetic chemicals (*27*). Using ITSN as reference of endogenous full agonist, ZCL278 and ZCL279 displayed PA features whereas ZCL367 is a *bona fide* Cdc42 inhibitor or inverse agonist (IA) (**Fig. 1G, H**, and **I**). The GTPase-Glo™ assay, which quantitates the amount of GTP remaining after a GTPase reaction, provided the independent support for our discovery (**Fig. 1J** and **K**).

**Figure 1.**
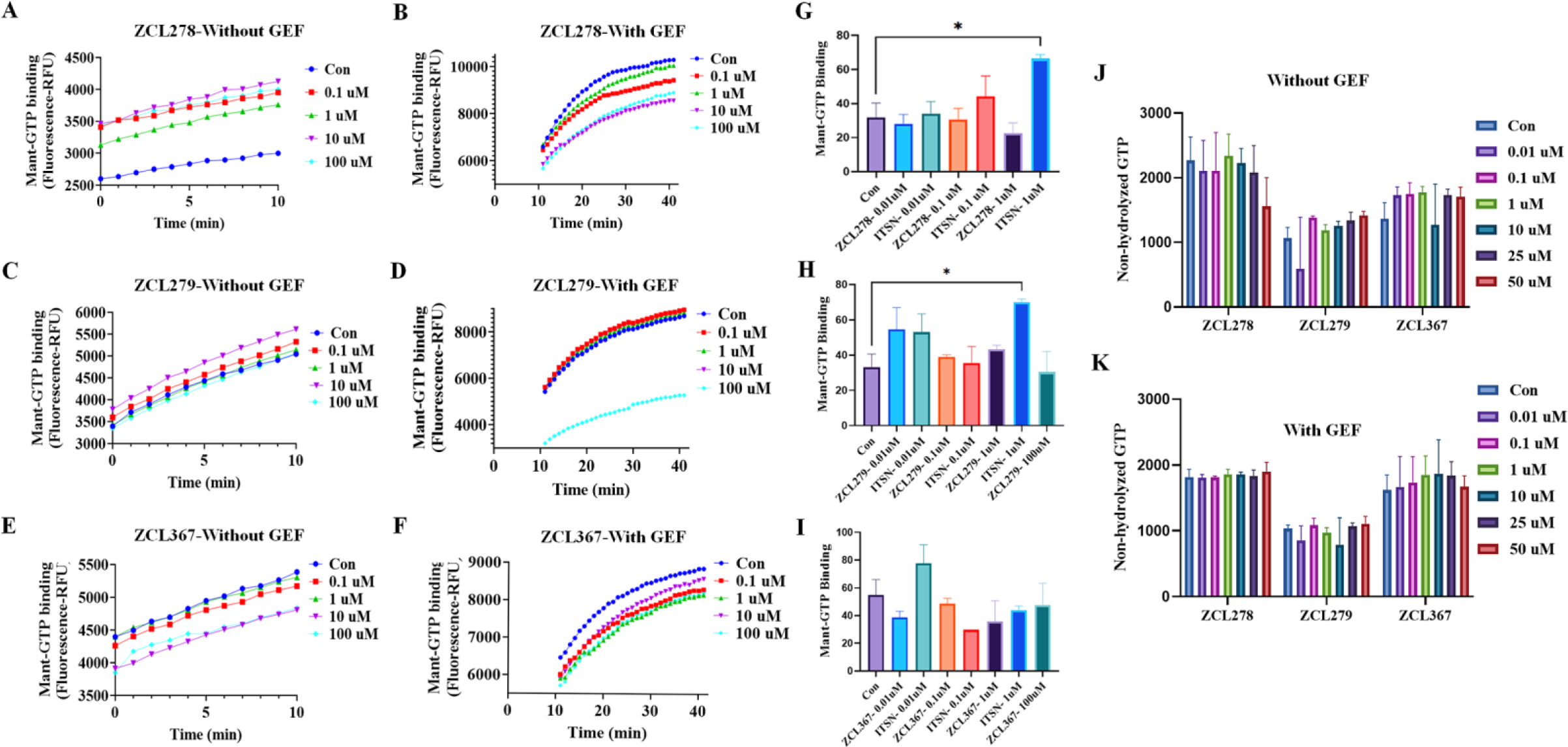
Biochemical functionality of agonistic, hormetic and antagonistic effects of HM model compounds *in vitro*. Cdc42 HM model compounds regulate Cdc42 GTPloadingand its GEF-mediated activation. ZCL278 and ZCL279 (at low concentrations) increase Cdc42 GTP loading while ZCL367 decreases Cdc42 GTP loading. **(A)** ZCL278, **(C)** ZCL279 and **(E)** ZCL367 without GEF. ZCL278, ZCL279 (at high concentrations) and ZCL367 decrease GEF-mediated Cdc42 GTP loading. **(B)** ZCL278, **(D)** ZCL279 and **(F)** ZCL367 with GEF. HM model compounds act through GEF like mechanism. **(G-I)** The amount of GTP loading on Cdc42 for ITSN as a full agonist were comparable to those for HMs in different concentrations (0.01-10 µM). **(G)** ZCL278, **(H)** ZCL279 and **(I)** ZCL367**. (J-K)** HM model compounds have no effect on GTP hydrolyzation with and without ITSN. Treatment of Cdc42 by different concentrations (0.01-50 µM) of ZCL278, ZCL279 and ZCL367 without and with GEF do not have effects on hydrolyzation of GTP significantly compared to control. Control for all samples without ITSN was DMSO and Cdc42, for all samples with GEF was DMSO, Cdc42 and ITSN. All data are presented as mean±SEM from duplicates or triplicates from at least two independent experiments. ANOVA compared treatments to their respective control (* p < 0.05, ** p < 0.01, *** p < 0.001, # p < 0.0001). Note: The Guanine nucleotide exchange (GEF) reaction was performed using the relative fluorescence intensity of N-MAR-GTP exchange that monitors GDP release during the exchange reaction after treatment by different concentrations (0.01-100 µM) of ZCL278, ZCL279 and ZCL367 on Cdc42 in absence or presence of ITSN. The GTPase-GLO assay was conducted to measure unhydrolyzed GTP levels in the presence of HM model compounds.

## Structural simulation of binding mechanisms of different Cdc42-ITSN HM classes

These findings raised the intriguing possibility that there may be more varieties of PAs and IAs in the existing SMM libraries, and these three model classes of Cdc42 modulators can possibly be defined by their distinct binding mechanisms using computer-aided drug discovery (CADD) analysis.

We employed Site MAP of Schrödinger software to identify the top ranked potential protein binding pockets at the interface of Cdc42-ITSN (PDB code: 3QBV). Specific residues and site scores for each binding pocket are summarized (See **Supplementary Table S1, Fig. S1** and **S2**). For Cdc42 not complexed with ITSN, three different binding pockets were detected, which included the GDPbindingsite on Cdc42, a region adjacent to the GDPbindingsite, and thespecific residues of the structural motifs “Switch I” (Val36, Phe37) and “Switch II” (Ala59, Tyr64, Leu67, Leu70, Ser71). We named these detected pockets as “GDP”, “Effector”, and “Switch 1&2” pockets, respectively (**Fig. 2A**). For Cdc42-ITSN complex, besides GDP and Effector pockets, additional three binding pockets were identified including “Cdc42-DH” domain, “DH-PH” domain of ITSN, and “Switch 2 of Cdc42 and DH” domain of ITSN (**Fig. 2C**).

**Figure 2.**
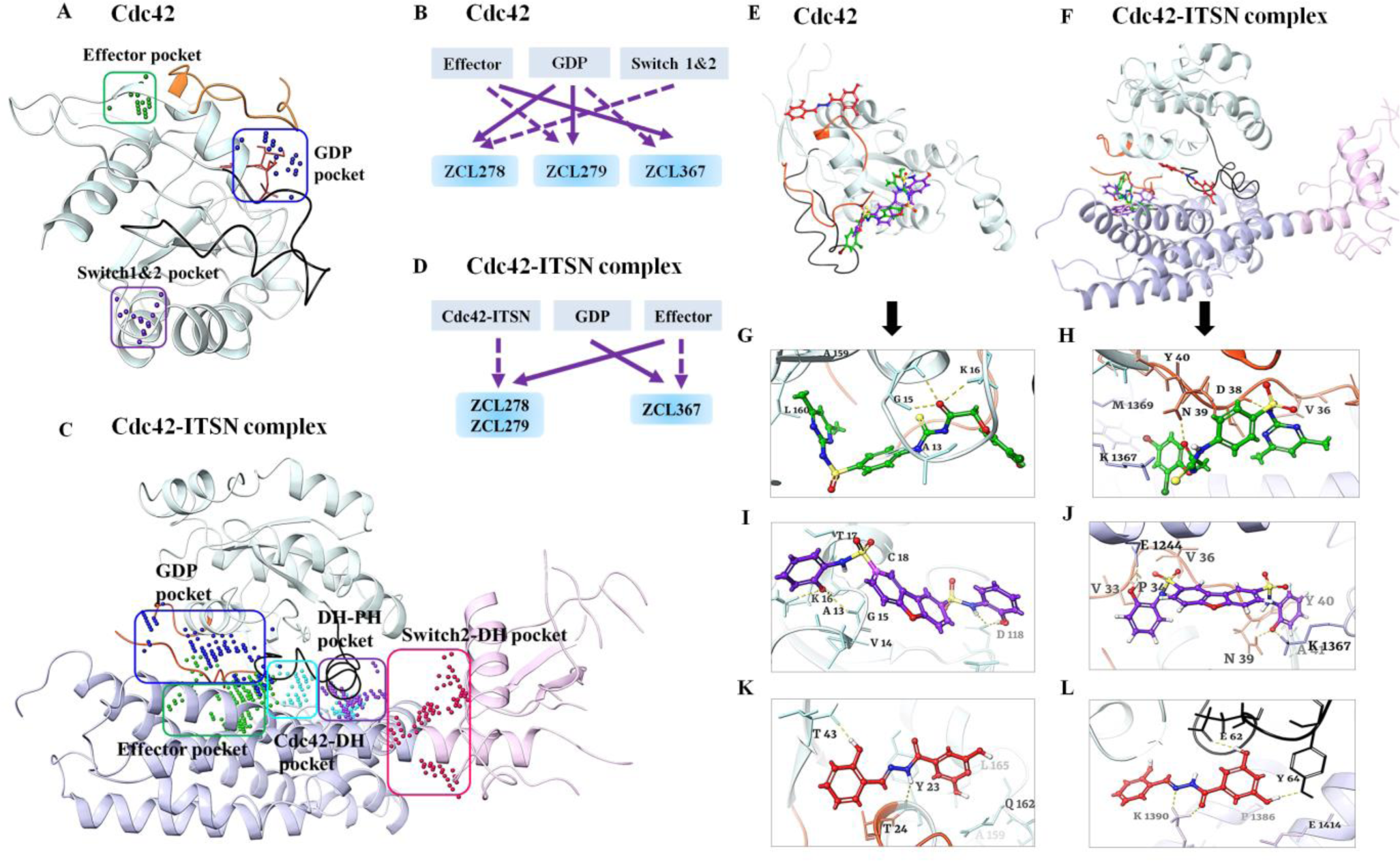
Binding modes of Cdc42 HM model compounds. Different predicted binding pockets for Cdc42 HM model compounds. **(A** and **C)** 3D structural illustration of different drug-binding pocket of Cdc42 (transparent cyan, switch 1 (a.a.24-40) region is in orange color & switch 2 (a.a.57-76) region in black color) and Cdc42-ITSN complex (DH and PH domain of ITSN are shown purple and pink ribbons) (PDB: 3QBV). The GDP, effector and switch1&2 binding pockets for Cdc42 and fivebindingpockets including GDP, Effec tor, Cdc42-DH domain of ITSN, DH-PH domain of ITSN and switch2 of Cdc42 and DH domain of ITSN were detected. (Effector and GDP pockets were the same for Cdc42 and Cdc42-ITSN). Cdc42 model compounds bind preferably in different binding pockets. **(B** and **D)** Schematic diagrams of preferential binding pockets for ZCL278, ZCL279 and ZCL367 for Cdc42 and Cdc42-ITSN complex. First and second preferential binding pockets are shown by solid and dotted lines respectively. ZCL278 and ZCL279 bind preferably to GDP pocket while ZCL367 preferentially binds Effector pocket of Cdc42. The second preferential pockets of Cdc42 for ZCL278, ZCL279 and ZCL367 are Switch1&2, Effectorand GDPpockets respectively. In the presence of ITSN, for ZCL278 and ZCL279, first preferential binding pocket of Cdc42-ITSN complex is Effector and the second preferential pocket is Cdc42-DH domain of ITSN pocket while ZCL367 binds to GDP pocket and effector pocket preferably. (**E-F**) Superimpose of ZCL278 (green), ZCL279 (purple) and ZCL367 (red) in Cdc42 and Cdc42-ITSN complex. **(G-L)** Cdc42 HM model compounds show different poses with a certain extent of overlap with each other. **(G** and **I)** ZCL278 (green) and ZCL279 (purple) bind to GDP pocket of CdC42 *via* two hydrogen bonds with Lys16, GLy15, as well as hydrophobic interactions with Val14, Ala13 and Val33 of Cdc42. **(K)** ZCL367 has shown hydrogen bonds with Tyr23 and Thr43, as well as hydrophobic interactions with Val42 and Leu165 in the effector binding pocket of Cdc42 (PDB: 3QBV). **(H** and **J)** In the presence of ITSN, ZCL278 (green) and ZCL279 (purple) interact with effector pocket of CdC42-ITSN *via* hydrogen bonds with Asn39 and Lys1367, as well as hydrophobic interactions with Pro34 and Val36. **(L)** ZCL367 formed hydrogen bonds with Asp63, Glu62 and Tyr64, as well as hydrophobic interactions with Ala13 and Val33 in the GDP binding pocket of Cdc42-ITSN.

We then re-docked ZCL278, ZCL279 and ZCL367 to all identified pockets of the Cdc42 and Cdc42-ITSN complex to investigate the binding modes of these model compounds. The first and secondpreferential bindingsites of Cdc42 for these three compounds are shown in **Figure 2B**. ZCL278 and ZCL279 bind most preferably to GDP pocket while ZCL367 preferentially binds Effector pocket. The second preferential pockets of Cdc42 for ZCL279, ZCL278 and ZCL367 are Effector, Switch 1&2, and GDP pockets, respectively (**Fig. 2B**). We designated the binding modes as the *preferential binding pocket order* (PBPO). In the presence of ITSN, the PBPO for these compounds were different. For ZCL278 and ZCL279, the first preferential binding pocket on Cdc42-ITSN complex is Effector, and the second preferential binding pocket is Cdc42-DH domain of ITSN, while ZCL367 binds to GDP pocket preferably followed by Effector pocket (**Fig. 2D**). **Figures 2E** and **F** showed superimposed images of ZCL278, ZCL279, and ZCL367 on Cdc42 and Cdc42-ITSN complex, respectively. These results revealed that the binding modes of ZCL278 and ZCL279 are alike while ZCL367 interacts with different binding pockets of Cdc42 and Cdc42-ITSN complex from ZCL278 and ZCL279.

The molecular interactions of these model compounds with Cdc42 in the absence and presence of ITSN are further revealed in **Figures 2G-L (See also Supplementary Fig. S1-2).** These compounds made interactions with a variety of Cdc42 and Cdc42-ITSN residues via hydrogen bond, halogenic and hydrophobic binding **(Supplementary Table S2).** The docking result showed that the oxygen atom of hydroxyl group and nitrogen amide group of ZCL367 formed hydrogen bonds with Thr43 and Tyr23 of Cdc42 (**Fig. 2K**) while in the presence of ITSN, two oxygen atoms of ortho of hydroxyl group made interaction with the Tyr64, Glu62 and Asp63 *via* hydrogen bonds. Furthermore, Lys1390 of ITSN formed three centered hydrogen bonds with nitrogen atom and oxygen carbonyl of ZCL367 (**Fig. 2L**). Additionally, carbonyl oxygen of ZCL278 interacted with Gly15 and Lys16 of Cdc42 through hydrogen interactions (**Fig. 2G**). In the presence of ITSN, Asp38 and Asn39 of Cdc42 as well as Lys 1367 of ITSN formed hydrogen bonds with oxygen atom of carbonyl, amide nitrogen atoms of sulfonamides and thiourea moiety of ZCL278, respectively (**Fig. 2H**). As shown in **Figure 2I**, the nitrogen atom of sulfonamide moiety and oxygen atom hydroxyl group of ZCL279 formed an hydrogen bond with Asp118, Gly15 and Lys16 from Cdc42 (**Fig. 2I**). In the presence of ITSN, oxygen atom of hydroxyl group of ZCL279 formed hydrogen bond with Glu1244 of ITSN. The other three centered H-bonds were formed between oxygen group of ZCL279 and Asn39 of Cdc42 and Lys1367 of ITSN (**Fig. 2J**). ZCL278 and ZCL279 showed similar interactions with GDP and Effector pockets of Cdc42-ITSN complex. It seems that in the presence of ITSN, these compounds could make more interactions with Cdc42 compared to that in the absence of ITSN as a GEF. Dockingscoresof these compounds also confirmed that, in the presence of ITSN, compounds bind to Cdc42 with a higher score compared to those for Cdc42 without ITSN (**Supplementary Table S2**). Compared to ZCL278, ZCL279 showed stable interactions with both Cdc42 and Cdc42-ITSN. For ZCL279, the ligand-protein interactions are favorable and strong, confirmed and supported by molecular dynamics (MD) studies (See Supplementary **Fig. S3**-**5**). On the other hand, for ZCL278, it does show interactions although they are more moderate. Protein-ligand interactions between Cdc42 and ZCL367 are similarly stable and strong regardless of whether GEF was present or not (See Supplementary Fig. S3-5).

## Homeostatic modulating effects on Cdc42 activation in cells

We then investigated the model compounds that represent the distinct classes (ZCL278, ZCL279, and ZCL367) in a cell-based assay to determinewhether their different interactionmodes can be manifested in their differential ability to modulate Cdc42 activation in biological systems. **Figure 3** shows that, with a G-LISA platform that measures the GTP bound form of Cdc42 from lysates of Swiss 3T3 cells, ZCL278 promoted Cdc42 activation (**Fig. 3A**). On the other hand, ZCL279 promoted Cdc42 activation at the concertation up to low micromoles (10 µM) but inhibited theactivationat higher concentrations (**Fig. 3B**). Under the same experimental condition, ZCL367 suppressed Cdc42 activation in a straight concentration-dependent manner (**Fig. 3C**). These findings are in general agreement with the *in vitro* GEF assays on the three model compounds.

**Figure 3.**
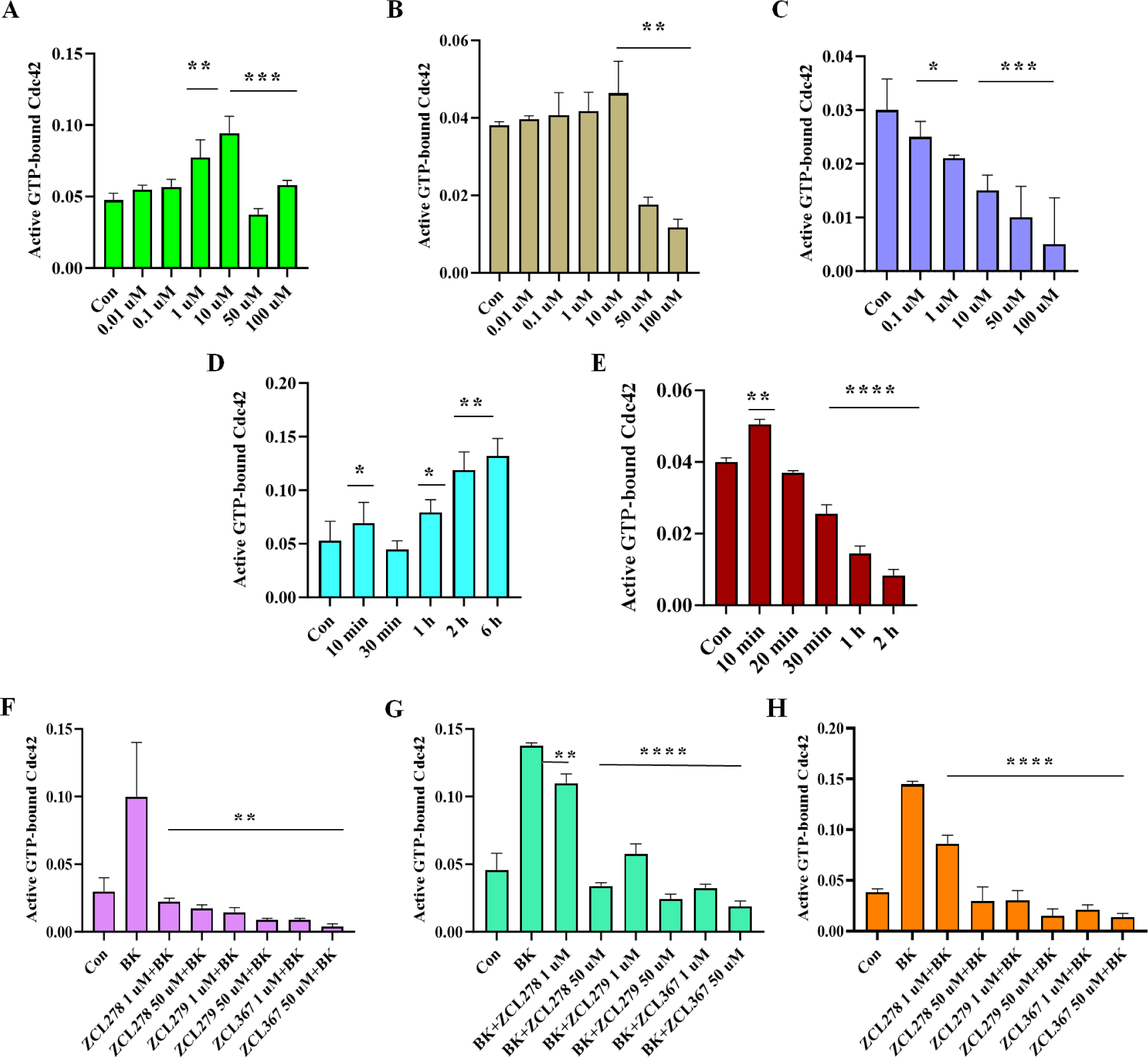
Cell-based functionality of Cdc42 activation by Cdc42 HM model compounds. Cdc42 activation measured in a G-LISA assay. Swiss 3T3 cells were serum starved overnight and were incubated for 15 min with different concentrations (0.01, 0.1, 1, 10, 50 and100 uM) of HM model compounds. **(A**) ZCL278, (**B**) ZCL279, and (**C**) ZCL367. ZCL278 and ZCL279 (at the concertation up to low micromoles (10 uM)) promoted Cdc42 activation while ZCL367 inhibited the Cdc42 activation; Cdc42 HM model compounds modulate Cdc42 activation in a time-dependent manner. Serum-starved Swiss 3T3 cells were incubated at different time intervals (10, 20, 30, 60, 120 and 360 min) with 50 uM of (**D**) ZCL278 and (**E)** ZCL279. Cdc42 HM model compounds regulate activation of Cdc42 by bradykinin (BK). Serum-starved Swiss 3T3 cells were incubated with 1 and 50 µM of ZCL278, ZCL279 and ZCL367 and then stimulated with Cdc42 activator bradykinin for 20 min. Incubation time were **(F)** 1h, (**G)** 15 min and **(H)** 6hrs. A constitutively active Cdc42 was used as positive control. Negative controls included vehicle-only controls as well as untreated cellular lysates. ZCL278 at low concentrations led to a higher GTP bounded-Cdc42 level while ZCL279 and ZCL367 led to a decreased level of GTP-bounded Cdc42 compared to control. All data are presented as mean±SEM from duplicates from three independent experiments. ANOVA compared treatments to their respective control (* p < 0.04, ** p < 0.004, *** p < 0.0004, **** p < 0.000 1).

However, it is interesting to note that with the 15-minute treatment durations as shown above (**Fig.3A**), 50 and 100 µM ZCL278 appeared to display a setback in activation pattern in that they displayed antagonistic effects although not quite as potent as the inhibitory ZCL367 (Compare **Fig. 3A** and **3C**). Therefore, we set to determine whether the differential effects of ZCL278, ZCL279, and ZCL367 on Cdc42 activation are time dependent. **Figure 3D** shows that Cdc42 activation by ZCL278 at 50 µM formed an initial peak at 10 minutes. ZCL278 then displayed an antagonistic effect at 30 minutes followed by resuming a strong agonistic effect up to 6 hours without showing the plateau. As a comparison, ZCL279 at 50 µM, although showing moderate activating effects at 10 minutes, suppressed Cdc42 activation thereafter in a time-dependent manner (**Fig. 3E**). Therefore, in general, ZCL278 promoted Cdc42 activation whereas ZCL279 manifested a biphasic and hormetic effect on Cdc42 activation in cell-based assay, consistent with the *in vitro* findings.

Bradykinin (BK) is widely known to activate Cdc42 signaling in a variety of cells (*28–30*) including Swiss 3T3 cells. Using G-LISA assays, GTP bound, activate Cdc42 levels in lysates of serum-starved Swiss 3T3 cells treated with ZCL278, ZCL279 and ZCL367 at 1 and 50 µM in different time points were quantified *via* PAK binding. **Figure 3G** and **H** showed that when Cdc42 is prior activated by BK, ZCL278 at low concentrations sustained a higher GTP bound Cdc42 level after treatment for 15 minutes and 6 hours when compared to control (vehicle) but antagonized the BK stimulation. On the other hand, ZCL367 treatments for 15 minutes and 6 hours at 1 and 50 µM all led to a decreased level of GTP bound Cdc42 compared to control. However, at high concentrations ZCL278 and ZCL279 acted like ZCL367 that inhibited Cdc42 activation to below control level (**Fig. 3F-H**). This is surprising as GEF assays *in vitro* and G-LISA in cells all demonstrated that ZCL279 promoted Cdc42 activation at low micromolar concentrations.

## HM effects of model compounds on actin remodeling in cells

Next, we explored the phenotypic effects of these three model compounds on Swiss 3T3 cells because BK induced a robust actin remodeling due to Cdc42 activation (*25, 26, 28*). Serum-starved Swiss 3T3 cells showed distinct focal adhesions and ridges around cell edges (**Fig. 4A: Arrows and Arrowhead**). BK stimulation led to a loss of focal adhesion while the smooth cell edges gave way to microspikes or filopodia (**Fig. 4B: Asterisks**). ZCL278 at 50 µM promoted microspikes or filopodia much like BK (**Fig. 4C: Asterisks**). When the cells were treated with BK, ZCL278 antagonized the BK effects on the induction of microspikes or filopodia (**Fig. 4D**). ZCL279 at 1-10 µM did stimulate the formation of microspikes and filopodia (Data not shown), although at 50 µM, it showed moderate formation of microspikes or filopodia with additional morphological alterations **(Fig. 4E: Asterisks**). This morphological anomaly was exacerbated when the cells were treated with BK and ZCL279 (**Fig. 4F**). Therefore, consistent with the findings from both GEF and G-LISA examinations, the effects of ZCL279 were hormetic. ZCL367 suppressed microspikes and filopodia and recovered ridges and focal adhesions in both BK stimulated or unstimulated cells (**Fig. 4G** and **4H: Arrows and Arrowheads**).

**Figure 4.**
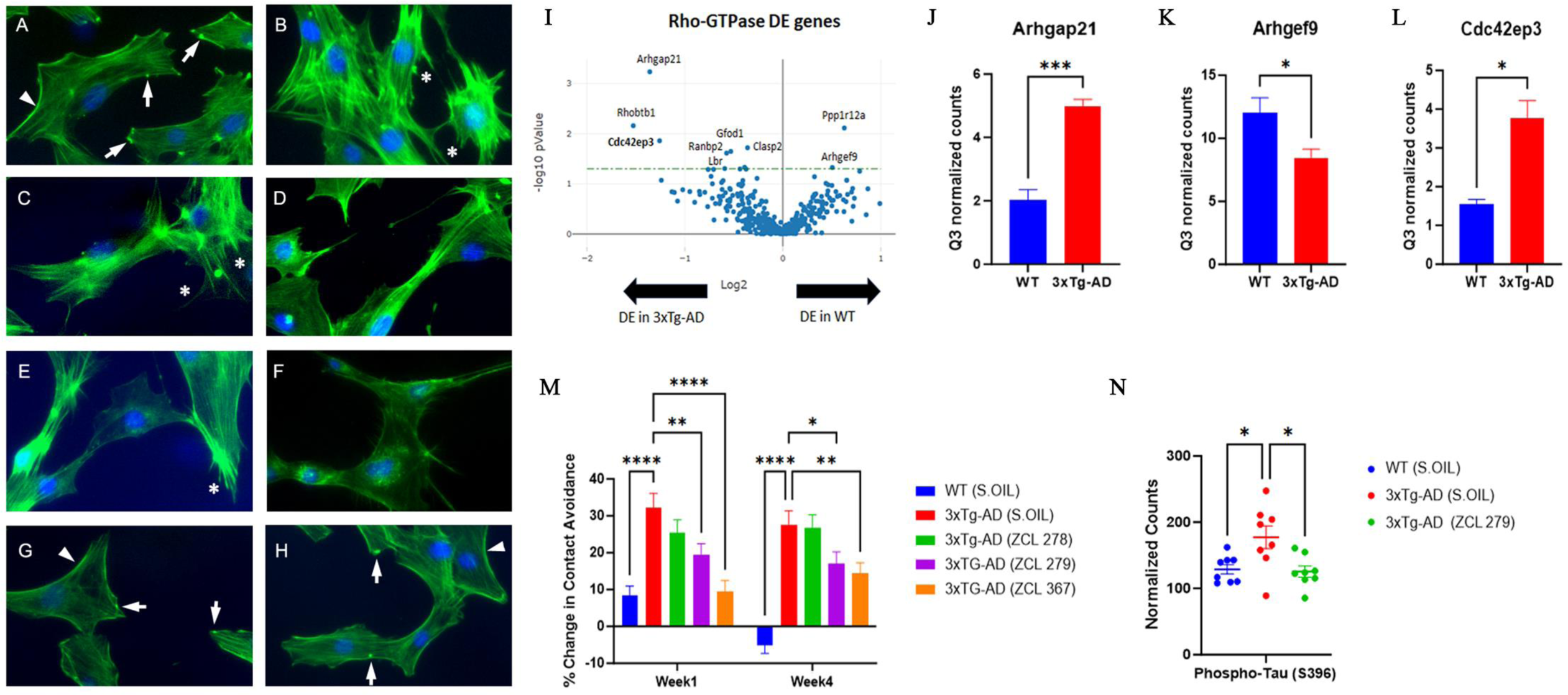
Biological functionality modulated by Cdc42 HM model compounds. Cdc42-based actin remodeling in cultured cells. Serum starved Swiss 3T3 cells were treated with HM model compounds and DMSO was used as a vehicle control. Cells were fixed and stained with rhodamine–phalloidin to label filamentous actin (Red). Cell nuclei were stained with Hoechst (Blue). Arrows point to the focal adhesion whereas arrowheads depict cell peripheral ridges. Asterisks demonstrate filopodia or microspikes. Bar: 20 µm. **(A)** Vehicle, **(B)** BK, **(C)** ZCL278, **(D)** ZCL278+BK, **(E)** ZCL279, **(F)** ZCL279 +BK, **(G)** ZCL367 and **(H)** ZCL367+BK. **(I)** Volcano plot demonstratingdifferentiallyexpressed (DE) genes in the RhoA, Rac1, and Cdc42 GTPase signalingnetwork in WTand 3xTg-AD mice. **(J-L)** Bar graph showing examples of Cdc42 signaling-associatedgenes that demonstratedsignificant differences in expression in 3xTg-AD compared to WT. n=2 mice per group. Data were presented as mean ± SEM * P ≤ 0.05, ** P ≤ 0.01, *** P ≤ 0.001. **(J)** Arhgap21: Rho GTPase Activating Protein 21, **(K)** Arhgef9: Cdc42 Guanine Nucleotide Exchange Factor 9, **(L)** Cdc42ep3: Cdc42 effector protein 3. **(M)** Dysfunctional 3xTg-AD mouse social contact behavior attenuated by HM model compounds. n= 15 mice per group* P ≤ 0.05, ** P ≤ 0.01, *** P ≤ 0.001, **** P ≤ 0.0001. **(N)** Scatter plot showing examples of Cdc42 HM model compound treatment that reduced phospho-Tau (S396) expression in 3xTg-AD mice compared to 3xTg-AD (Sesame oil). n= 2 mice per group with 4 brain regions analyzed per mouse. * P < 0.05, ** P <0.01, *** P <0.001, **** P < 0.0001.

## HMs modified interactive behavior in AD mice *in vivo*

Cdc42-ITSN dysregulation has long been suspected to contribute to neurogenerative diseases such as Alzheimer’s diseases (AD) and related dementias (ADRD) (*31*). We recently discovered that small GTPase signaling such as RhoA was spatially dysregulated in the brain of triple transgenic mouse model (3xTg-AD) bearing AD-like mutations in presenilin-1 (PS1M146V), amyloid precursor protein(APPsw), and microtubule-associated protein tau (Tau301P) (*32*). When these mice were examined by unbiased RNA transcriptomic analyses using GeoMx, changes in Cdc42 signaling modulatory elements including downregulation of *Argef9* mRNA and upregulation of *Argap21* mRNA were seen in the 3xTg-AD mouse brain (**Fig. 4I, 4J** and **4K**). The dysregulation of the major Cdc42 effector in layer 3 and 4 cortical neurons in brain, *cdc42ep3*, was reported in psychiatric disorders such as schizophrenia (*33*). We found that *cdc42ep3* was upregulated in the 3xTg-AD mouse brain (**Fig. 4L**). We then examined the social contact behavior of 3xTg-AD mice. Under the condition whentheywere left unapproached, 3xTg-AD mice showed reduced spontaneous activity (*34*). However, when they were approached and lifted by an experimenter, they quickly ran away (**Fig. 4M**). All three model compounds, to varying degrees, attenuated this contact avoidance behavior of 3xTg-AD mice (**Fig. 4M**). AD associated protein phospho-tau (S396) levelswere reducedin 3xTg-AD mice upon treatment with ZCL279 (**Fig. 4N**). These effects were consistent with the ADME (Absorption, distribution, metabolism, and excretion) and pharmacokinetics studies that these compounds passed blood brain barrier (*35*), demonstrating their potential functionality in modulating AD-like neuropathogenesis.

## Experimental profiling of HMs

Since these three model classes of Cdc42 modulators demonstrated significant functionality in BK-Cdc42 activation of actin remodeling and modified AD-like behavior in a mouse model, we questioned whether there are more compounds in our top ranked SMMs which may display the same or similar properties. The top ranked compounds were initially obtained by high throughput virtual screening based on structural fitness and binding scores (*25*). Because higher bindingscoresdo not necessarily translate to higher functionality, we performed exhaustive analyses with over 2,500 GEF assays to profile the GTP loading activities on all 44 top ranked compounds derived from molecular docking-based high throughput virtual screening.

This GEF-based activity screening revealed a spectrum of Cdc42 HMs that can be categorized into five classes of different functionalities (**Fig. 5A-J**; Supplemental **Table S3** and **Fig. S6**). They displayed different modes of pharmacological activity in the absence and presence of GEF, respectively. Among the five classes, three classes were found to possess activities similar to those observed for the model compounds ZCL278, ZCL279, and ZCL367. The compounds of ZCL278 class, or Class I, promoted GTP loading to Cdc42 in the absence of GEF but reduced this activity in the presence of GEF (**Fig. 5A** and **B**). The compounds of ZCL279 class, or Class II, elevated GTP loading to Cdc42 at the concertation up to low micromoles (from 0.2 nM to 10 µM) but suppressed GTP loading at higher concentrations with and without GEF (**Fig. 5C** and **D**). The compounds of ZCL367 class, or Class III, suppressed GTP loading to Cdc42 with and without GEF (**Fig. 5E** and **F**).

**Figure 5.**
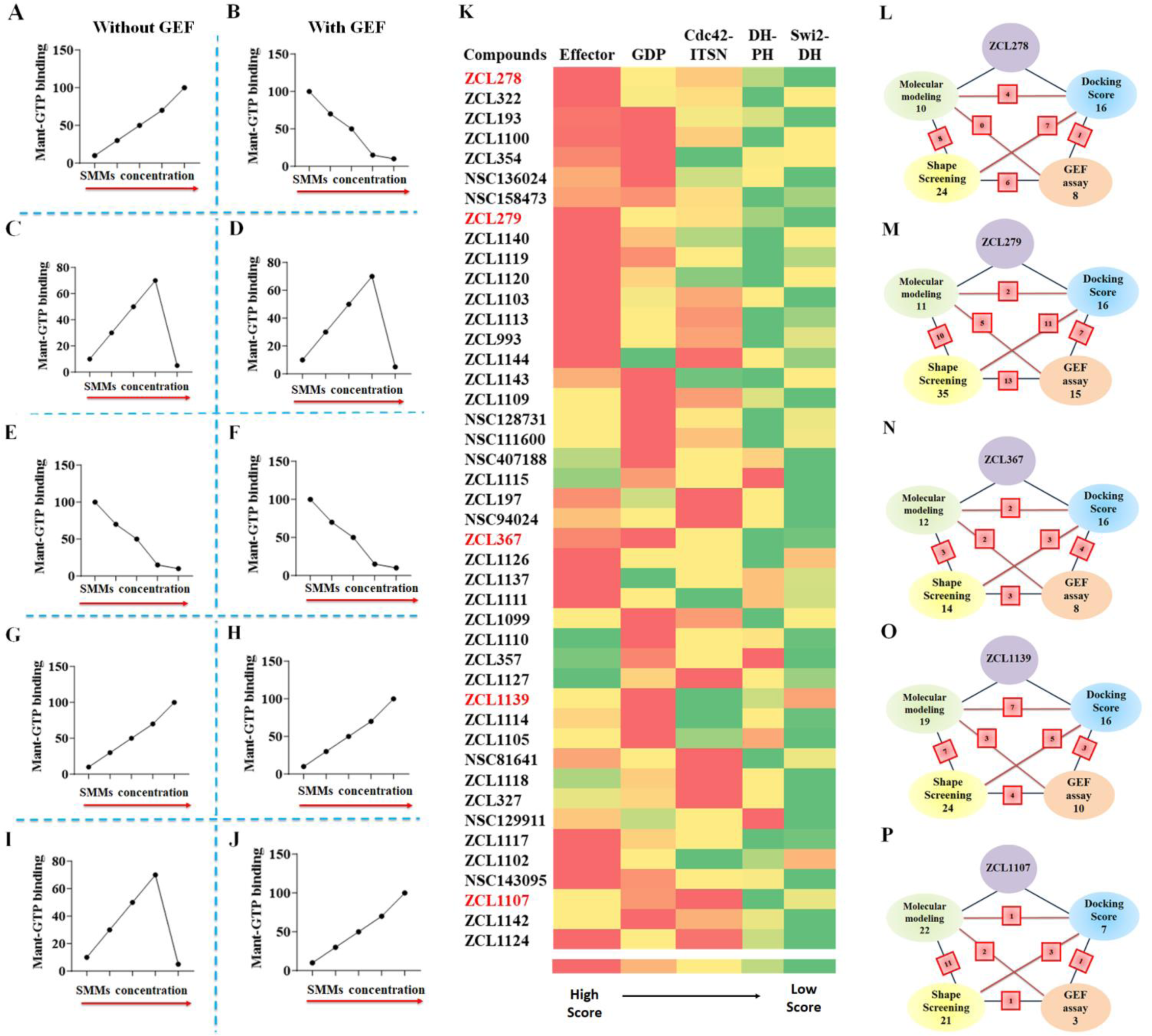
Holistic profiling of Cdc42 HMs using biochemical functionality-based screening and CADD-based modeling. **(A-J)** Schematic representation of all five functionally distinct classes of Cdc42 HM from SMM library by Cdc42GEF assay screening. The red arrows indicate increasing concentration. **(A-B)** ZCL278-like compounds (Class I) increased GTP loading to Cdc42 in the absence of GEF but decreased GTP loading to Cdc42 in the presence of GEF, respectively. **(C-D)** ZCL279-like compounds (Class II) elevated GTP loading to Cdc42 at the concertation up to low micromoles (from 0.2 nM to 10 uM), but inhibited GTP loading to Cdc42 at higher concentrations with and without GEF. **(E-F)** ZCL367-like compounds (Class III) inhibited GTP loading to Cdc42 in the absence and presence of GEF. **(G-H)** ZCL1139-like compounds (Class IV) increased GTP loading to Cdc42 in the absence and presence of GEF. **(I-J)** ZCL1107-like compounds (Class V) elevated GTP loading to Cdc42 at the concertation up to low micromoles (from 0.2 nM to 10 uM), but inhibited GTP loading to Cdc42 at higher concentrations in the absence of GEF. However, they further enhanced GTP loading to Cdc42 in the presence of GEF. **(K)** Color coded spectrum showing PBPO of each compound in SMM library for classification of HMs based on dockingscores. Five identified ligand-protein bindingpockets in interface of Cdc42 and ITSN were used to design PBPO for SMMs. Red reflects the best docking score (the most preferable pocket), whereas green reflects a minimum docking score (the less preferable pocket). The first preferential binding pockets of Cdc42-ITSN complex for 16 compounds were GDP pockets whereas 17 compounds showed maximum docking score into Effector binding pockets. (**L-P)** Schematic summary of classification of SMMs into **(L)** ZCL278-Class I, **(M)** ZCL279-Class II, **(N)** ZCL367-Class III, **(O)** ZCL1139-Class IV and **(P)** ZCL1107-Class V using four different screening methods. Green: Molecular modeling; Blue: Molecular docking; Yellow: 3D Phase Shape; Pink: Cdc42GEF assay. For each class, the number of identified compounds based on applied screening methods are shown. The red square boxes show the shared number of same compounds identified between the two connected screening methods. Compound ZCL993 was identified by all *in silico* and Cdc42GEF-based screening methods as a ZCL279-like Class II compound.

The GEF-based activity screening identified two additional novel classes of compounds that displayed different modes of action from the ZCL278 (Class I), ZCL279 (Class II), and ZCL367 (Class III) classes. The compounds of ZCL1139class, or Class IV, promoted GTPloading to Cdc42 regardless of GEF presence or not (**Fig. 5G** and **H**), thus presenting the features of *bona fide* activators or agonists. The compounds of ZCL1107 class, or Class V, promoted GTP loading to Cdc42 at low concentrations up to 10 µM, but reduced GTP loading to Cdc42 at high concentrations in the absence of GEF, much like ZCL279 class or Class II (compare **Fig. 5I** with **Fig. 5C**). However, when GEFis present, the Class V compounds promoted GTPloadingto Cdc42 even at 100 µM concentration (**Fig. 5J**).

## CADD profiling of HMs

To explore whether there is potential relationship between the experimental classification of HMs based on their modulation of GEFactivity and the CADD predictedclassification of HMs, we applied several Schrödinger-enabled analytic solutions.

We first used classic docking score for prediction of the preferential binding pocket order (PBPO) in Cdc42-ITSN for our model compounds. Then, this PBPO strategy was appliedto profile the top ranked 44 compounds derived from the high throughput screening of SMM libraries, which was based on the five binding pockets in interface of Cdc42-ITSN (**Fig. 2C**). The PBPO of Cdc42 and Cdc42-ITSN for all 44 compounds was thus determined. Most compounds showed preferential binding to Effector and GDP binding pockets (**Fig. 5K**). We compared the PBPO of all identified HMs to that of each model compound. For example, PBPO of Cdc42-ITSN for ZCL279 is: Effector > Cdc42-DH domain of ITSN > GDP > DH-PH domain of ITSN > Switch 2-DH domain of ITSN. Among the compounds from the existing HM libraries, ZCL993 and ZCL1124 were found with having the same PBPO as ZCL279. This method was also employed to assign compounds to ZCL278, ZCL367, ZCL1139 and ZCL1107 classes based on their PBPO (**Fig. 5K**).

We additionally applied a structure-based pharmacophore hypothesis generation for the model compounds. The docked ZCL278 and ZCL279 on Effector pocket, ZCL367 and ZCL1139 on GDP pocket of Cdc42-ITSN as well as ZCL1107 on Cdc42-DH domain of Cdc42-ITSN complex (their preferential binding pockets) were used to build pharmacophore hypothesis using Phase module of Schrödinger suite (*36*). Pharmacophore modeling provides six built-in types of pharmacophore features: hydrogen bond acceptor (A), hydrogenbond donor (D), hydrophobic (H), negative ionizable (N), positive ionizable (P), and aromatic ring (R). The identified pharmacophoric hypotheses features for ZCL278, ZCL279, ZCL367, ZCL1139 and ZCL1107 were RNRAR, RNDRRR, DDRDDR, RAARD and RAA containing five, six, four and three features, respectively (See **Supplementary Fig. S7A-E)**. We examined all 44 HMs and identified for matches with pharmacophore hypotheses of ZCL278, ZCL279, ZCL367, ZCL1139 and ZCL1107 (See **Supplementary** T**able S4**). For example, ZCL993 was again found with having matched pharmacophoric feature to ZCL279.

Shape-based methods to align and score ligands were also employed for flexible ligand superposition and virtual screening as ligand-based classification (*37, 38*). Using Schrödinger’s Phase Shape, 3D ligand alignments for model compounds and the rest of the 44 HMs with the matching scores to assign HMs to ZCL278, ZCL279, ZCL367, ZCL1139 and ZCL1107 classes are shown in Supplementary Data (**Table S4** and **Fig. S7F-J**). For instance, ZCL993 and some other compounds were identified with similar shapes to ZCL279. Finally, a hierarchical clustering analysis of the 44 HMs using Tanimoto similarities index was performed (*39, 40*). From a total of 9 classes, ZCL278, ZCL279 and ZCL367 belong to the largest cluster containing 29 compounds (see **Supplementary Table S5** and **Fig. S8**). Some compounds were found to be functionally similar to the model compounds ZCL278, ZCL279, ZCL367 by most of the CADD profiling prediction **(Supplementary Fig. S9**).

The CADD analysis with PBPO revealed similar HM profile in a color-coded spectrum to these five distinct classes of Cdc42 HMs identified by GEF-based activity profiling (**Fig. 5A-K**). Although more data points may help their better match-up, this comparison provided some insights how to best find functionally similar compounds to the five distinct classes that were identified in our primary high throughput screening. In a schematic overview, the number of identified HMs based on the screening methods in the intersection of ITSN and Cdc42 is shown (**Fig. 5L-P**). Some compounds that show similarities to the model compounds were identified as the same compounds using different screening methods (See **Supplementary Table S4**). For instance, NSC136024, NSC158437, NSC94024 were identified as ZCL278-like Class I compounds based on both GEF activity assay and Shape screening methods. Also, in ZCL279-like Class II compounds, ZCL993 was found by all employed screening methods. For ZCL367-like Class III compounds, ZCL1111 was identified according to GEF activity assay and molecular modeling screening methods. Molecular modeling, Shape screening and GEF assay revealed that ZCL1105 could be in ZCL1139-like Class IV and ZCL1142 was similar to ZCL1107-like Class V.

Finally, we conducted the same comparison of experimental classification and the CADD predicted profiling on top ranked library compounds for targeting Cdc42 (without ITSN) (See **Supplementary Table S6** and **Fig. S10**). Interestingly, the class prediction for some compounds was the same whether they target Cdc42 with or without ITSN. For instance, ZCL993, ZCL1105 and ZCL1142 were identified by the majority of screening methods to be in ZCL279-, ZCL1139- and ZCL1107-like classes respectively. In another aspect, this analysis showed the impact of each screening method to find a compound which can mimic the pharmacological functionality of interests (**Fig. 5**). The average rate of prediction of each screening methods for all five HM classes are as follow: Shape screening (54.2%) > Molecular docking (34.2%) > Molecular modeling (30.8%). Considering that the number of identified compounds by different screening methods are not identical and studying more compounds can lead to increased probability of matching model like compound classes, we calculated the predicting rate based on the number of identified compounds whichare the samebetween GEFassay and other screeningmethods compared to total number of identified compounds for each screening methods. The calculated prediction rate per number for molecular docking was comparable to that for Shape screening (21.4 and 20.6 for molecular docking and Shape screening respectively). The results showed that the PBPO docking was the most effective CADD method for prediction of an HM category that can mimic the pharmacological functionality based on GEF-activity screening.

## Discussion

HMs such as PAs play significant roles in modern therapeutics in human disease management. For example, Buspirone, a dopamine and serotonin 5-HT receptor modulator, is a PA to treat neurodegenerative diseases, psychiatric disorders, anddrugaddiction (*4, 41*). However, prominent PAs are often discoveries towards limited categories of proteins like membrane receptors and ion channels. PAs have especially been rare encounters for *bona fide* cytoplasmic targets, such as small GTPases, which are essential players in many cellular functions from cancer cell proliferation (e.g., Ras oncogene) to neuronal synaptic remodeling (Rho, Rab and Arf in neurodegenerative diseases) (*9, 31*). The current study enabled a systemic identification and classicization of HMs including PAs repertoire for Cdc42; thus, we highlight a new model path to actively search and profile HMs in potentially any protein-protein interaction (PPI) landscape.

This novel enabling classification approach could have far-reaching implications, especially for targeting the two most devastating chronic human diseases such as cancer and neurodegenerative diseases. These diseases likely develop as results of dysregulation of multifaceted biological processes. For example, large omics data are now available that demonstrate many genedysregulations arespatially affected and cell/tissue selective. In AD mouse models, we showed that small Rho GTPase signaling is even dysregulated at both spatial and planar levels in brain (*8*). Therefore, homeostatic targeting may have profound benefits in addition to the traditional and simple inhibitory or stimulatory treatment regimens. Fig. **5A-J** showed that the five classes of Cdc42 modulators cover wide range of effects on GTP loading depending on whether GEF activation is concurrently present or not. It is possible that in the diseased tissues, some HMs may have the capability to stimulate where Cdc42 is over-suppressed or reduce where Cdc42 is over activated. These HMs can thus target diseased cells or tissues while protect tissues that are unaffected.

Our findings show clear feasibility to now move *ab initio* hit identification from relying on binding and PPI to activity-based functional space. This is significant because binding strength does not always lead to prediction of desired pharmacological functions while activity-based functionality assays are labor intensive for initial screening. Our studies show that we are now at the doorstep for linking artificial intelligence/machine learning (AI/ML) tools in CADD to experimental identification and classification. Hence, our findings are encouraging in that with the increased database, AI/ML-based models can be used for predicting novel HM classes much like the AlphaFold for protein structure prediction (*42–44*). Thus, our studies may indeed lead to paradigm shift in how to approach chemical space in drug discovery.

Another implication of our findings is that now we can anticipate performinga single study to enable profiling of all HM classes in an entire PPI modulator repertoire for a given target, significantly reducing the duplicating efforts in identifying hits of interests for a specific functionality. This would favor shifting the burden of discovery to early stage to enhance desired hit identification. With the aid of automation and AI/ML, we will be able to leave behind much fewer corners of the therapeutic landscape that are unexplored and thus accelerate drug discovery for *difficulty-to-treat* diseases such as cancer and ADRD.

## Material and Methods

### 1. Reagents and Cell Lines

Cdc42-GEF assay kit (BK100), GLISA kit (BK127) to measure Cdc42 activity, and His-DBS (DH/PH domain) were purchased from Cytoskeleton, Inc. (Denver, CO). For GLO assay, GTPase-GLO assay kit were purchased from Promega (V7681) (Madison, WI). Swiss 3T3 cells were cultured as described. ZCL278 was purchased from Cayman chemical (Ann Arbor, MI) or synthesized in the laboratory. ZCL279 and ZCL367 were synthesized in the laboratory. Some SMMs were obtained from National Cancer Institute Experimental Therapeutics Program. Swiss 3T3 cells were purchased from American type Culture collection (ATCC) (Manassas, VA) and cultured as described. Compounds were administered to cells in 5% DMSO (Sigma Aldrich).

### 2. Experimental methods

#### 2.1. Cdc42-GEF assay

Cdc42-GEF activation assay (Cytoskeleton) was performed per manufacturer’s instructions (*26*). Briefly, a solution was prepared that contained testing compounds (0.01–100 μM) and a mixture of purified Cdc42 GTPase, exchange buffer with Mant GTP, and the nanopure water. The fluorescence in 485/535 nm for excitement/emission was monitored to obtain the background/baseline. Then, the GEF protein was added and the absorbance was recorded for an additional on Biotek Synergy HT or Biotek Synergy H1 (Agilent, Winooski, VT). DMSO was employed as vehicle control. The linear slopes were calculated by using GraphPad Prism version 10.0.2(232) (La Jolla, CA).

#### 2.2. G-LISA assay

The Cdc42 G-LISA Activation Assay measures the level of GTP bound Cdc42 protein in cell lysates. Serum-starved Swiss 3T3 cells were treated with different concentrations of ZCL278, ZCL279 and ZCL367 compounds. For time dependent G-LISA, ZCL278 and ZCL279 were tested for different time intervals including 10 min, 20 min, 30 min, 1 hour, 2 hours and 6 hours. For stimulated Cdc42 G-LISA, bradykinin was used to activate Cdc42 in Swiss 3T3 cells, followed by treatment with either ZCL278, ZCL279 or ZCL367 for 15 min, 1 hour or 6 hours. The cells were lysed and protein lysates were used for G-LISA analysis per manufacturer’s (Cytoskeleton) instructions. After placing the collected lysates in a 96 well Cdc42-GTP binding plate, any unbound GTP was washed out and the bound GTP levels were measured on Biotek Synergy HT (Agilent, Winooski, VT).

#### 2.3. GLO assay

GTPase activityassays were carriedout using GTPase Glo assay kit (Promega, V7681)(*45*). GTP-hydrolysis activities of Cdc42 after treatment with HM compounds with or without GEF (e. g. ITSN) were determined. Reactions were initiated by adding 10 µM GTP in GTPase-GAP solution containing 1mM DTT, Cdc42, GAP, and GEF buffer. Reactions were incubated for 90 minutes at room temperature. To the completed GTPase reactions, we added 10 µl of GTPase-Glo™ Reagent and incubated reactions for 30 minutes at room temperature. Then, in the final step, detection reagent was added, and the plate was incubated for 5 - 10 minutes at room temperature. The unhydrolyzed GTP that was converted to ATP was recorded on BioTek Synergy HT (Agilent, Winooski, VT).

### 3. Computational methods

#### 3.1. Ligand preparation

The 44 top ranked HMs which are derived from the high throughput screening of SMM libraries were used for secondary screening. The LigPrep module of Schrödinger software was used to prepare, neutralize, desalt, and adjust tautomer of the library. Then, OPLS4 force field was employed to minimize their energy (*46*)(LigPrep, Schrödinger, LLC, New York, NY, 2023.).

#### 3.2. Molecular docking

Several CADD methods, tools and approaches were employed in this study. All computational studies were carried out by the Maestro version 2022.03 of the Schrödinger Small Molecule suite. Structure- and ligand-based pharmacophore modelling as well as Molecular Dynamics (MD) simulations were performed using Phase module and Desmond version 2022.03 of Schrödinger Small Molecule suite. Briefly, the crystallographic structure of Cdc42-ITSN complex (PDB code: 3QBV) was downloaded from Protein Data Bank at RCSB (https://www.rcsb.org/) (*47*) and prepared using the protein preparation wizard module of the Schrödinger software package (*48*). After the introduction of hydrogen bonds, missing side chains and loops were filled. Subsequently, the OPLS4 force field was employed for optimization and minimization. Site Map feature was used for detection of binding pockets of Cdc42-ITSNcomplex (*49*).

Next, the binding cavities were determined using the receptor grid generation module around the probable binding pocket. The docking studies were conducted using the Glide module of Schrödinger software. Initially, The Glide Extra Precision(G-XP) mode was usedfor predicting the bindingmode of HM compounds, and the top-ranked output was analyzed (*50*). Then, to search and score all 44 compounds, the screened library was subjected to docking using high throughput virtual screening (HTVS) mode of Glide. The obtained docking scores were color coded.

#### 3.3. Molecular dynamics (MD) simulation

MD simulation was performed to elucidate the effectiveness of the screened compounds by molecular docking. MD simulations were carried out using Desmond module of Schrödinger suite v2022-3. The simulation box contained the TIP3P water as solvent whereas simulation time, temperature and relaxation time were 500 ns, 300 K and 1 ns, respectivelyfor theligands (ZCL278, ZCL279 and ZCL367) on the NPT ensemble platform. The OPLS_2004 force field parameters were used in all simulations. The water molecules were explicitly indicated using the simple point charge model. The TIP3P water model in an orthorhombic shape was selected after minimizing the volume, with 15 Å × 15 Å × 15 Å periodic boundary conditions in the Protein-ligand complex’s x, y, and z-axis. The cutoff radius in Coulomb interactions was 9.0 Å. The Martyna-Tuckerman-Klein chain coupling scheme with a coupling constant of 2.0 ps and the Nosé - Hoover chain coupling scheme were employed for the pressure and the temperature control respectively. An r-RESPA integrator where the short-range forces were updated every step and the long-range forces were updated every three steps were used for calculation of nonbonded forces. The trajectories were saved at 500 ps intervals for analysis. The ligand andprotein interactions wereanalyzed using the Simulation Interactiondiagram tool implemented in Desmond MD package. The RMSD of the ligand and protein atom positions with time were employed to monitor the stability of MD simulations.

#### 3.4. E-pharmacophore modeling and virtual screening

In the Phase module, the E-Pharmacophore generation panel was opened and the option “create a pharmacophore model using receptor-ligand complex” was selected to create a pharmacophore model from the Cdc42-ITSN complex with HM compounds. Subsequently, the SMM library was screened against the generated pharmacophoric features *via* the Phase Ligand Screening panel, resulting in a group of compounds that matched the features (*51, 52*).

#### 3.5. Shape-based screening

The Shape-based model of HM compounds were generated by theshape screeningin Phase module of Schrödinger version 2022-03. Then, 3D-screening methods were employed for screening of the 44 HM compounds. Based on each query of HM model compounds, a group of compounds were identified with matched features to the HM model compounds.

#### 3.6. Clustering of HM compounds in the screened library

Using Schrödinger Canvas suite version 2022-03, a hierarchical clustering analysis for screened HM library was carried out (*40, 53*). The molecular fingerprint was calculated from the two-dimensional structure of the compounds in the form of <u>e</u>xtended <u>c</u>onnectivity <u>f</u>inger<u>p</u>rint 4 (ECFP4). The metric of the Tanimoto similarity and the average cluster linkage method, which clusters according to the average distance between all inter-cluster pairs, were employed for a hierarchical clustering analysis.

### 4. In vivo Study

#### 4.1. Transgenic mice

The Institutional Animal Care and Use Committee (IACUC) of East Carolina University reviewed and approved the studies involving use of animals. 3xTg-AD mouse model contained the familial AD mutations in amyloid precursor protein (APPswe), microtubule-associated protein tau (TauP301L), and presenilin1 (PS1M146V), whereas a non-Tg B6;129SF1/J was used as a wildtype (WT) control for the studies. The Jackson Laboratory (Bar Harbor, ME) provided both mouse groups(*32*). Mice werebredandhoused in groupsof two to four in transparent polyethylene cages with free access to food and water. PCR analysis on the tail tissue was performed in accordance with the animal use protocol (AUP) to confirm the genotype of the colony.

#### 4.2. Mouse tissue fixation and preparation

Mice were first anesthetized using isoflurane followed by perfusion of the brain with ice cold PBS followed by 4% Paraformaldehyde (PFA: sc-281692, Santa Cruz Biotechnology, Inc., CA) in PBS. The brains were removed, transferred to 4% PFA for 48 hours, and then moved to a 30% sucrose (sx-1075-1, EMD Millipore, MA) solution which acted as a cryoprotectant. Once the brains settled to thebottomof thestorage vials, they were embedded in optimal cuttingtemperature compound (O.C.T) (Sakura 4583) and frozen with cold isopentane. Using a cryostat, 8 µm serial coronal slices of brain tissue were taken.

#### 4.3. Behavioral evaluation

Six-month-old C57/BL6 (WT) and 3xTg-AD mice with an average weight of 24 g from Jackson Laboratory (Bar Harbor, Maine, United States of America) were used. All mice were subjectedto two weeks of trainingfor familiarizingbehavioral test procedures. Subsequently, mice were assayed to establish a baseline of behavior tests before HM compound treatment. Then, mice were separated into four treatment groups (Control, ZCL278, ZCL279, and ZCL367). All compounds were dissolved in sesame oil with 5% DMSO and each treatment was administrated via an intraperitoneal injection. All treatment groups of mice were injected every other day except weekends. Control group was treated with 5 % DMSO in sesame oil. HM compound groups were treated with 20 μg/g ZCL278, ZCL279, or ZCL367 in sesame oil. On the day following drug treatment, all mice were subjected to social interaction evaluations for four weeks. The mouse behavioral status was scored usinga relative scale of 1-5 (1 is Calm, 2 is Responsive, 3 is Restless, 4 is Agitative, and 5 is Aggressive).

#### 4.4. Gene expression analysis

Gene expression was analyzed using Geomx mouse whole transcriptome atlas (Nanostrings Technology, Inc) per manufacturer’s instruction. Briefly, 5 µm thick brain sections were mounted on super frost slides. The slides were baked in a 60°C oven for 30 minutes. Slides were rehydratedby submergingin 100% ethanol to 95% ethanol and finally PBS. Antigen retrieval was performed on the slides using 1x Tris-EDTA (pH 9.0). Slides were incubated with Proteinase K solution at 37°C for 15 minutes, after which they were washed with PBS for 5 minutes. The slides were post fixed with a 10 % (Neutral Buffered Formalin) NBF buffer. Slides were then incubated with RNA detection probes which were bound to photocleavable oligonucleotide tags. The slides were incubated with the probes overnight at 37°C. Slides were washed to remove off target probes. The oligonucleotide tags were then released from specific regions of the tissue using ultraviolet exposure. As 61 barcodes were released, a microcapillary system was used to collect the tags which were then transferred to a microtiter plate. The barcodes were read on the Illumina sequencer. A total of 19,963 genes were investigated in 52 regions of interests (ROIs). Any gene that was above the limit of quantification of confidence threshold LOQ2, defined as the geometric mean multiplied by the geometric standard deviation to the second power, was included. A total of 10,560 genes were expressed above the LOQ2. To reduce variance from segment size and the differential segment cellularity, the count values obtained were normalized to the third q uartile (Q3) of selected targets that were above the limit of LOQ. Q3 normalization divides the number of counted probes in one segment by the ^3r^d quartile value for that segment.

#### 4.5. Protein expression analysis

Fixed-frozen mouse coronal brain sections were incubated with a cocktail of 87 AD related protein antibodies followed by treatment of photocleavable oligonucleotide containing tags. The ROIs on the tissue were selected and the selected area were illuminated by ultraviolet light to dissociate the oligonucleotide tags that are bound to the antibody of interest. The dissociated tags were then collected in a 96 well plate and analyzed. The data was normalized using area normalization in which the counted number of cleaved probes was normalized to the geomean of the area selected to correct for variability of surface area.

### 5. Statistical analysis

Statistical analyses were performed using GraphPad Prism version 10.0.2(232) (La Jolla, CA). For typical invitro studies, data were presented as mean±SEM from at least two independent experiments with a total of four replicates. For initial method development using HM model compounds, more than three independent experiments were conducted. For in vivo studies, data are presented as mean±SEM from replicates and are representative of results from experiments conducted at least twice. *P*-values <0.05 were considered significant. Differences between HM treated groups were analyzed using one-way ANOVA. Also, differences between WT or 3xTg-AD mice were analyzed using an unpaired Student’s *t*-test. Comparison of mice with different drugtreatments over a periodof four weeks was performed by usinga two-way ANOVA. *P*-values < 0.05 were considered significant.

## Acknowledgments

We thank Byron Aguilar, Yi Zhu, and Joani Zary-Oswald for technical assistance.

## Author contributions

S.M. and F.A. contributed equally to this study. S.M. conducted all Cdc42GEF and G-LISA assays as well as Molecular docking and Molecular dynamics analyses. F.A. conducted GTPase GLO assay and CADD-based PBPO including Molecular docking screening, molecular modeling, Shape screening and clustering. S.M. and F.A. cross-validated each other’s analyses. S.N., S.M., F.A., and C.B. performed *in vivo* mouse experiments. C.B. and Q.L. conducted Swiss 3T3 cell culture experiments with imaging analyses. S.N. analyzed GeoMx protein and transcriptomics data. Q.L. and Y.H.C designed the study and performed data analyses. J.T. discussed, wroteand editedthe manuscript. All authors contributedto the writingof the manuscript. F.A. and Q.L. compiled the references.

## Data and materials availability

All data supporting the findings of this study are available in the main text and/or the supplementary materials. The identities of some compounds docked in this study and the docking results, including number, SMILES, and docking score, are available upon request from the corresponding author in accordance to institutional technology safety and privacy policy.

## Supplementary materials

Figs. S1 to S10

Tables S1 to S6

---

**Supplementary Figure S1.**
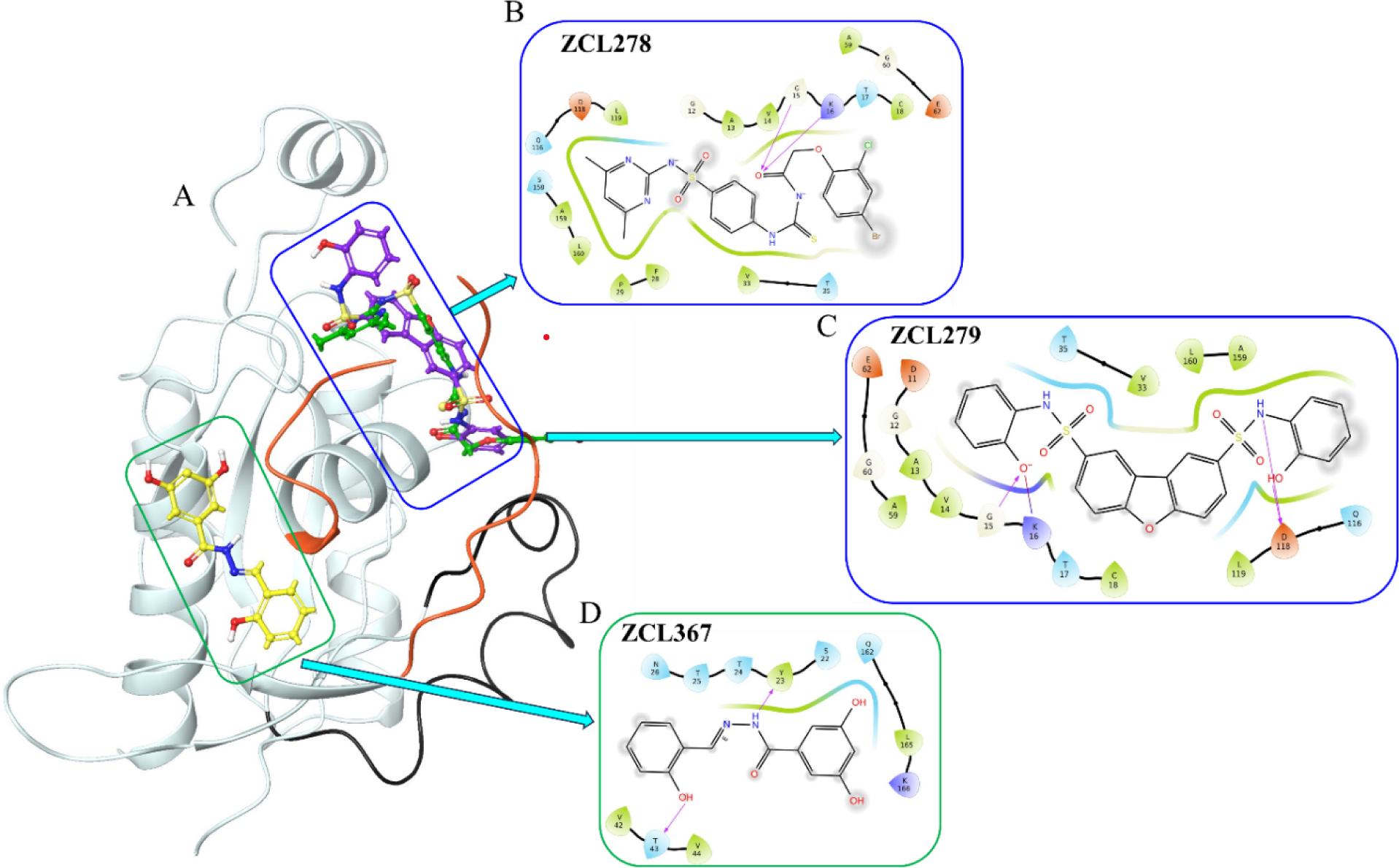
Superimpose of model compounds ZCL278, ZCL279, and ZCL367 and their interactions with Cdc42. **(A)** 3D structural illustration of different model compounds (ZCL278, ZCL279, and ZCL367 are shown in green, purple, and yellow, respectively) on Cdc42 (transparent cyan, where switch 1 (24-40) region is in orange color & switch 2 (57-76) region is in black color) (PDB:3QBV). **(B-D)** 2D representation of active site binding residues of Cdc42 with ZCL278 at GDP site, ZCL279 at GDP site, and ZCL367 at Effector site. The magenta arrows represented hydrogen bonds. Residues at the binding site were depicted in different colors based on the type of interactions like hydrophobic (green), polar (cyan), charged positive (violet) and charged negative (red), respectively.

**Supplementary Figure S2.**
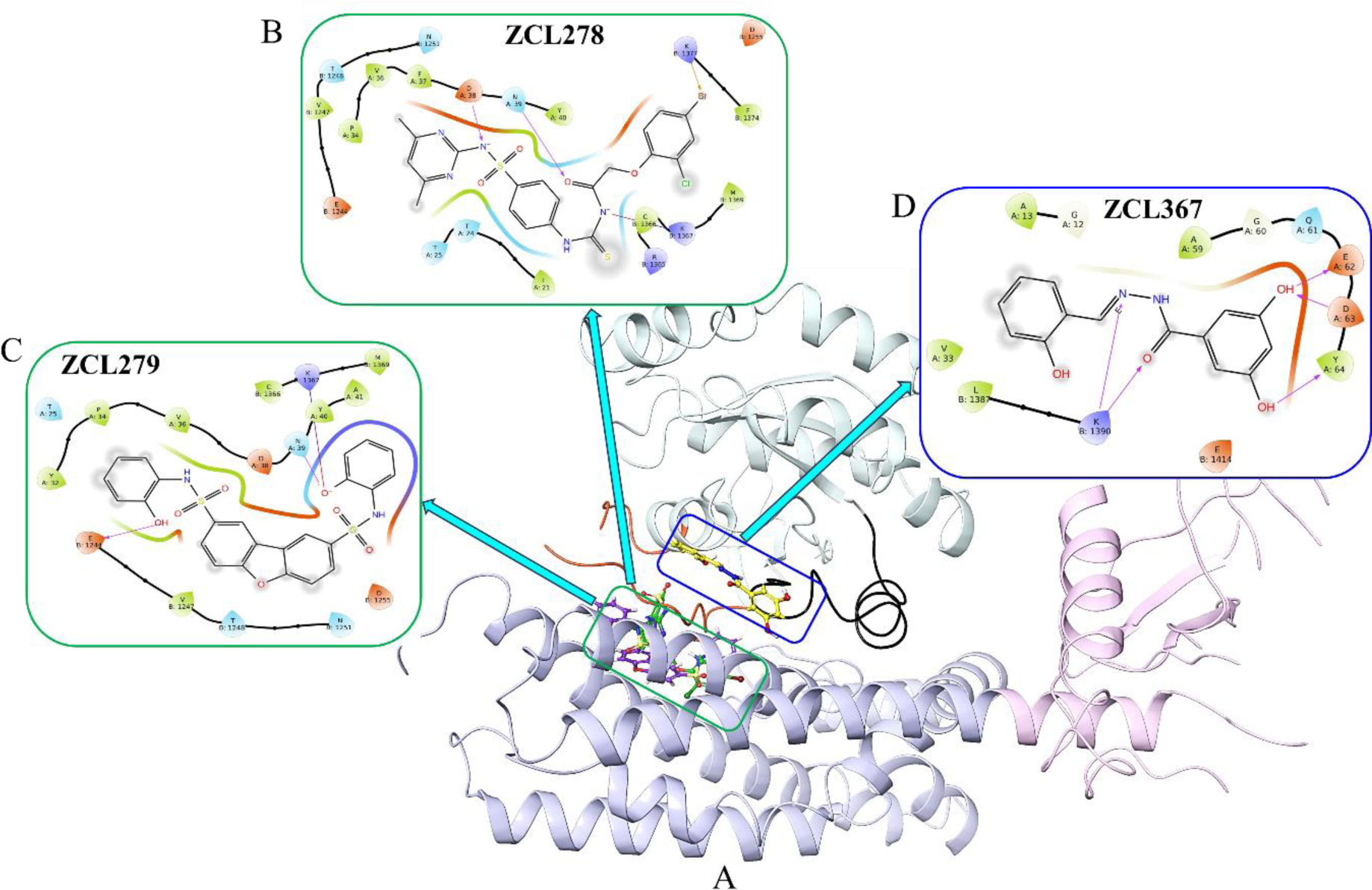
Superimpose of model compounds ZCL278, ZCL279, and ZCL367 and their interactions in the Cdc42-ITSN complex. **(A)** 3D Structural illustration of different model compounds (ZCL278, ZCL279 and ZCL367 are shown in green, purple, and yellow, respectively) on Cdc42-ITSN (transparent cyan, where switch 1 (24-40) region is in orange color & switch 2 (57-76) region is in black color. DH and PH domains of ITSN are shown as purple and pink colored ribbons (PDB: 3QBV). **(B-D)** 2D representation of active site binding residues of Cdc42-ITSN complex with ZCL278 at Effector site, ZCL279 at Effector site, and ZCL367 at GDP site. The magenta arrows represented hydrogen bonds. Residues at the binding site were depicted in different colors based on the type of interactions like hydrophobic (green), polar (cyan), charged positive (violet), and charged negative (red), respectively.

**Supplementary Figure S3.**
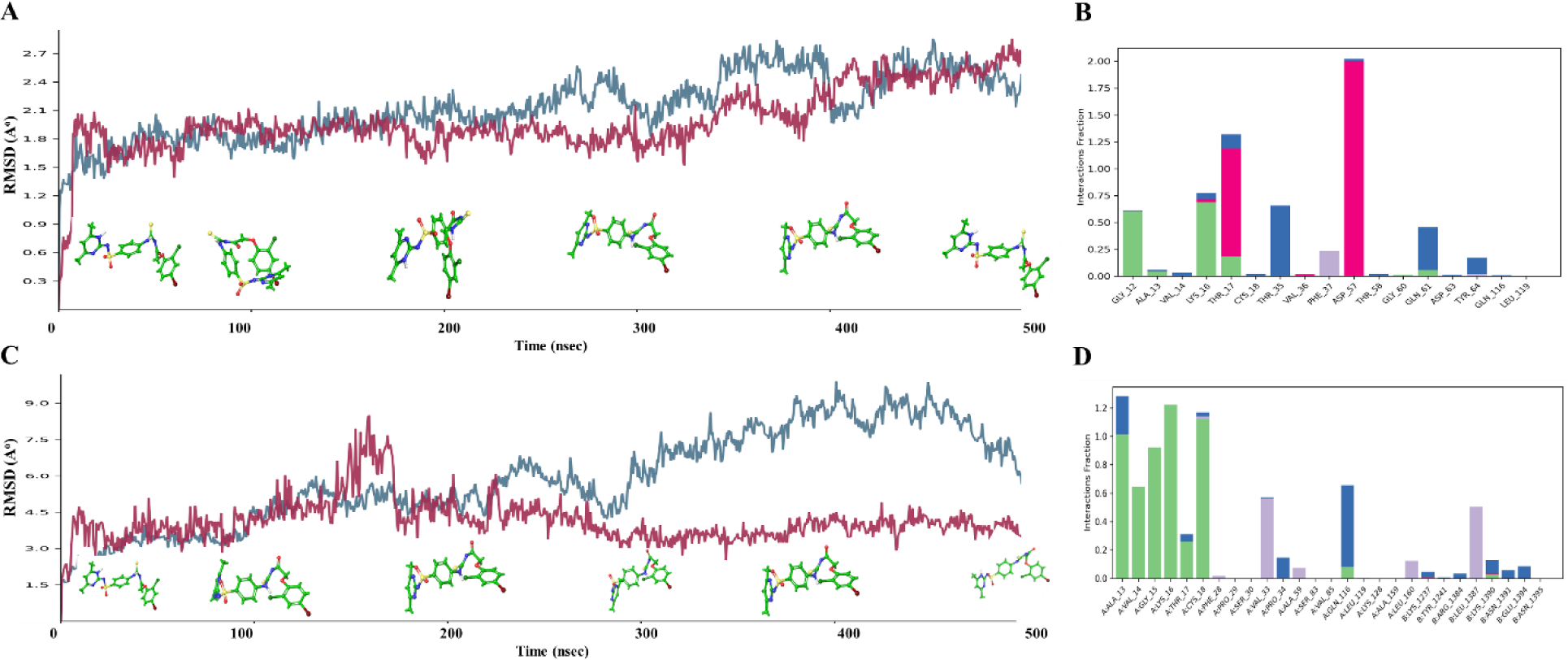
(**A** and **C)** The protein-ligand RMSD plots of the ZCL278 complexed with proteins during 500 ns molecular dynamics (MD) simulation using Schrödinger software Desmond module. Y-axis represented the RMSD values of ligands and protein in molecular distance unit Angstrom, while X-axis demonstrated time in nanoseconds (nsec). The RMSD value of C-alphas (alpha-Carbon) of Cdc42 and Cdc42-ITSN were stabilized from 1.01 A° to 2.4 A° and 1.66 A° to 4.21 A°, respectively, indicating stable conformations. (**A)** Cdc42 (3QBV-chainA) and (**C)** Cdc42-ITSN (3QBV-chainAB). (**B** and **D)** Histogram representation of protein–ligand contacts of ZCL278 and proteins during 500 ns molecular dynamics simulation analysis. Here, X-axis represented the residues and Y-axis represented the interaction fraction. **(B)** Cdc42 (3QBV-chain A) and (**D)** Cdc42-ITSN (3QBV-chain AB). The green color represents H-bond, violet represents hydrophobic, pink represents ionic, and blue represents water bridge interactions.

**Supplementary Figure S4.**
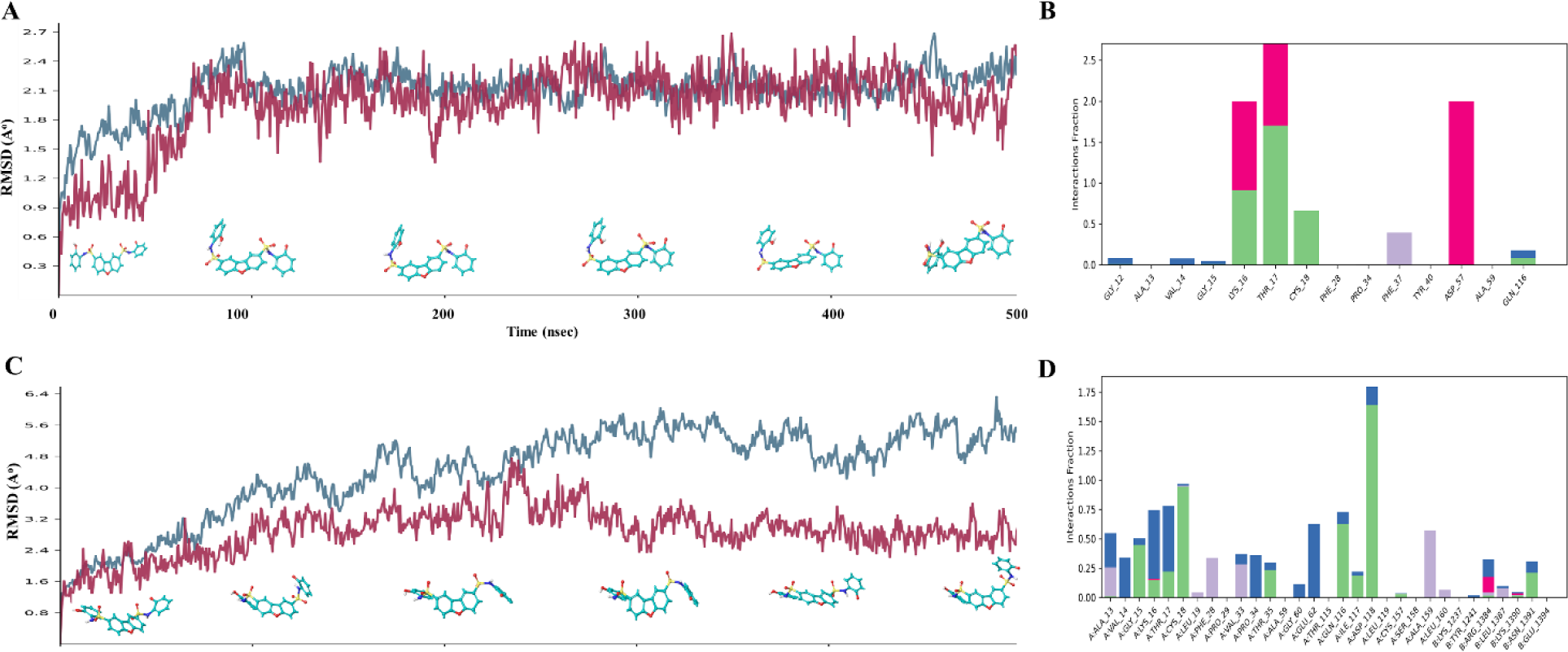
(**A** and **C)** The protein-ligand RMSD plots of ZCL279 complexed with proteins during 500 ns molecular dynamics (MD) simulation using Schrödinger software Desmond module. Y-axis represented the RMSD values of ligands and protein in molecular distance unit Angstrom, while X-axis demonstrated time in nanoseconds (nsec). The RMSD values of C-alphas (alpha-Carbon) of Cdc42 and Cdc42 ITSN were stabilized from 0.4 A° to 2.5 A° and 0.8 A° to 1.8 A° after MD simulation for 500 ns indicating stable conformations. (**A)** Cdc42 (3QBV-chainA) and (**C)** Cdc42-ITSN (3QBV-chain AB). (**B** and **D)** Histogram representation of protein–ligand contacts of ZCL279 and proteins during 500 ns MD simulation analysis. Here, X-axis represented the residues and Y-axis represented the interaction fraction. (**B)** Cdc42 (3QBV-chain A) and (**D)** Cdc42-ITSN (3QBV-chain AB). The green color represents H-bond, violet represents hydrophobic, pink represents ionic, and blue represents water bridge interaction.

**Supplementary Figure S5.**
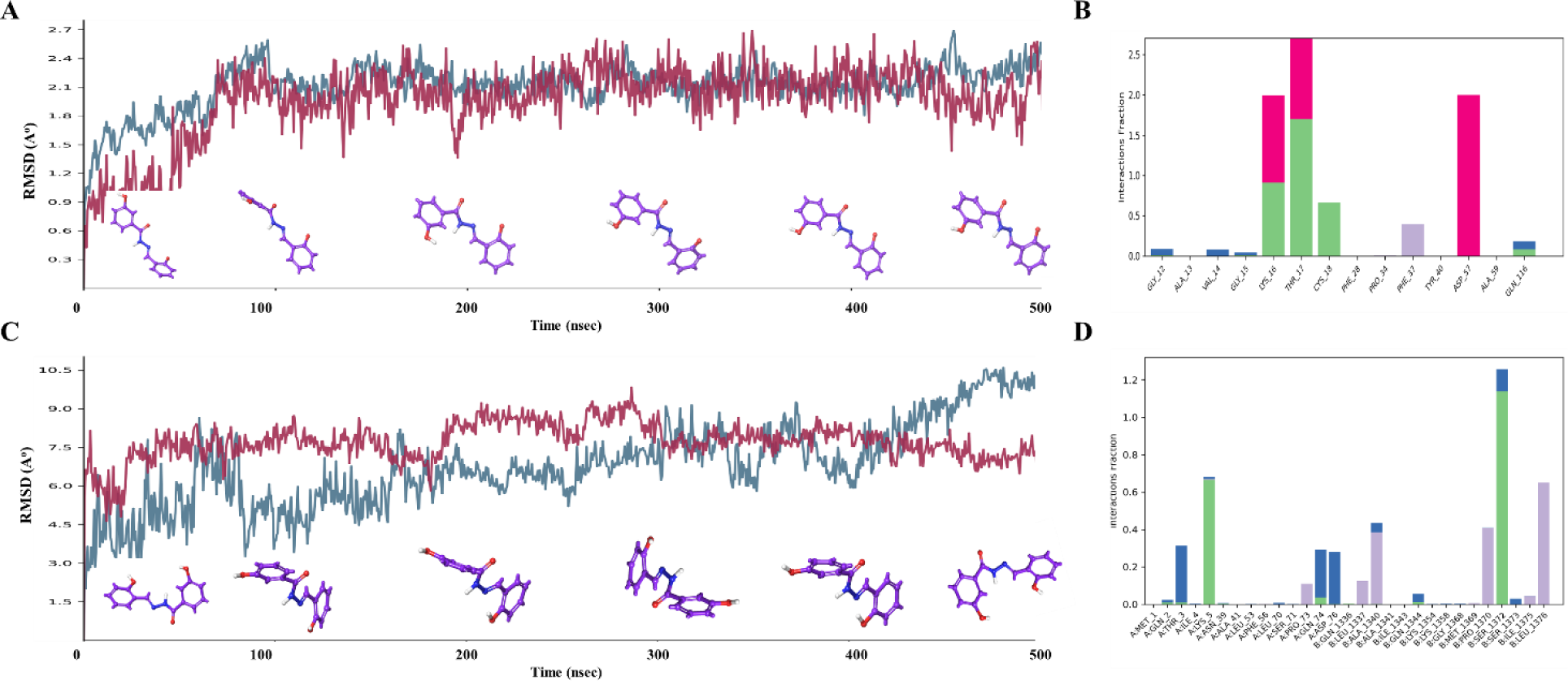
(**A** and **C)** The protein-ligand RMSD plots of ZCL367 complexed with protein during 500 ns molecular dynamics (MD) simulation using Desmond software. Y-axis represented the RMSD values of ligands and protein in molecular distance unit Angstrom, while X-axis demonstrated time in nanoseconds (nsec). The RMSD values of C-alphas (alpha-Carbon) of Cdc42 and Cdc42 ITSN were changed from 0.4 A° to 1.8 A° and 4.0 A° to 6.0 A°, respectively, indicating stable conformations. (**A)** Cdc42 (2QRZ) and (**C)** Cdc42-ITSN (3QBV). (**B** and **D)** Histogram representation of protein–ligand contacts of ZCL367 with protein during 500 ns MD simulation analysis. Here, X-axis represented the residues and Y-axis represented the interaction fraction. Different colors in the histogram represented different types of bond interaction fraction. **(B)** Cdc42 (2QRZ) complex and (**D)** Cdc42-ITSN (3QBV). The green color represents H-bond, violet represents hydrophobic, pink represents ionic, and blue represents water bridge interaction.

**Supplementary Figure S6.**
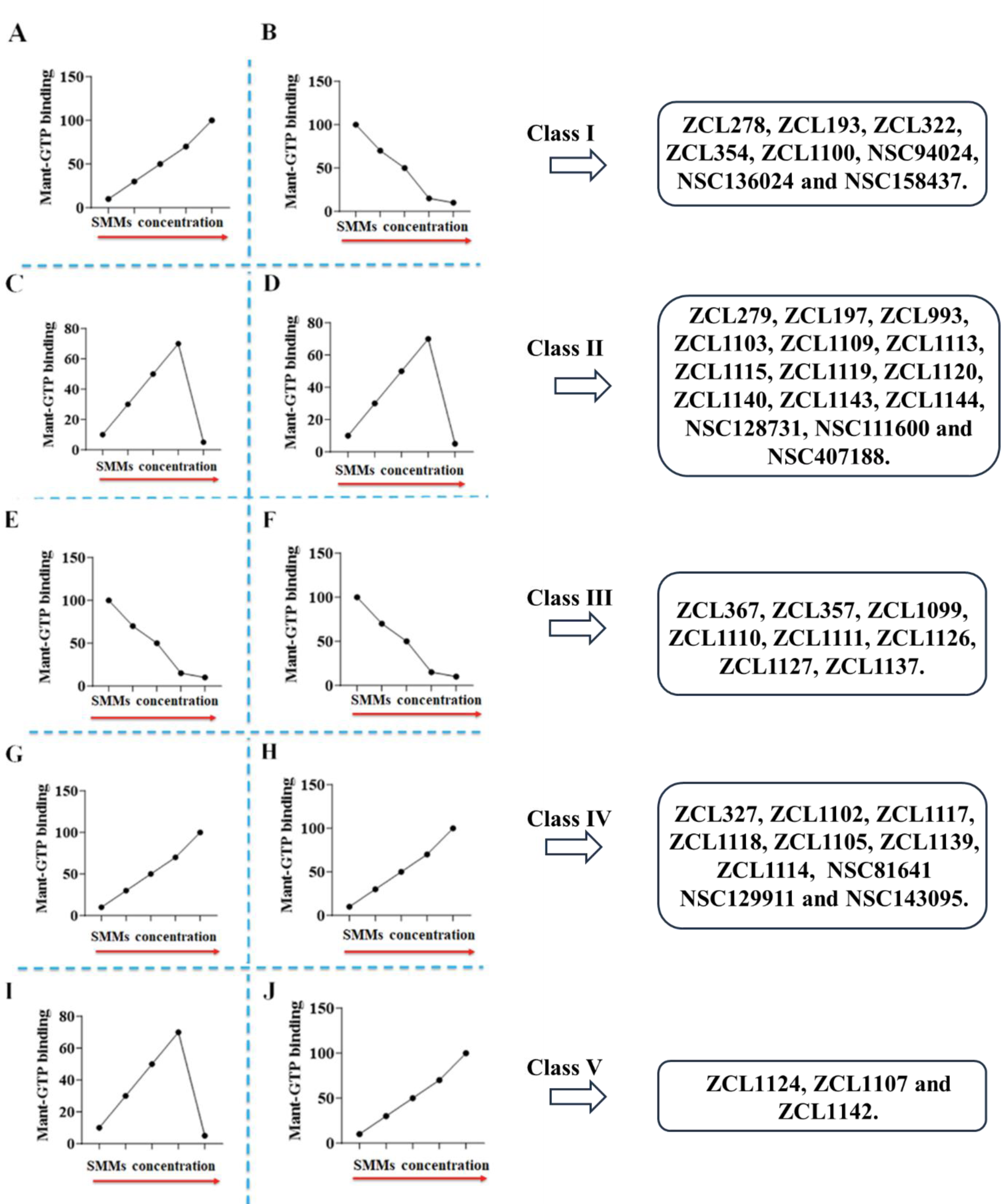
Biochemical classification of the five model compounds and profiling the top ranked compounds from the HM library into these five classes. **A, C, E, G** and **I**: Cdc42GEF assays conducted in the absence of GEF. **B, D, F, H**, and **J**: Cdc42GEF assays conducted in the presence of GEF. The red arrows indicate increasing SMM concentrations; the profiled compounds from the SMM library are shown in boxes for each class. Note: The guanine nucleotide exchange reaction was performed using the relative fluorescence intensity of N-MAR-GTP exchange that monitors GDPreleaseduringthe exchange reactionafter treatment by different concentrations (0.01-100 µM) of each screened compound on Cdc42 in the absence or presence of GEF. The control for all Cdc42GEF assays without GEF presence was DMSO and Cdc42, and for all Cdc42GEF assays with GEF presence was DMSO, Cdc42 and GEF.

**Supplementary Figure S7.**
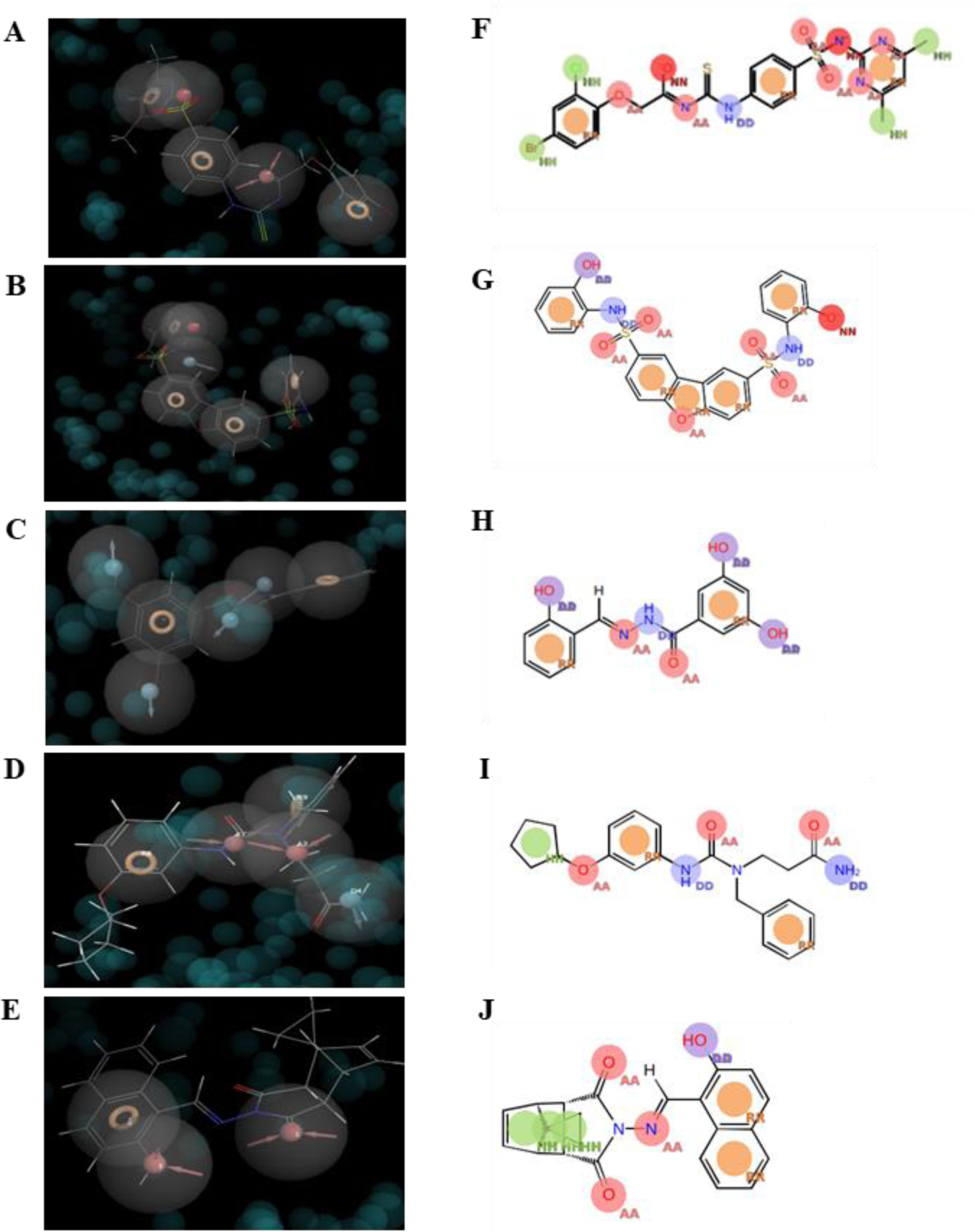
Pharmacophoric hypothesis and Shape screening of Class I-V model compounds. E-Pharmacophoric hypothesis of ZCL278 and ZCL279 on Effector pocket of Cdc42-ITSN, ZCL367 and ZCL1139 on GDP pocket of Cdc42-ITSN, as well as ZCL1107 on Cdc42-DH domain of Cdc42-ITSN complex. They were generated using the Phase module of Schrödinger software. Pharmacophore modeling is shown by hydrogen bond acceptor (A), hydrogen bond donor (D), hydrophobic (H), negative ionizable (N), positive ionizable (P), and aromatic ring(R). The generated models were RNRAR, RNDRRR, DDRDDR, RAARD and RAA for ZCL278, ZCL279, ZCL367, ZCL1139 and ZCL1107 respectively. These models are employed for 3D screening on the HM library compounds to find the best matched compounds with each model. (**A)** ZCL278, (**B)** ZCL279, (**C)** ZCL367, (**D)** ZCL1139, and (**E)** ZCL1107. The shape-based query of (**F)** ZCL278, (**G)** ZCL279, (**H)** ZCL367, (**I)** ZCL1139 and **(J)** ZCL1107 was created using Phase module of Schrodinger software and used for high throughput screening on SMM libraries. AA, DD, NN, RR, HH stands for aromatic ring, double bond, nitrogen atom, ring and hydrogen atom respectively.

**Supplementary Figure S8.**
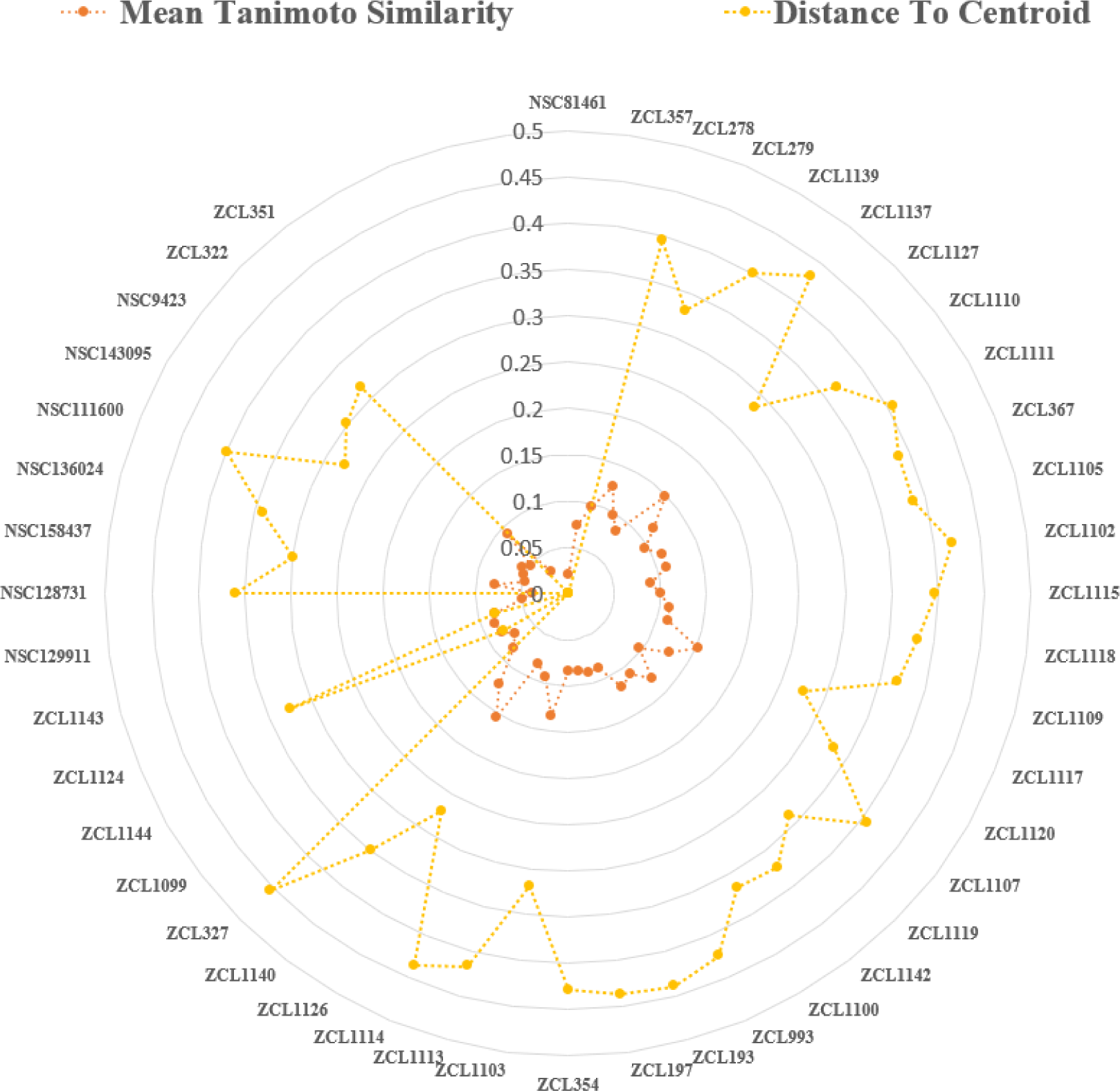
Molecular clustering analysis of Cdc42-ITSN HM repertoire. The similarity analysis of Cdc42-ITSN HM library was performed on the extended connectivity fingerprint 4 (ECFP4) of compounds using Schrödinger Canvas Suite version 2022-03. The metric of the Tanimoto similarity and the average cluster linkage method, which clusters according to the average distance between all the inter-cluster pairs, were employed for a hierarchical clustering analysis. Radial plots show different clusters of screening library based on Tanimoto similarities and distance to centroid for each compound.

**Supplementary Figure S9.**
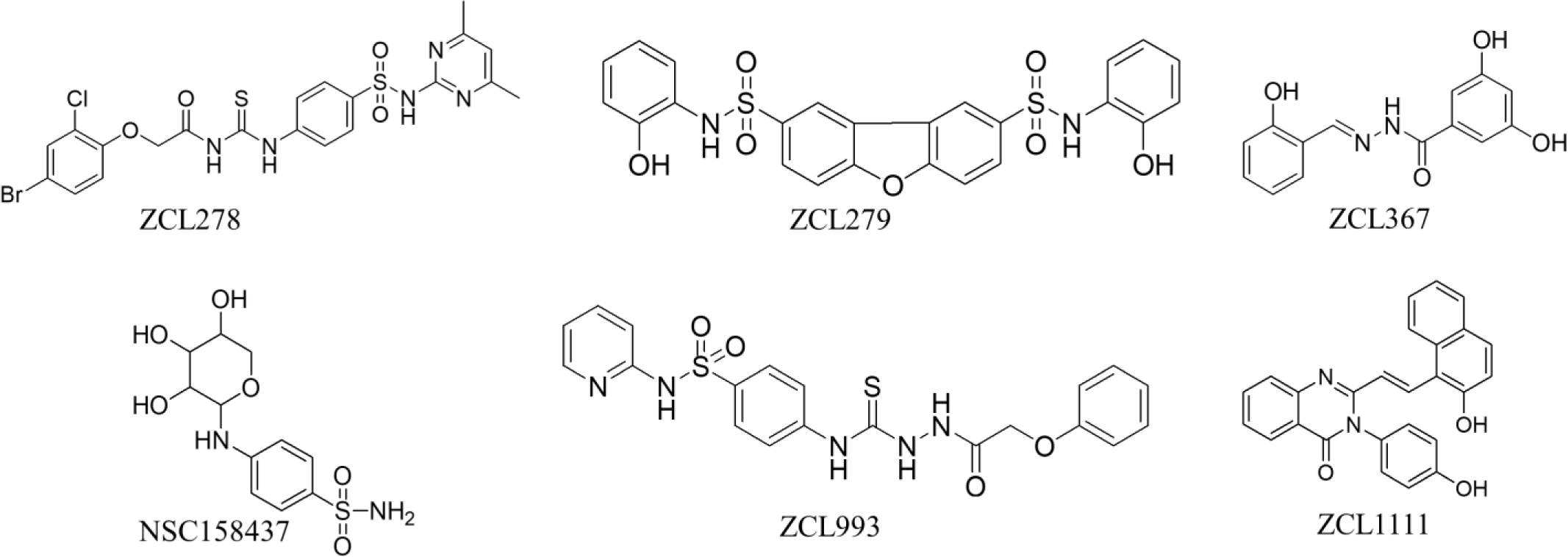
Structures of model compounds ZCL278, ZCL279, and ZCL367 and the predicted compounds with respective similar functions. Structures of ZCL278 and its functionally similar compound NSC158437. Structures of ZCL279 and its functionally similar compound ZCL993. Structures of ZCL367 and its functionally similar compound ZCL1111.

**Supplementary Figure S10.**
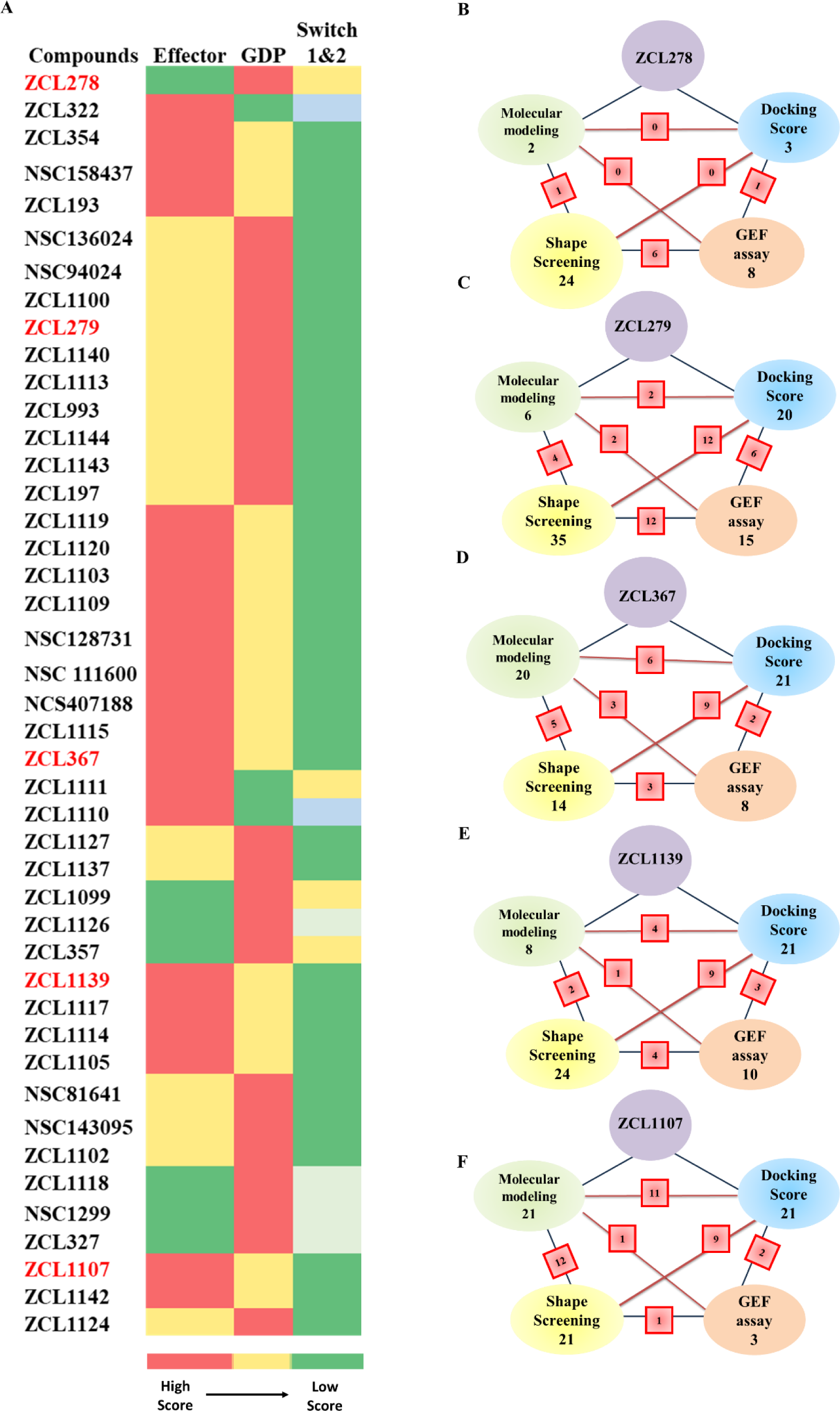
CADD predictability of different screening methods to identify model compound-like HM molecules. (**A**) Color coded spectrum showing PBPO of each compound in SMM library for classification of HMs based on docking scores. Three identified ligand-protein binding pockets in Cdc42 were used to design PBPO for SMMs. Red reflects the best docking score (the most preferable pocket), whereas green reflects a minimum docking score (the less preferable pocket). (**B-F**) Schematic summary of classification of SMMs into **(B)** ZCL278-Class I, **(C)** ZCL279-Class II, **(D)** ZCL367-Class III, **(E)** ZCL1139-Class IV and **(F)** ZCL1107-Class V using four different screening methods. Green: Molecular modeling; Blue: Molecular docking; Yellow: 3D Phase Shape; Pink: Cdc42GEF assay. For each class, the number of identified compounds based on applied screening methods are shown. The red square boxes show the shared number of same compounds identified between the two connected screening methods.

**Supplementary Table S1.**
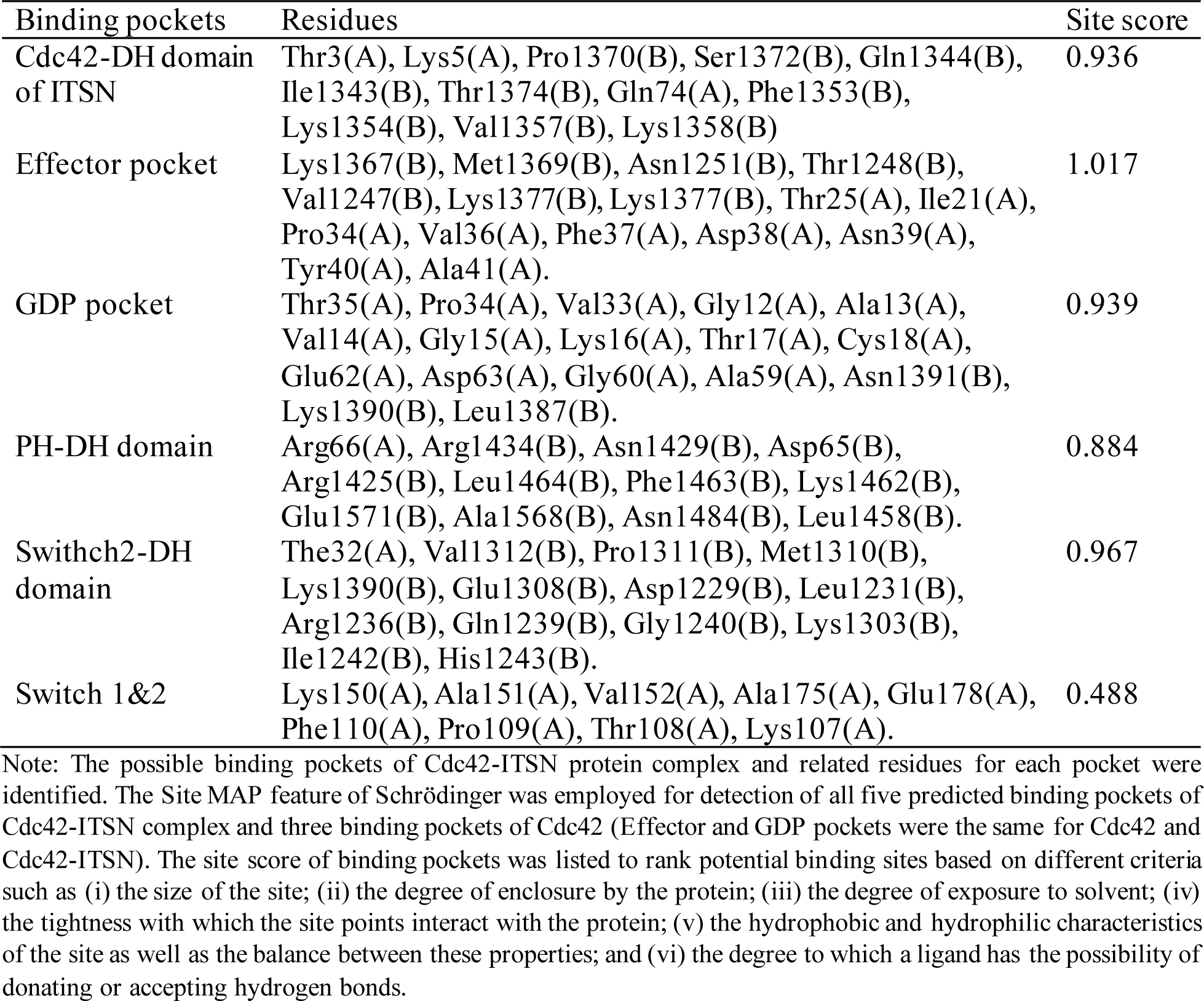
Site scores and residues of binding pockets in the interface of Cdc42-ITSN.

**Supplementary Table S2.**
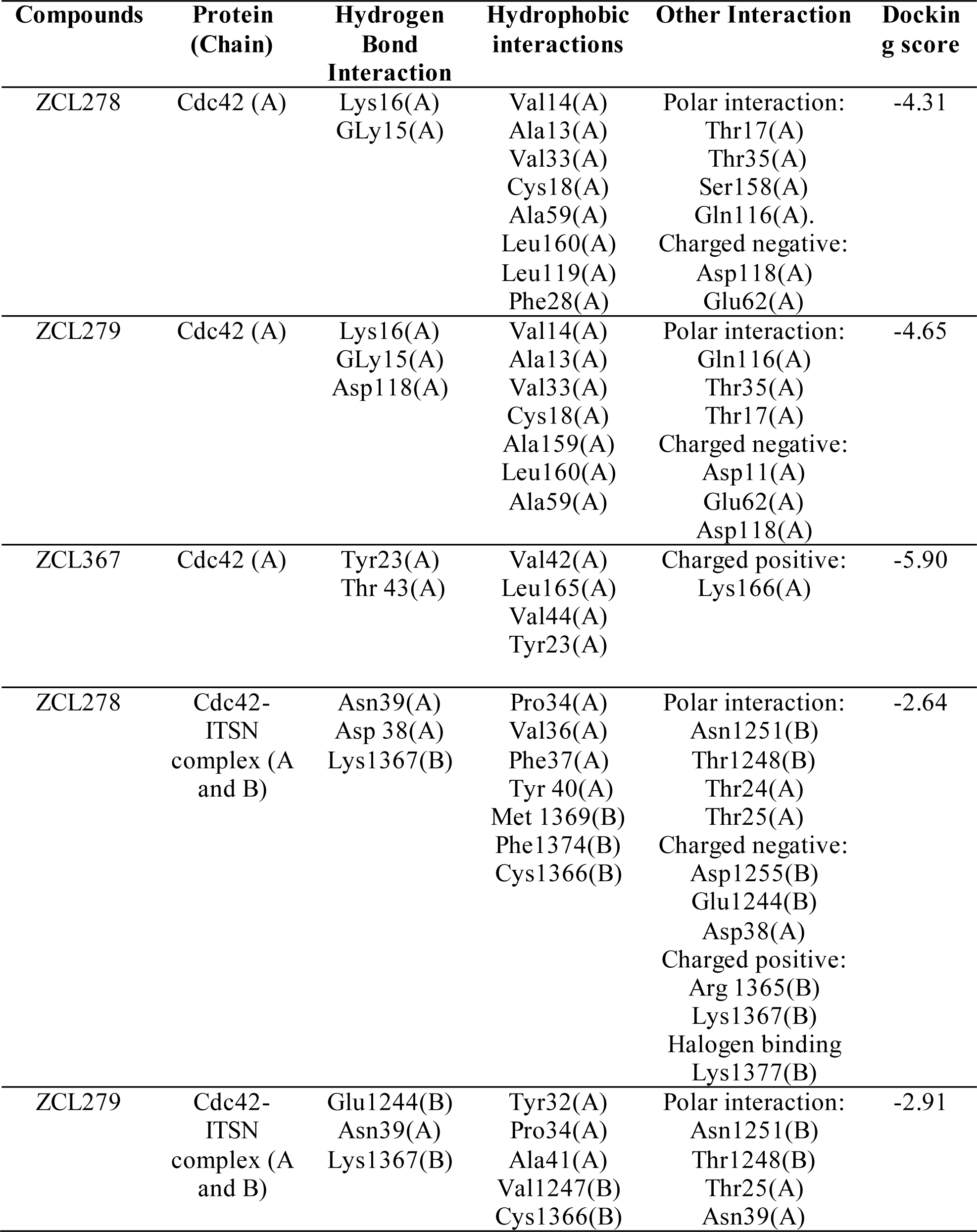
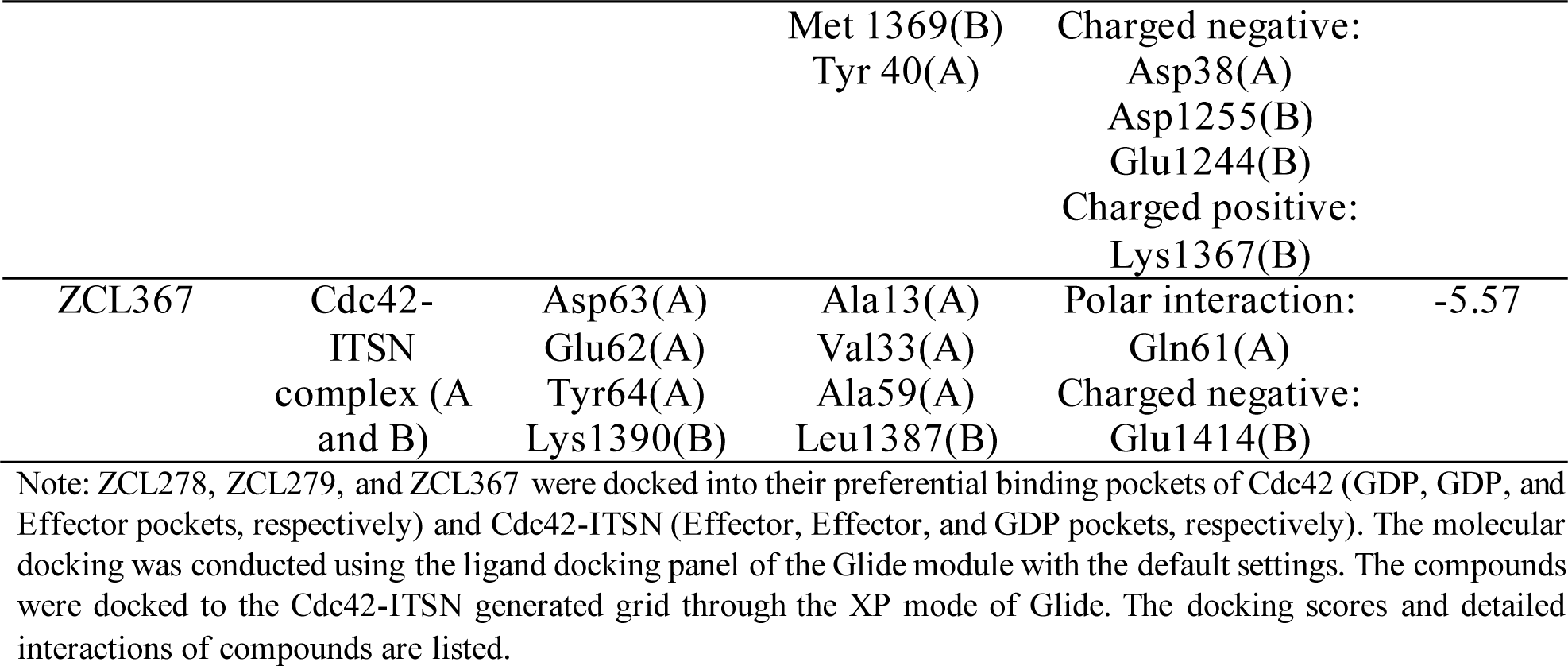
Interactions of ZCL278, ZCL279, and ZCL367 with Cdc42 and Cdc42-ITSN complex.

**Supplementary Table S3.**
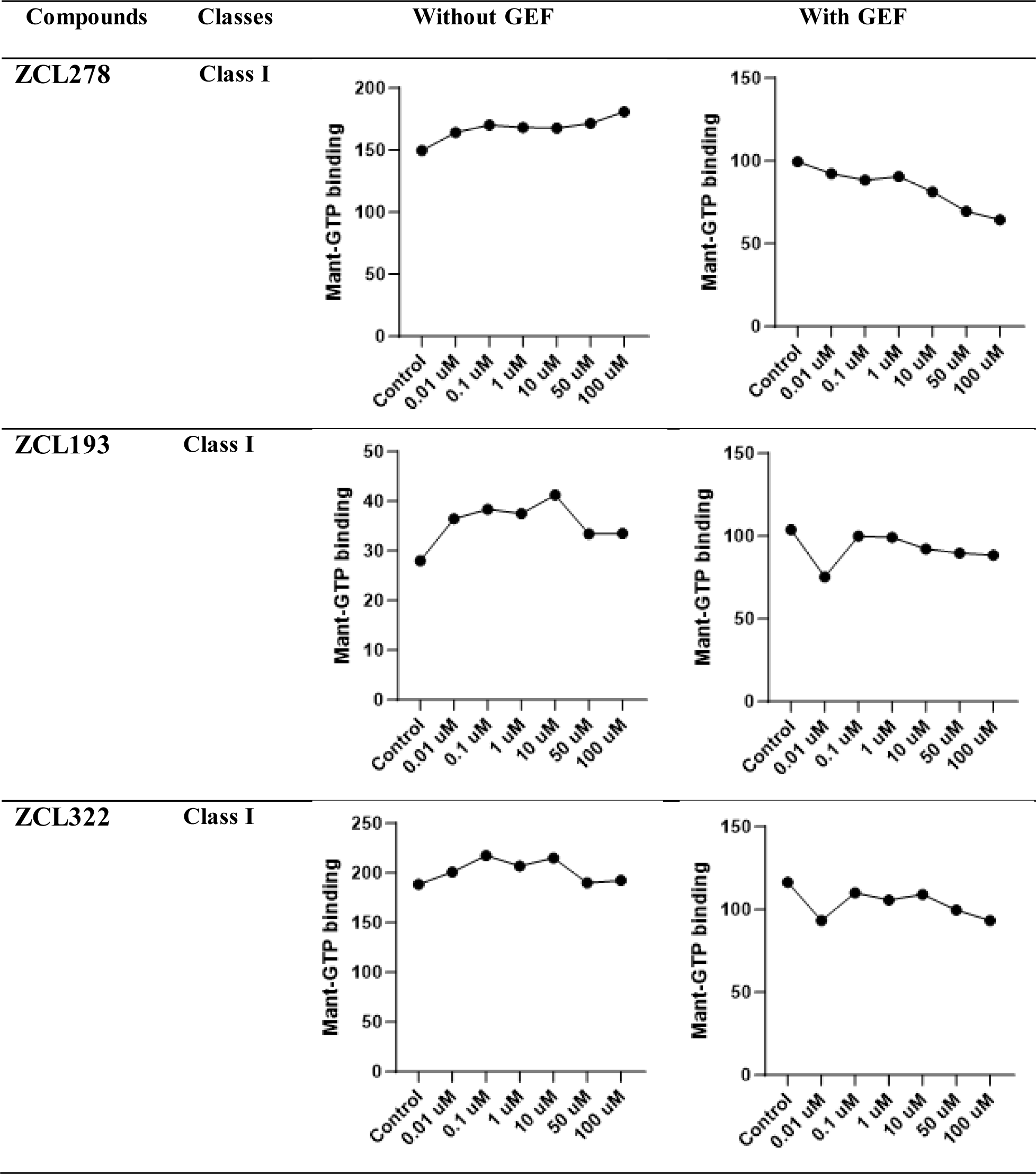
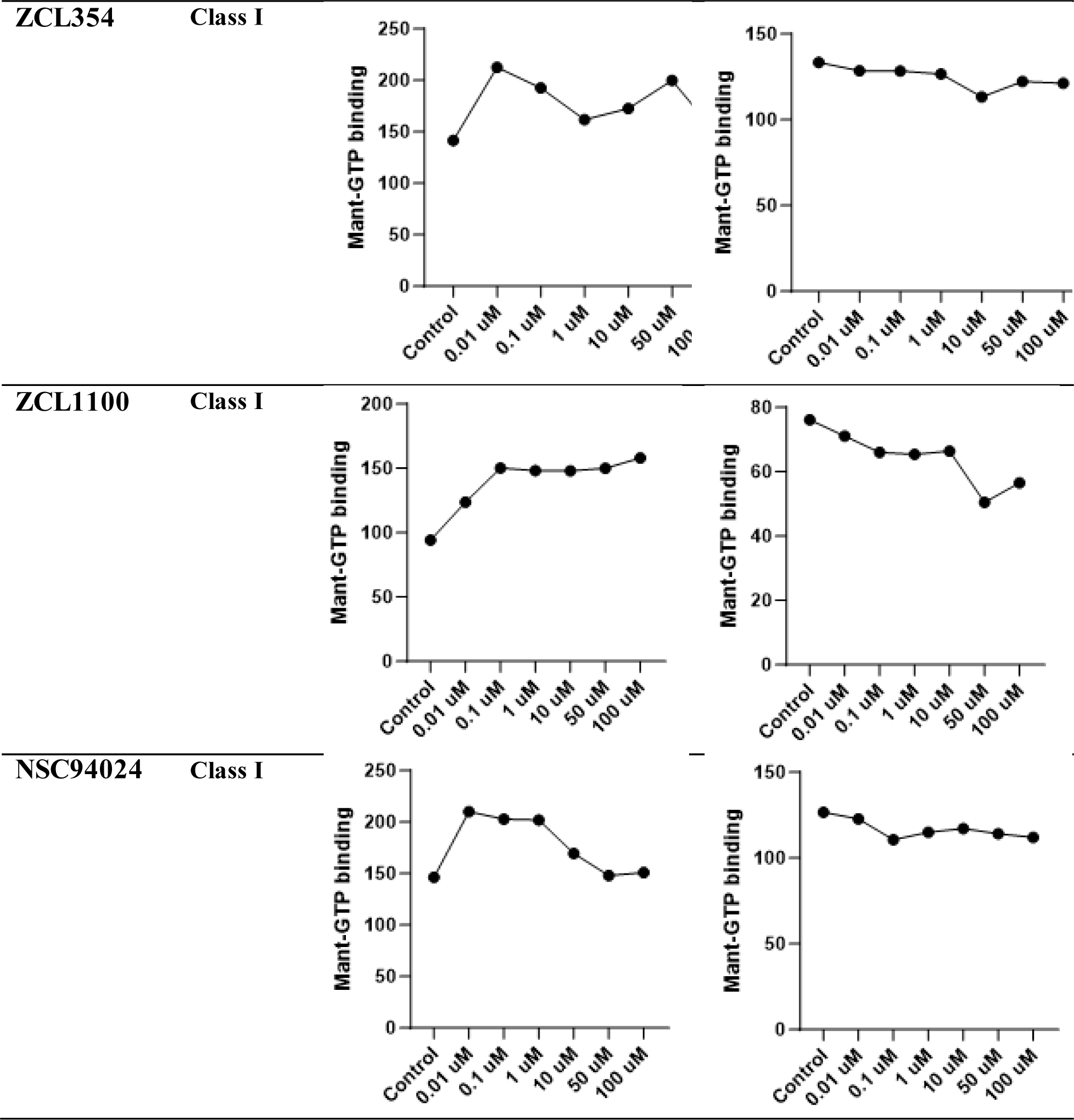
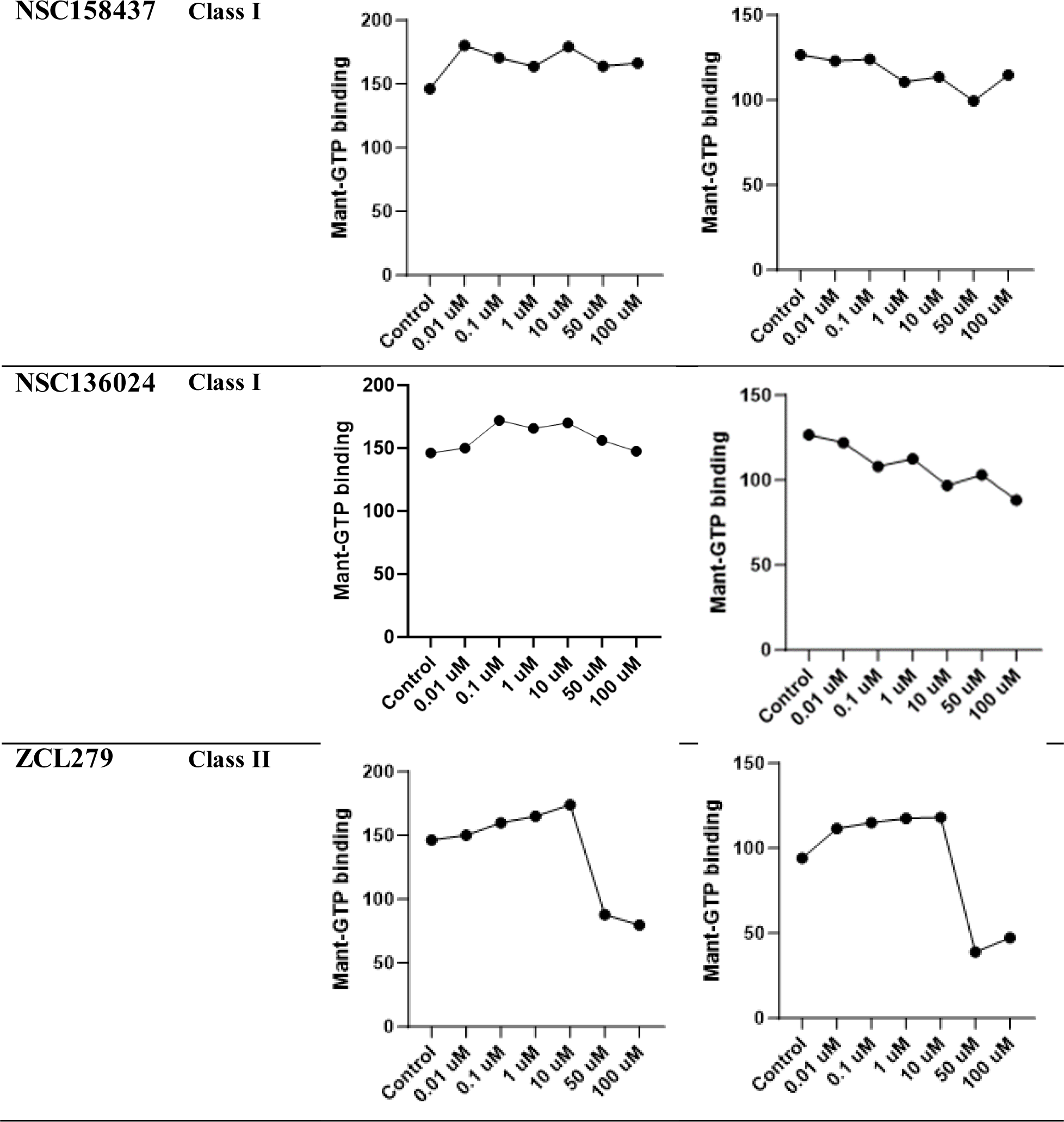
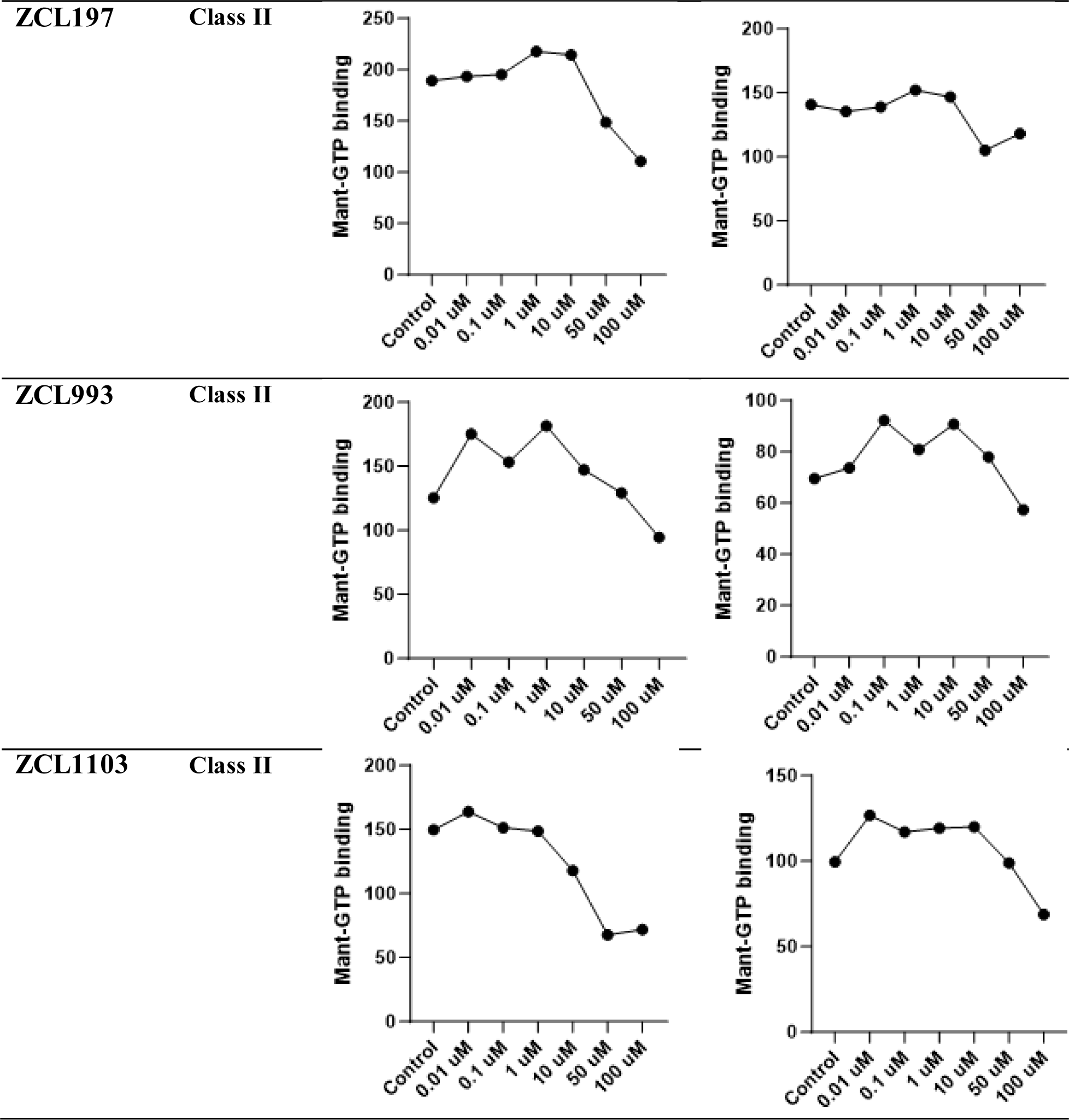
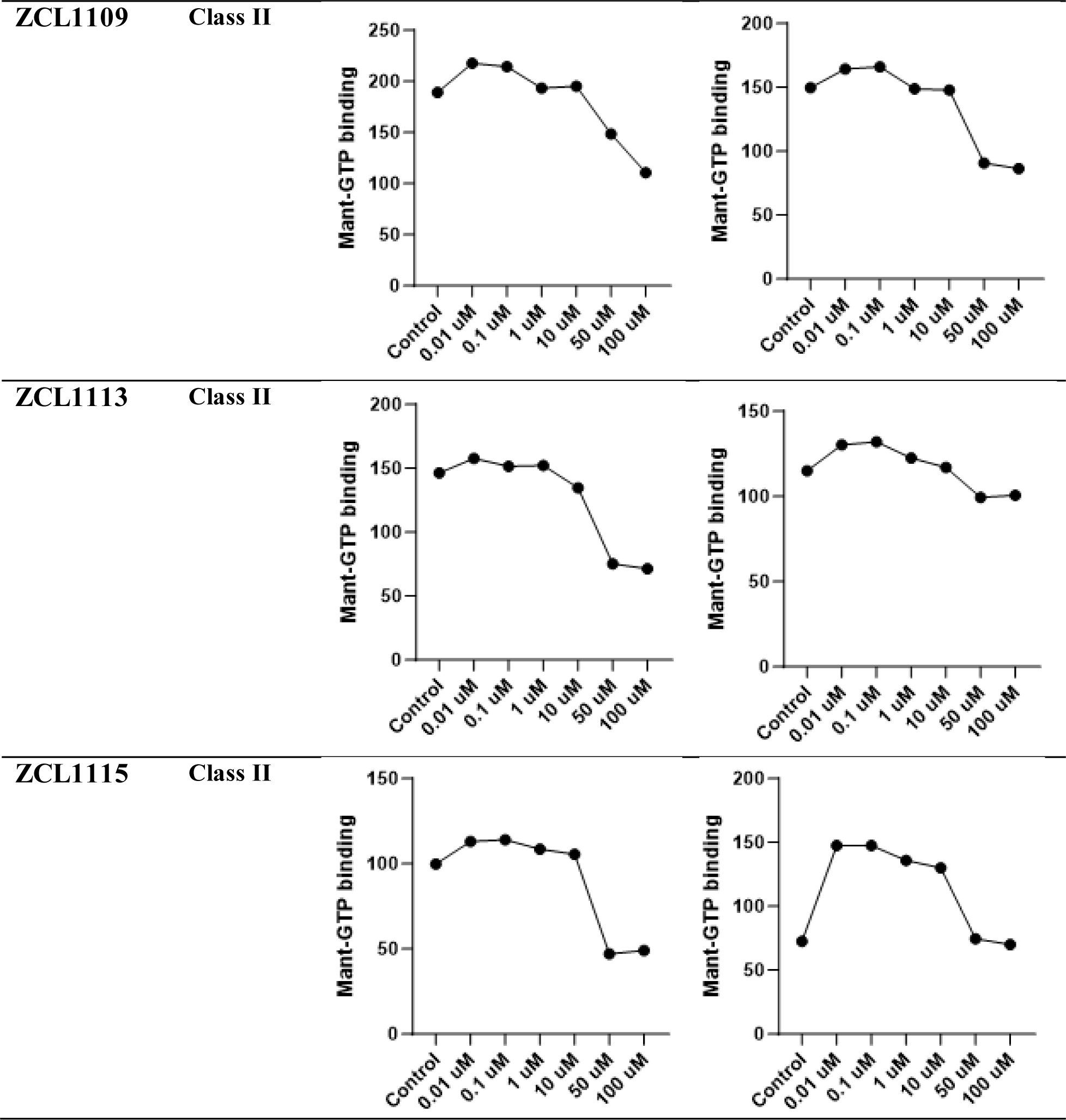
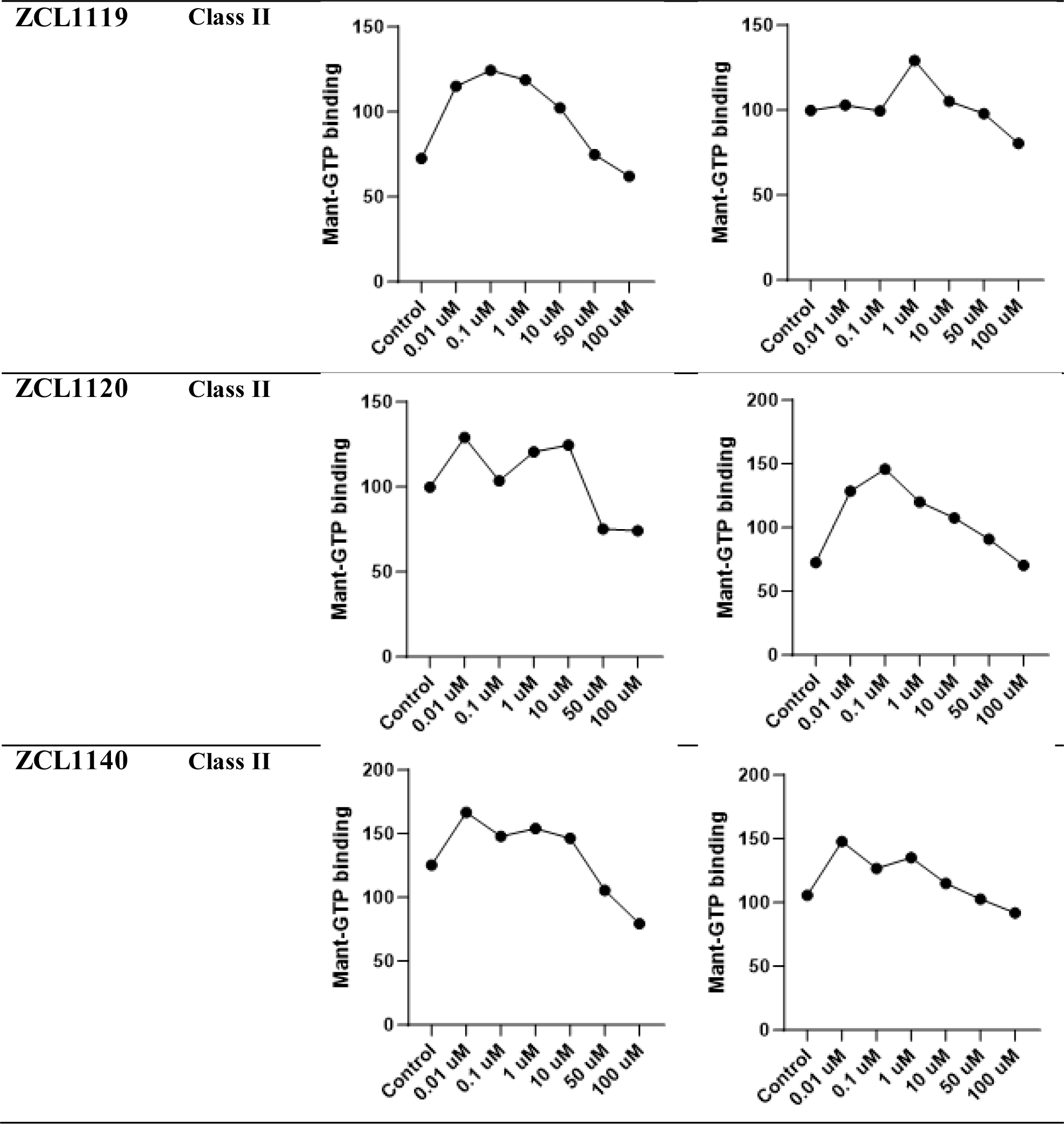
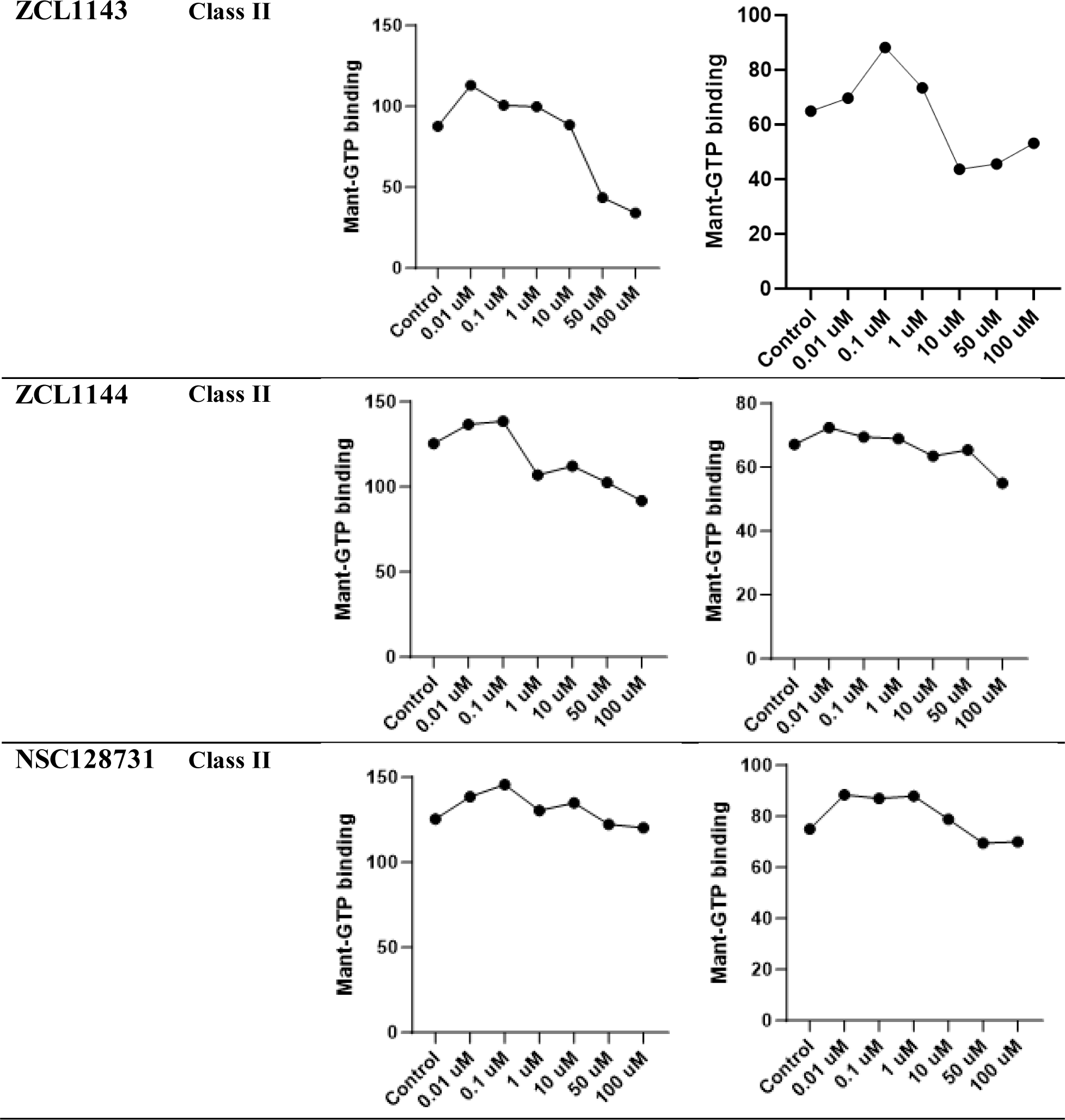
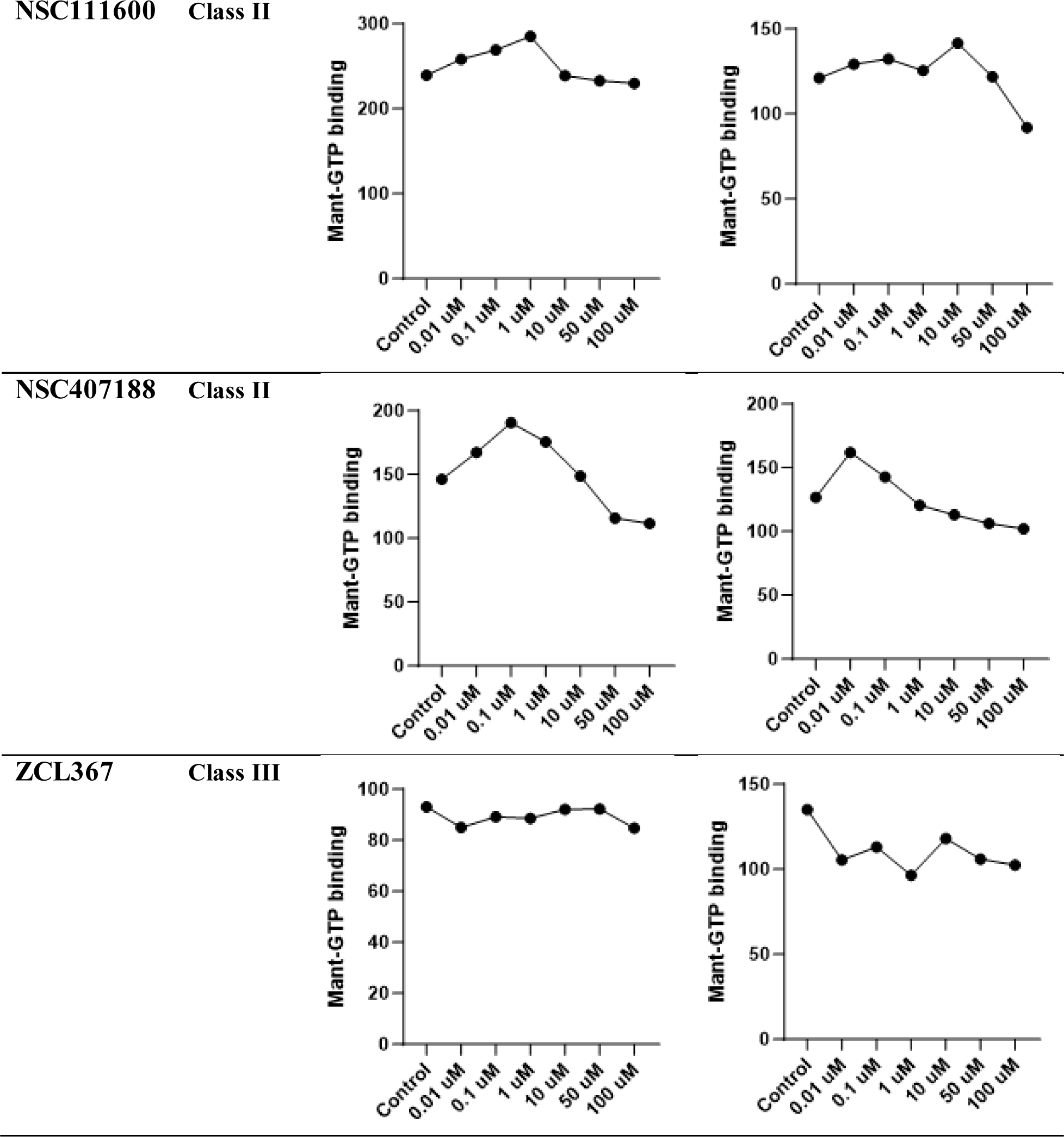
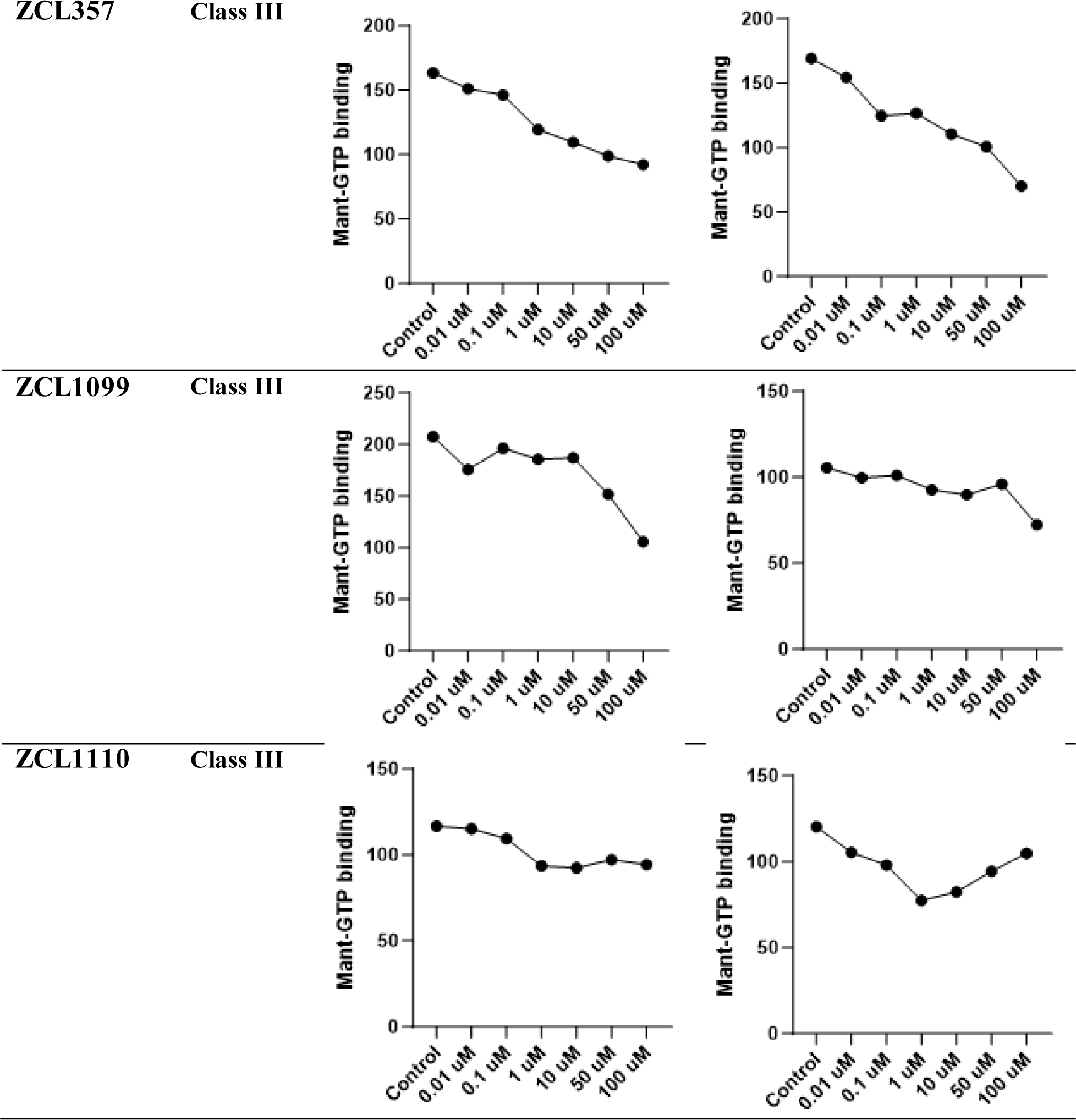
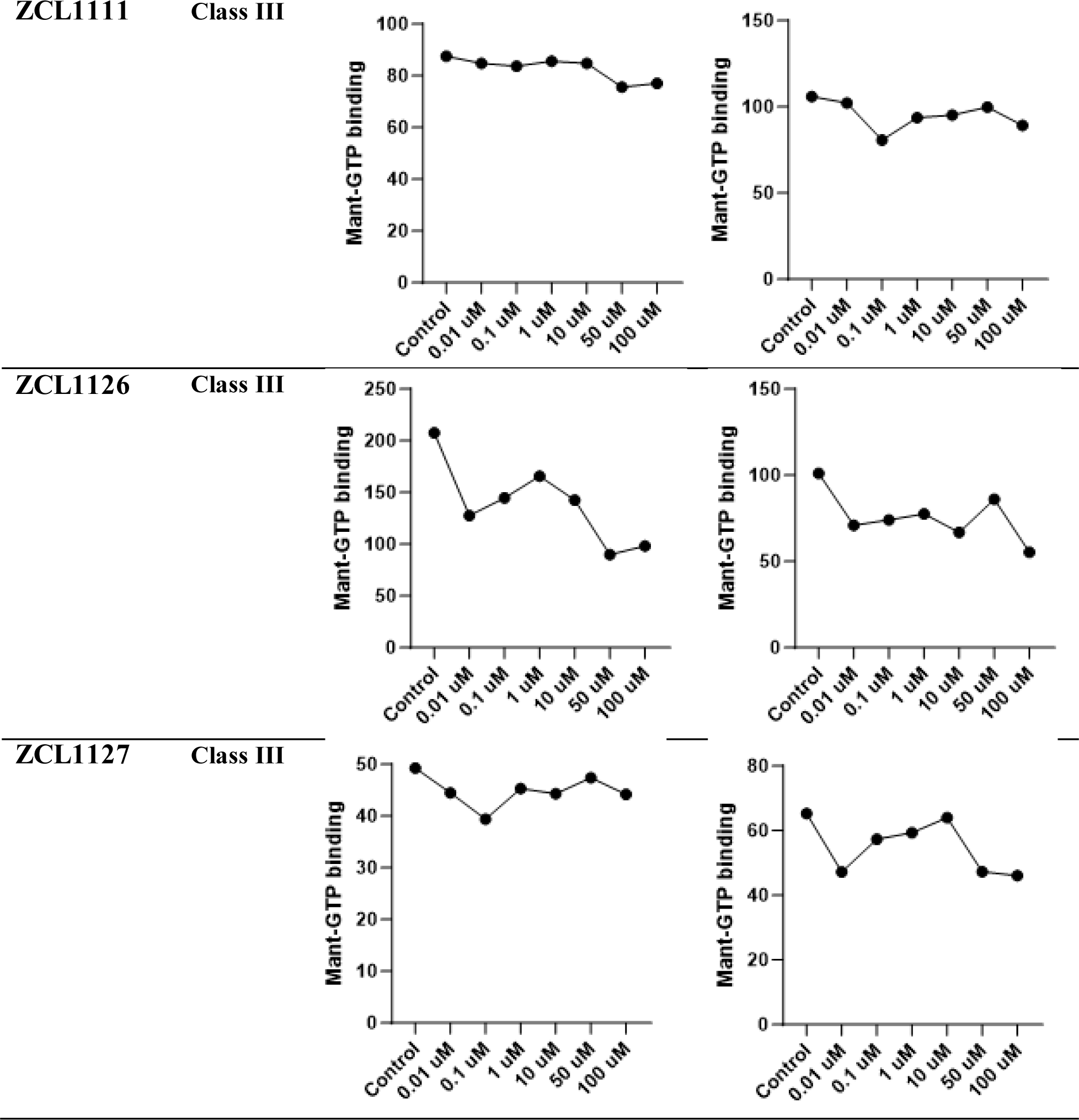
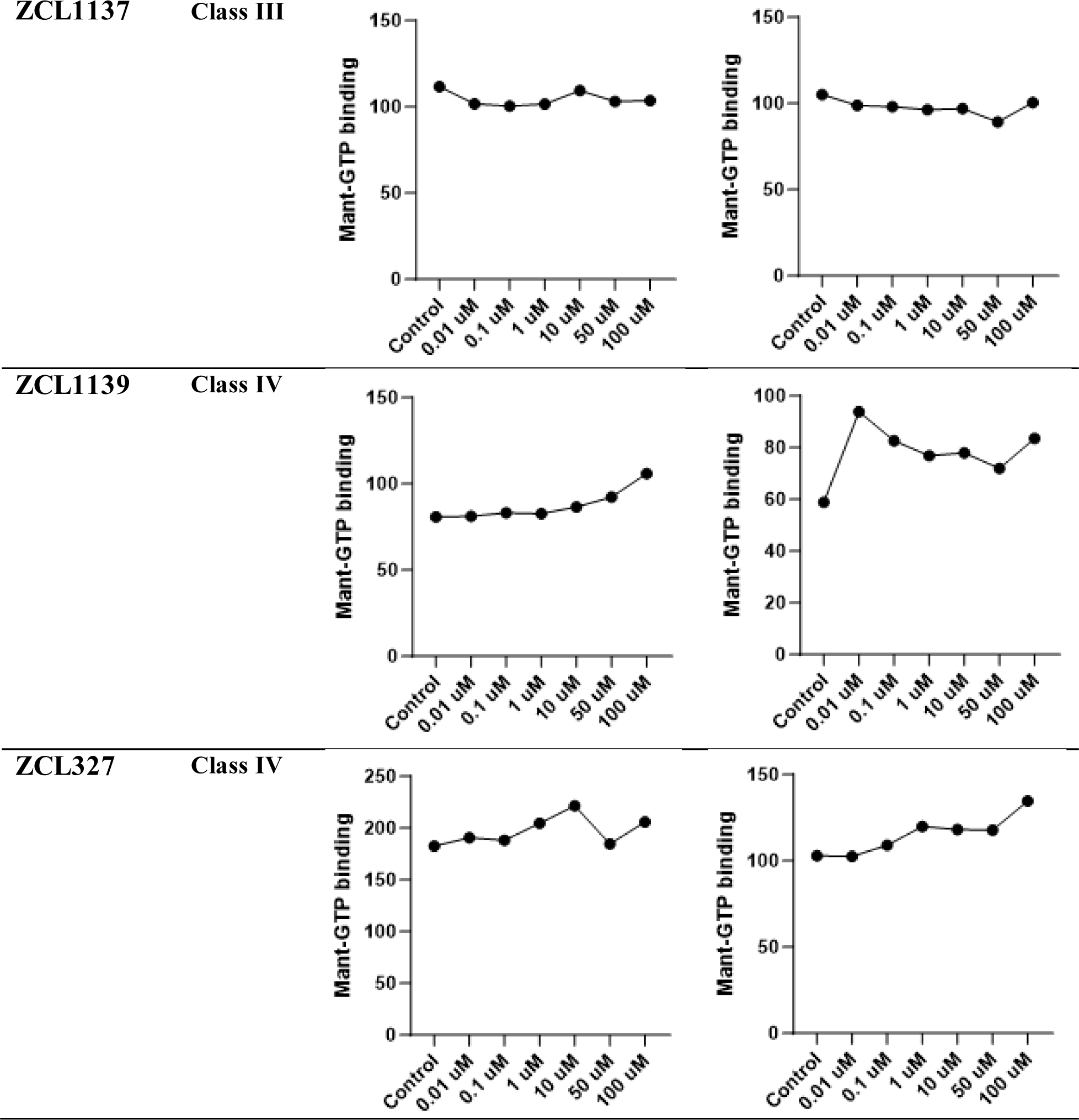
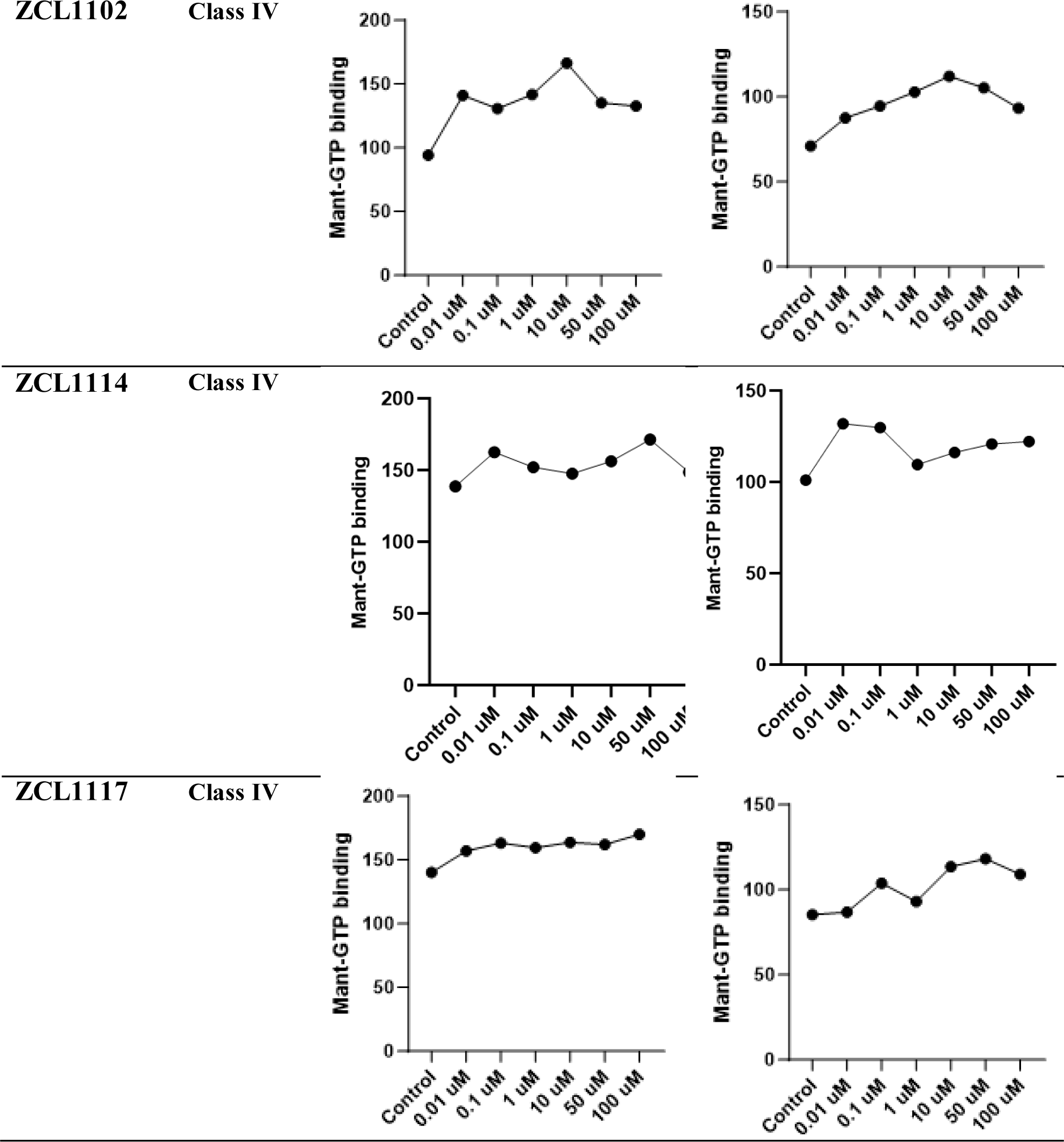
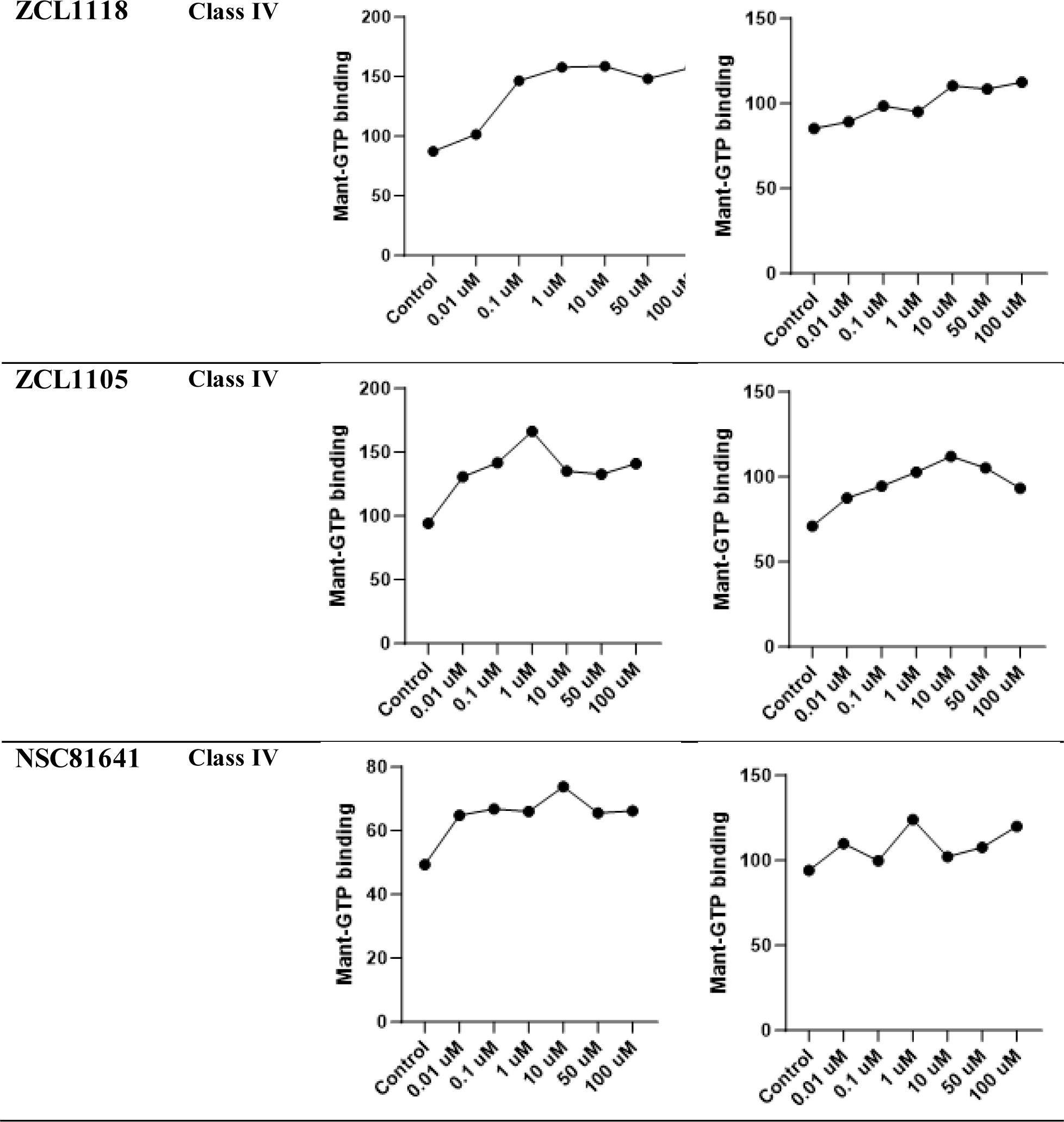
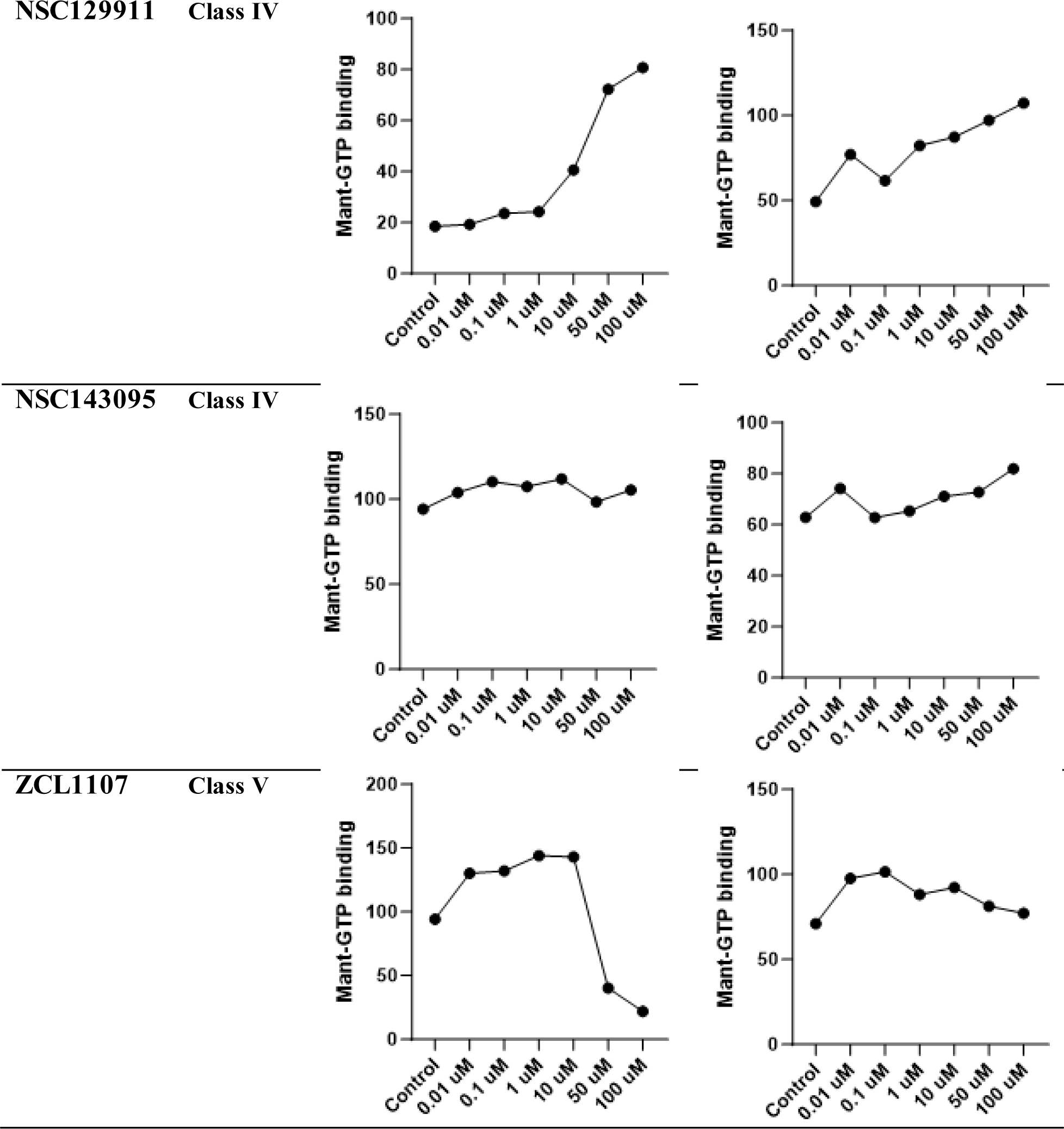
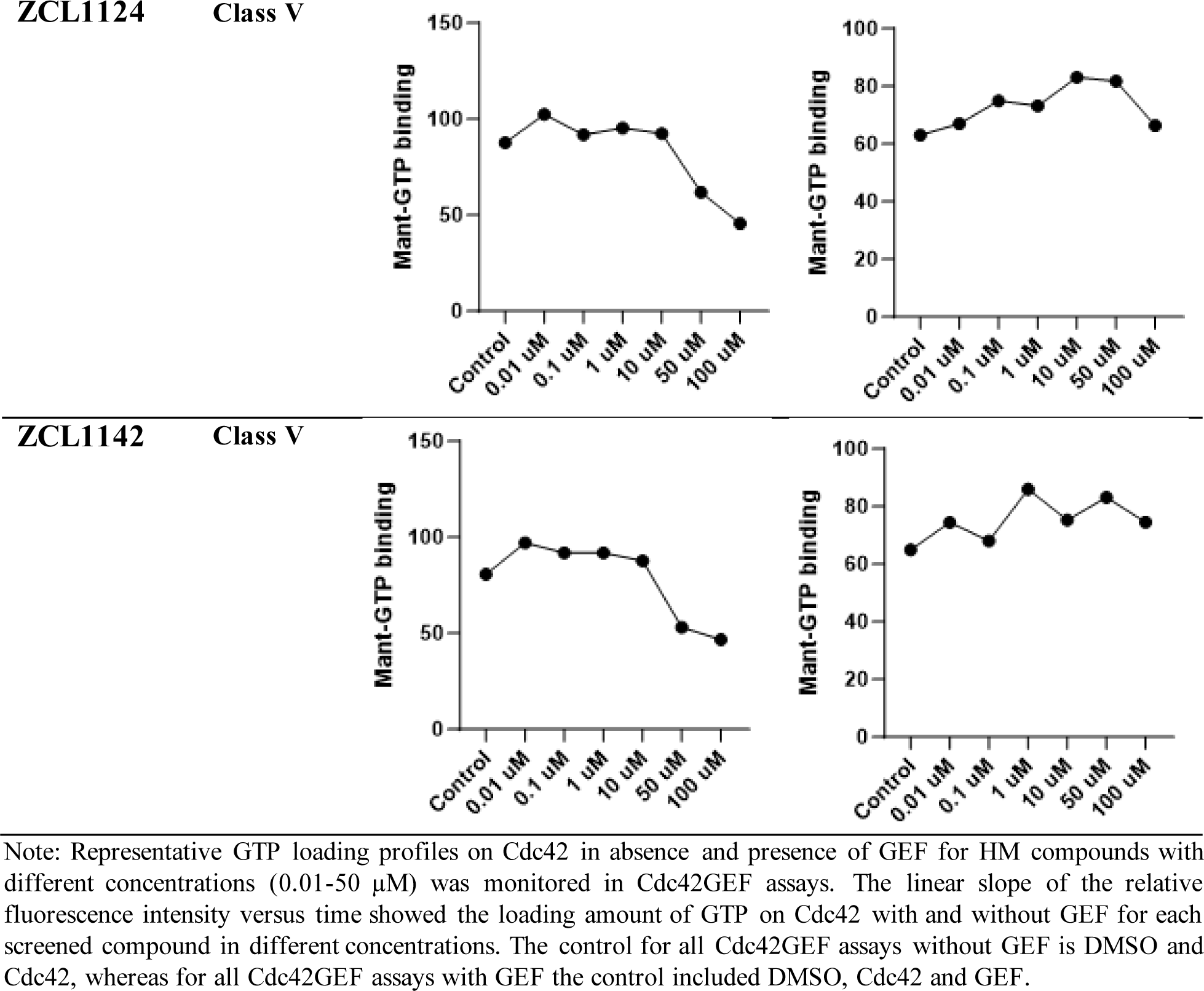
Biochemical evaluation of the five model compounds and new compounds screened from the HM libraries for GTP loading on Cdc42 with or without GEF.

**Supplementary Table S4.** Classification of HM library compounds into five model classes based on their interaction with Cdc42-ITSN complex by different screening methods.

| Screening Methods | ZCL278 | ZCL279 | ZCL367 | ZCL1139 | ZCL1107 |
| --- | --- | --- | --- | --- | --- |
| Molecular modeling (Screen score) <sup>a</sup> | ZCL279(1.43) | ZCL1127(1.55) | ZCL1142 (1.75) | ZCL354(1.71) | ZCL367(2.22) |
|  | ZCL1126(1.42) | NSC407188(1.46) | ZCL1102(1.71) | NSC94024(1.56) | NSC407188(2.21) |
|  | ZCL1103(1.38) | NSC94024(1.40) | ZCL1105 (1.60) | ZCL1105(1.55) | ZCL1117(2.19) |
|  | ZCL1127(1.36) | ZCL993 (1.39) | ZCL1127 (1.58) | NSC407188(1.55) | ZCL1142(2.09) |
|  | ZCL1099(1.29) | ZCL1143 (1.34) | ZCL1111 (1.58) | ZCL1139(1.55) | ZCL1118(2.01) |
|  | ZCL993(1.22) | NSC143095 (1.34) | ZCL1120 (1.51) | ZCL193(1.54) | ZCL193(1.99) |
|  | ZCL197(1.22) | NSC111600 (1.26) | ZCL1124 (1.49) | ZCL357(1.45) | ZCL279(1.85) |
|  | ZCL1105(1.12) | ZCL197 (1.24) | ZCL1117 (1.42) | NSC128731(1.42) | NSC94024(1.85) |
|  | ZCL1117(1.10) | ZCL367 (1.189) | ZCL993 (1.42) | NSC143095(1.35) | NSC136024(1.83) |
|  | ZCL1142(0.97) | ZCL193 (0.97) | ZCL1103(1.38) | NSC158437(1.33) | ZCL354(1.83) |
|  |  | ZCL354(0.54) | ZCL193(1.36) | ZCL1120(1.21) | NSC143095(1.81) |
|  |  |  | NSC158436(0.95) | ZCL1115(1.17) | NSC158437(1.73) |
|  |  |  |  | ZCL1103(1.10) | ZCL1109(1.72) |
|  |  |  |  | ZCL327(1.04) | ZCL327(1.68) |
|  |  |  |  | ZCL1299(1.03) | ZCL1111(1.67) |
|  |  |  |  | ZCL1111(0.99) | ZCL1115(1.67) |
|  |  |  |  | ZCL367(0.95) | ZCL1100(1.67) |
|  |  |  |  | ZCL279(0.94) | ZCL1105(1.65) |
|  |  |  |  | ZCL1137(0.94) | ZCL1124(1.57) |
|  |  |  |  |  | ZCL1144(1.54) |
|  |  |  |  |  | ZCL1299(1.50) |
|  |  |  |  |  | ZCL1103(1.48) |
| Shape screening (Matched score) <sup>b</sup> | ZCL197(0.62) | ZCL367(0.41) | ZCL1102(0.46) | ZCL1137(0.41) | ZCL1118(0.54) |
|  | ZCL993(0.4) | ZCL193(0.37) | ZCL1120(0.41) | ZCL357(0.41) | ZCL1119(0.52) |
|  | NSC158436(0.35) | ZCL1127(0.36) | NSC94024(0.39) | ZCL1119(0.39) | ZCL1111(0.41) |
|  | ZCL1127(0.34) | ZCL354(0.33) | ZCL1139(0.38) | ZCL1124(0.37) | ZCL1113(0.41) |
|  | ZCL1126(0.34) | ZCL993(0.33) | ZCL1107(0.38) | ZCL1120(0.36) | ZCL197(0.39) |
|  | ZCL1107(0.33) | NS158437(0.32) | NSC128731(0.37) | ZCL1103(0.36) | ZCL1102(0.39) |
|  | NSC136024(0.33) | ZCL1140(0.32) | ZCL1142(0.37) | ZCL1115(0.35) | ZCL1137(0.37) |
|  | NSC94024(0.33) | ZCL1142(0.31) | ZCL1137(0.37) | ZCL1109(0.35) | NSC94024(0.36) |
|  | ZCL1102(0.33) |  |  | ZCL367(0.35) | NSC136024(0.54) |
|  | ZCL1114(0.33) | ZCL1110(0.31) | ZCL1140(0.36) | ZCL1127(0.34) | ZCL1144(0.36) |
|  | NSC128731(0.33) | NSC143095(0.31) | ZCL1105(0.36) | ZCL1118(0.33) | ZCL1127(0.35) |
|  | ZCL1113(0.32) | ZCL197(0.31) | ZCL1126(0.35) | NSC111600(0.33) | ZCL1109(0.35) |
|  | ZCL193(0.30) | NSC94024(0.3) | ZCL1103(0.34) | NSC158437(0.32) | NSC128730(0.35) |
|  | ZCL354(0.3) | ZCL1117(0.3) | ZCL1111(0.34) | ZCL354(0.32) | NSC111600(0.34) |
|  | ZCL1105(0.3) | ZCL1100(0.3) | ZCL1114 (0.24) | ZCL1140(0.32) | ZCL1115(0.34) |
|  | NSC143095(0.3) | ZCL1111(0.3) |  | ZCL1126(0.31) | ZCL1142(0.34) |
|  | ZCL1144(0.27) | ZCL1126(0.3) |  | ZCL1099(0.31) | ZCL1105(0.34) |
|  | ZCL1119(0.27) | ZCL1099(0.3) |  | ZCL1117(0.31) | ZCL1140(0.33) |
|  | ZCL1118(0.26) | ZCL1102(0.29) |  | NSC143095(0.31) | NSC143095(0.33) |
|  | ZCL1120 (0.25) | ZCL1137(0.29) |  | NSC136024(0.30) | ZCL278(0.32) |
|  | ZCL1100(0.25) | ZCL1103(0.29) |  | NSC94024(0.3) | ZCL1117(0.32) |
|  | NSC111600(0.25) | NSC407188(0.28) |  | ZCL1105(0.29) |  |
|  | ZCL1140(0.25) | ZCL1120(0.28) |  | ZCL1144(0.29) |  |
|  | ZCL1111(0.24) | ZCL1144(0.27) |  | ZCL1114(0.29) |  |
|  |  | NSC111600(0.26) |  |  |  |
|  |  | ZCL1115(0.26) |  |  |  |
|  |  | ZCL1109(0.26) |  |  |  |
|  |  | ZCL1107(0.25) |  |  |  |
|  |  | ZCL1139(0.25) |  |  |  |
|  |  | NSC136024(0.25) |  |  |  |
|  |  | ZCL357(0.25) |  |  |  |
|  |  | ZCL1113(0.25) |  |  |  |
|  |  | ZCL1118(0.25) |  |  |  |
|  |  | ZCL1119(0.24) |  |  |  |
|  |  | NSC128731(0.24) |  |  |  |
|  |  | ZCL1114(0.24) |  |  |  |
| GEF assay | ZCL278 | ZCL279 | ZCL367 | ZCL327 | ZCL1124 |
| screening <sup>c</sup> | ZCL193 | ZCL197 | ZCL357 | ZCL1102 | ZCL1107 |
|  | ZCL322 | ZCL993 | ZCL1099 | ZCL1117 | ZCL1142 |
|  | ZCL354 | ZCL1103 | ZCL1110 | ZCL1118 |  |
|  | ZCL1100 | ZCL1109 | ZCL1111 | ZCL1105 |  |
|  | NSC94024 | ZCL1113 | ZCL1126 | ZCL1139 |  |
|  | NSC136024 | ZCL1115 | ZCL1127 | NSC81641 |  |
|  | NSC158437 | ZCL1119 | ZCL1137 | NSC129911 |  |
|  |  | ZCL1120 |  | NSC143095 |  |
|  |  | ZCL1140 |  | ZCL1114 |  |
|  |  | ZCL1144 |  |  |  |
|  |  | ZCL1143 |  |  |  |
|  |  | NSC128731 |  |  |  |
|  |  | NSC111600, |  |  |  |
|  |  | NSC407188, |  |  |  |
| Molecular | ZCL1137 | ZCL1137 | ZCL1144 | ZCL1144 | ZCL1107 |
| docking | ZCL1111 | ZCL1111 | NSC158437 | NSC158437 | ZCL1127 |
| preferential | ZCL1140 | ZCL1140 | ZCL193 | ZCL193 | ZCL1118 |
| binding | NSC143095 | NSC143095 | ZCL354 | ZCL354 | ZCL197 |
| pockets <sup>d</sup> | ZCL1119 | ZCL1119 | NSC136024 | NSC136024 | NSC94024 |
|  | ZCL1120 | ZCL1120 | ZCL1142 | ZCL1142 | NSC81641 |
|  | ZCL1117 | ZCL1117 | ZCL1109 | ZCL1109 | ZCL1114 |
|  | ZCL1103 | ZCL1103 | ZCL1099 | ZCL1099 |  |
|  | ZCL322 | ZCL322 | ZCL367 | ZCL367 |  |
|  | ZCL1113 | ZCL1113 | NSC128731 | NSC128731 |  |
|  | ZCL993 | ZCL993 | NSC111600 | NSC111600 |  |
|  | ZCL1124 | ZCL1124 | ZCL1110 | ZCL1110 |  |
|  | ZCL1126 | ZCL1126 | ZCL1105 | ZCL1105 |  |
|  | ZCL1102 | ZCL1102 | NSC407188 | NSC407188 |  |
|  | ZCL278 | ZCL278 | ZCL1139 | ZCL1139 |  |
|  | ZCL279. | ZCL279. | ZCL1114 | ZCL1114 |  |

**Supplementary Table S5.**
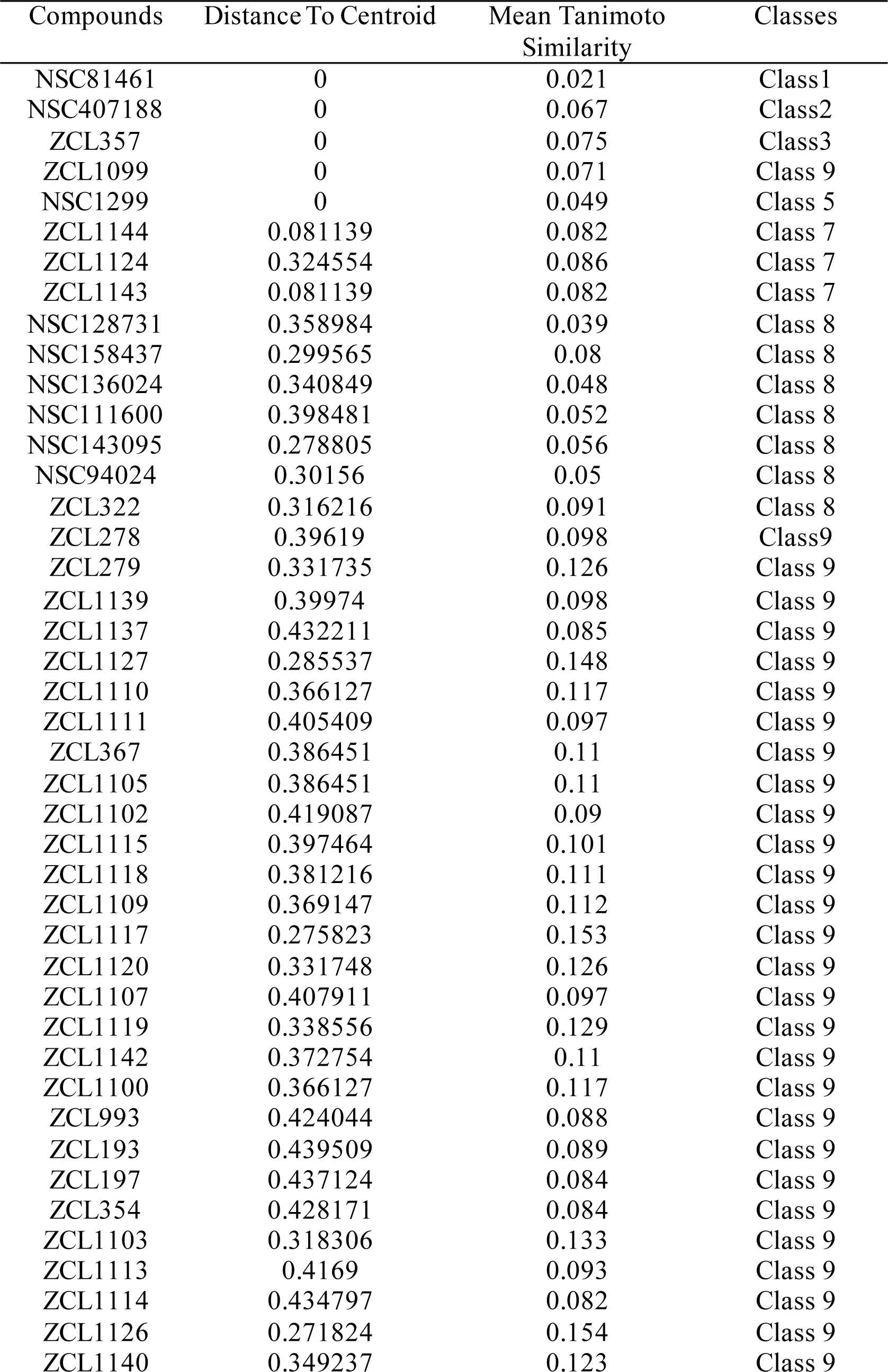
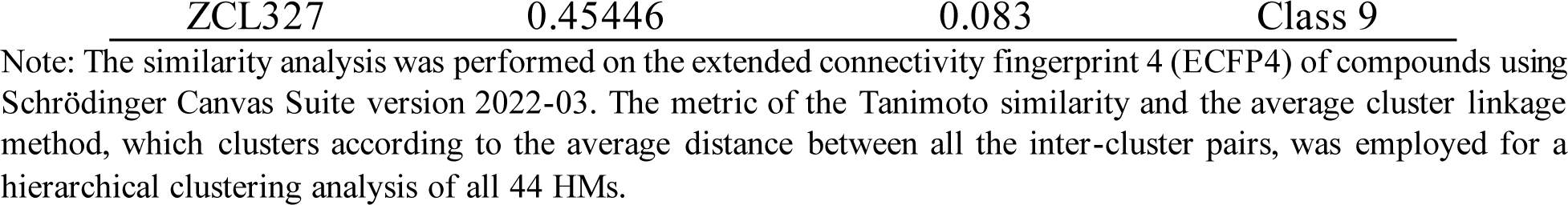
Summary of HM classification by hierarchical clustering.

| Compounds | Distance To Centroid | Mean Tanimoto<br>Similarity | Classes |
| --- | --- | --- | --- |
| NSC81461 | 0 | 0.021 | Class1 |
| NSC407188 | 0 | 0.067 | Class2 |
| ZCL357 | 0 | 0.075 | Class3 |
| ZCL1099 | 0 | 0.071 | Class 9 |
| NSC1299 | 0 | 0.049 | Class 5 |
| ZCL1144 | 0.081139 | 0.082 | Class 7 |
| ZCL1124 | 0.324554 | 0.086 | Class 7 |
| ZCL1143 | 0.081139 | 0.082 | Class 7 |
| NSC128731 | 0.358984 | 0.039 | Class 8 |
| NSC158437 | 0.299565 | 0.08 | Class 8 |
| NSC136024 | 0.340849 | 0.048 | Class 8 |
| NSC111600 | 0.398481 | 0.052 | Class 8 |
| NSC143095 | 0.278805 | 0.056 | Class 8 |
| NSC94024 | 0.30156 | 0.05 | Class 8 |
| ZCL322 | 0.316216 | 0.091 | Class 8 |
| ZCL278 | 0.39619 | 0.098 | Class9 |
| ZCL279 | 0.331735 | 0.126 | Class 9 |
| ZCL1139 | 0.39974 | 0.098 | Class 9 |
| ZCL1137 | 0.432211 | 0.085 | Class 9 |
| ZCL1127 | 0.285537 | 0.148 | Class 9 |
| ZCL1110 | 0.366127 | 0.117 | Class 9 |
| ZCL1111 | 0.405409 | 0.097 | Class 9 |
| ZCL367 | 0.386451 | 0.11 | Class 9 |
| ZCL1105 | 0.386451 | 0.11 | Class 9 |
| ZCL1102 | 0.419087 | 0.09 | Class 9 |
| ZCL1115 | 0.397464 | 0.101 | Class 9 |
| ZCL1118 | 0.381216 | 0.111 | Class 9 |
| ZCL1109 | 0.369147 | 0.112 | Class 9 |
| ZCL1117 | 0.275823 | 0.153 | Class 9 |
| ZCL1120 | 0.331748 | 0.126 | Class 9 |
| ZCL1107 | 0.407911 | 0.097 | Class 9 |
| ZCL1119 | 0.338556 | 0.129 | Class 9 |
| ZCL1142 | 0.372754 | 0.11 | Class 9 |
| ZCL1100 | 0.366127 | 0.117 | Class 9 |
| ZCL993 | 0.424044 | 0.088 | Class 9 |
| ZCL193 | 0.439509 | 0.089 | Class 9 |
| ZCL197 | 0.437124 | 0.084 | Class 9 |
| ZCL354 | 0.428171 | 0.084 | Class 9 |
| ZCL1103 | 0.318306 | 0.133 | Class 9 |
| ZCL1113 | 0.4169 | 0.093 | Class 9 |
| ZCL1114 | 0.434797 | 0.082 | Class 9 |
| ZCL1126 | 0.271824 | 0.154 | Class 9 |
| ZCL1140 | 0.349237 | 0.123 | Class 9 |

Note: The similarity analysis was performed on the extended connectivity fingerprint 4 (ECFP4) of compounds using Schrödinger Canvas Suite version 2022-03. The metric of the Tanimoto similarity and the average cluster linkage method, which clusters according to the average distance between all the inter-cluster pairs, was employed for a hierarchical clustering analysis of all 44 HMs.

**Supplementary Table S6.** Classification of HM library compounds into five model classes based on their interaction with Cdc42 without ITSN using different screening methods.

| Screening Methods | ZCL278 | ZCL279 | ZCL367 | ZCL1139 | ZCL1107 |
| --- | --- | --- | --- | --- | --- |
| Molecular modeling | ZCL197(1.38) | ZCL1142(1.24) | ZCL1124(1.92) | ZCL354(1.63) | ZCL367(2.30) |
|  | ZCL297(1.05) | ZCL1099(1.10) | ZCL327(1.80) | ZCL1111(1.53) | ZCL1111(2.30) |
|  |  | ZCL993(1.10) | ZCL1105(1.63) | ZCL193(1.42) | ZCL327(2.22) |
|  |  | ZCL1103(1.01) | ZCL279(1.63) | ZCL1137(1.42) | ZCL1142(2.18) |
|  |  | ZCL1126(0.94) | ZCL1102(1.61) | ZCL1142(1.40) | ZCL1105(2.14) |
|  |  | ZCL1299(0.76) | ZCL1127(1.59) | ZCL1115(1.31) | ZCL193(2.12) |
|  |  |  | ZCL193(1.58) | ZCL327(1.27) | ZCL322(2.10) |
|  |  |  | ZCL1109(1.57) | NSC129911(1.15) | NSC128730(1.99) |
|  |  |  | NSC143094(1.56) |  | NSC94024(1.95) |
|  |  |  | NSC94024(1.53) |  | ZCL1118(1.94) |
|  |  |  | ZCL1111(1.49) |  | ZCL279(1.90) |
|  |  |  | ZCL354(1.47) |  | NSC143095(1.88) |
|  |  |  | ZCL1117(1.24) |  | ZCL354(1.80) |
|  |  |  | ZCL357(1.17) |  | ZCL1137(1.78) |
|  |  |  | ZCL993(1.16) |  | NSC136024(1.75) |
|  |  |  | NSC158436(1.04) |  | NSC407188(1.74) |
|  |  |  | ZCL322 (1.04) |  | ZCL1119(1.70) |
|  |  |  | NSC128731(1.02) |  | ZCL357(1.63) |
|  |  |  | ZCL1126(1.01) |  | ZCL1100(1.62) |
|  |  |  | NSC129911(0.75). |  | ZCL1113(1.58) |
|  |  |  |  |  | ZCL1117(1.62) |
| Shape screening (Matched score)<sup>b</sup> | ZCL197(0.62) | ZCL367(0.41) | ZCL1102(0.46) | ZCL1137(0.41) | ZCL1118(0.54) |
|  | ZCL 993(0.42) | ZCL193(0.37) | ZCL1120(0.41) | ZCL357(0.41) | ZCL1119(0.52) |
|  | NSC158436(0.35) | ZCL1127(0.36) | NSC94024(0.39) | ZCL1119(0.39) | ZCL1111(0.41) |
|  | ZCL 1127(0.34) | ZCL354(0.33) | ZCL1139(0.38) | ZCL1124(0.37) | ZCL1113(0.41) |
|  | ZCL1126(0.34) | ZCL993(0.33) | ZCL1107(0.38) | ZCL1120(0.36) | ZCL197(0.39) |
|  | ZCL1107(0.33) | NSC158437(0.32) | NSC128731(0.37) | ZCL1103(0.36) | ZCL1102(0.39) |
|  | NSC136024(0.33) | ZCL1140(0.32) | ZCL1142(0.37) | ZCL1115(0.35) | ZCL1137(0.37) |
|  | NSC94024(0.33) | ZCL1142(0.31) | ZCL1137(0.37) | ZCL1109(0.35) | NSC94024(0.36) |
|  | ZCL1102(0.33) | ZCL1110(0.31) | ZCL1140(0.36) | ZCL367(0.35) | NSC136024(0.54) |
|  | ZCL1114(0.33) | NSC143095(0.31) | ZCL1105(0.36) | ZCL1127(0.34) |  |
|  | NSC128731(0.33) | ZCL197(0.31) | ZCL1126(0.35) | ZCL1118(0.33) | ZCL1144(0.36) |
|  | ZCL1113(0.32) | NSC94024(0.3) | ZCL1103(0.34) | NSC111600(0.33) | ZCL1127(0.35) |
|  | ZCL193(0.30) | ZCL1117(0.3) | ZCL1111(0.34) | NSC158437(0.32) | ZCL1109(0.35) |
|  | ZCL354(0.3) | ZCL1100(0.3) | ZCL1114 (0.24) | ZCL354(0.32) | NSC128730(0.35) |
|  | ZCL1105(0.3) | ZCL1111(0.3) |  | ZCL1140(0.32) | NSC111600(0.34) |
|  | NSC143095(0.3) | ZCL1126(0.3) |  | ZCL1126(0.31) | ZCL1115(0.34) |
|  | ZCL1144(0.27) | ZCL1099(0.3) |  | ZCL1099(0.31) | ZCL1142(0.34) |
|  | ZCL1119(0.27) | ZCL1102(0.29) |  | ZCL1117(0.31) | ZCL1105(0.34) |
|  | ZCL1118(0.26) | ZCL1137(0.29) |  | NSC143095(0.31) | ZCL1140(0.33) |
|  | ZCL1120 (0.25) | ZCL1103(0.29) |  | NSC136024(0.30) | NSC143095(0.33) |
|  | ZCL1100(0.25) | NSC407188(0.28) |  | NSC94024(0.3) | ZCL278(0.32) |
|  | NSC111600(0.25) | ZCL1120(0.28) |  | ZCL1105(0.29) | ZCL1117(0.32) |
|  | ZCL1140(0.25) | ZCL1144(0.27) |  | ZCL1144(0.29) |  |
|  | ZCL1111(0.24). | NSC111600(0.26) |  | ZCL1114(0.29) |  |
|  |  | ZCL1115(0.26) |  |  |  |
|  |  | ZCL1109(0.26) |  |  |  |
|  |  | ZCL1107(0.25) |  |  |  |
|  |  | ZCL1139(0.25) |  |  |  |
|  |  | NSC136024(0.25), |  |  |  |
|  |  | ZCL357(0.25) |  |  |  |
|  |  | ZCL1113(0.25) |  |  |  |
|  |  | ZCL1118(0.25), |  |  |  |
|  |  | ZCL1119(0.24) |  |  |  |
|  |  | NSC128731(0.24) |  |  |  |
|  |  | ZCL1114(0.24) |  |  |  |
| GEF assay | ZCL278 | ZCL279 | ZCL367 | ZCL327 | ZCL1124 |
| screening <sup>c</sup> | ZCL193 | ZCL197 | ZCL357 | ZCL1102 | ZCL1107 |
|  | ZCL322 | ZCL993 | ZCL1099 | ZCL1117 | ZCL1142 |
|  | ZCL354 | ZCL1103 | ZCL1110 | ZCL1118 |  |
|  | ZCL1100 | ZCL1109 | ZCL1111, | ZCL1105 |  |
|  | NSC94024 | ZCL1113 | ZCL1126 | ZCL1139 |  |
|  | NSC136024 | ZCL1115 | ZCL1127 | NSC81641 |  |
|  | NSC158437 | ZCL1119 | ZCL1137 | NSC129911 |  |
|  |  | ZCL1120 |  | NSC143095 |  |
|  |  | ZCL1140 |  | ZCL1114 |  |
|  |  | ZCL1144 |  |  |  |
|  |  | ZCL1143 |  |  |  |
|  |  | NSC128731 |  |  |  |
|  |  | NSC111600 |  |  |  |
|  |  | NSC407188 |  |  |  |
| Molecular | ZCL357 | ZCL1137 | ZCL1119 | ZCL1119 | ZCL1119 |
| docking | ZCL1099 | ZCL1102 | ZCL1117 | ZCL1117 | ZCL1117 |
| preferential | ZCL278 | ZCL1140 | ZCL1105 | ZCL1105 | ZCL1105 |
| binding |  | NSC143095 | ZCL1107 | ZCL1107 | ZCL1107 |
| pockets <sup>d</sup> |  | ZCL1113 | ZCL1120 | ZCL1120 | ZCL1120 |
|  |  | ZCL993 | ZCL1103 | ZCL1103 | ZCL1103 |
|  |  | ZCL1124 | NSC158437 | NSC158437 | NSC158437 |
|  |  | ZCL1144 | ZCL193 | ZCL193 | ZCL193 |
|  |  | ZCL358 | ZCL354 | ZCL354 | ZCL354 |
|  |  | ZCL1143 | ZCL1114 | ZCL1114 | ZCL1114 |
|  |  | NSC136024 | ZCL1142 | ZCL1142 | ZCL1142 |
|  |  | ZCL197 | ZCL1109 | ZCL1109 | ZCL1109 |
|  |  | ZCL1127 | NSC128721 | NSC128721 | NSC128721 |
|  |  | NSC94024 | NSC111600 | NSC111600 | NSC111600 |
|  |  | NSC81641 | ZCL1139 | ZCL1139 | ZCL1139 |
|  |  | ZCL279 | NSC407188 | NSC407188 | NSC407188 |
|  |  | ZCL1126 | ZCL1115 | ZCL1115 | ZCL1115 |
|  |  | ZCL351 | ZCL367 | ZCL367 | ZCL367 |
|  |  | ZCL1118 | ZCL322 | ZCL322 | ZCL322 |
|  |  | NSC129911 | ZCL1110 | ZCL1110 | ZCL1110 |
|  |  |  | ZCL1111 | ZCL1111 | ZCL1111 |

